# Epigenetic regulators partition the genome by evolutionary and developmental constraint

**DOI:** 10.64898/2026.10.01.755350

**Authors:** Yusuke Kijima, JoAnne Villagrana, Minming Wang, Rosaria Tornisiello, Nicolas Dias, Jason Tien-Chi Hou, Kevin Tse, Jennifer Wang, Helene Kretzmer, Kaelyn Sumigray, Tuğçe Aktaş, Zachary D Smith

## Abstract

Many epigenetic regulators play essential roles in animal development, but precisely how they function across different cell types without clear sequence preferences remains a foundational challenge. Here, we employed single-cell RNA sequencing to survey the mutant phenotypes of 30 essential regulatory pathways across over 450 gastrulation stage mouse embryo replicates, including essential factors associated with repressive chromatin states, RNA surveillance/processing, and other genomic roles. We then use this integrated manifold to partition mutant responses into a developmentally-resolved interaction network, revealing a hierarchy of heterochromatin-associated interactions that act orthogonally to Polycomb Repressive Complex-mediated control of embryo patterning. Compared to the embryo proper, we find that the developing placenta and yolk sac can accommodate broad swings in repetitive element expression, a resilience in keeping with the relative evolutionary recency and phenotypic plasticity of these tissues. Dense temporal sampling of Human Silencing Hub (HUSH) mutant embryogenesis further supports the notion that the gastrulating embryo is particularly constrained and depends on multiple layers of insulation, including the need to suppress evolutionarily young genetic elements to stably license highly conserved, phylotypic programs. In combination, our work highlights the power of single cell-resolved developmental genetics to contextualize the roles of epigenetic regulators, which restrict genomic action to support robust cellular identities and embryological forms.

## Introduction

The development of multicellular organisms requires precisely timed control of encoded gene regulatory networks, which guide cell fate decisions towards a reproducible body plan^1–3^. This concept has often been described as an “epigenetic landscape” and extended to include chemical modifications to chromatin, which act as molecular barriers (or ridges in the landscape) between developing lineages to constrain transcriptional activity towards appropriate outcomes^4,5^. The potential for chromatin state to instruct cell identity has inspired massive efforts to uncover its basic blueprints, including massive genomic atlases of diverse histone and DNA modification states from fertilization throughout the natural life span and across major diseases^6–9^. Combined with extensive genetic and biochemical characterization in cell lines, these works have inspired a model where many genomic regulators act by stabilizing differentiation and cell identity in the face of fluctuating environments^10–12^.

In mice, mutant phenotypes for many epigenetic regulators emerge shortly after embryo implantation, which is a period of accelerated growth, differentiation, and morphogenesis; however, why and how comparatively sequence-independent enzymes are so essential to the earliest stages of organismal development remains unclear^10^. Notably, this window also overlaps with a morphologically and molecularly conserved phylotypic period, in which germ layer formation and anterior-posterior patterning take place^13,14^, implying that epigenetic regulation tunes the activity of evolutionarily emergent and conserved programs. As evocative as these ideas have been, the complex and dynamic nature of early embryogenesis makes it difficult to connect what we know about a given regulator’s biochemistry or genomic occupancy to its precise developmental role^10^. This challenge is further complicated by the breadth of any given regulator’s described activities and interactions, including functions beyond gene expression such as retrotransposon suppression, parent-specific imprinting, or splicing fidelity, as well as chromosome-scale roles in centromere maintenance, X chromosome (ChrX) silencing in females, or nuclear topology^15–18^. Sophisticated in vitro assays and cell culture experiments have provided precise mechanistic details into how certain modification states are established or propagated, but how different classes of regulators coordinate dynamic developmental processes in vivo remains to be explored^19–22^.

The rise of single-cell genomics over the past decade has offered new opportunities to reinvestigate animal development in high resolution. Single-cell data can be used to embed embryogenesis as a continuous “manifold” that offers a holistic view of individual cellular and molecular states, including the opportunity to evaluate regulator function with temporal resolution^23–31^. Recent studies have leveraged this strength to characterize the roles of congenital mutations, transcription factors, and epigenetic pathways such as DNA methylation and oxidation^27,32–36^. However, as with more extensive work conducted in stem cells, previous studies have generally focused on only one or several related modification states, leaving the interconnected or context-specific roles of these multi-layered regulators unaddressed. Accordingly, cell type-resolved single-cell analysis of zygotic mutants would provide a unique opportunity to investigate how generic biochemical regulators may support conserved versus emergent developmental programs according to their own distinct evolutionary histories and selective pressures^37^.

To begin investigating the epigenetic control of mammalian development, we employed a CRISPR/Cas9-based “Crispant” strategy^33,35,38^ that enables the integrative analysis of multiple regulator classes across a densely-sampled manifold of mouse gastrulation and early organogenesis. We genetically perturbed 30 nuclear regulators involved in a broad range of genomic functions, including the assembly of distinct heterochromatin states, co-transcriptional regulation, and retrotransposon control, and performed single-cell RNA-seq to compare their mutant phenotypes across multiple scales (**Fig. 1a**). Using highly polymorphic sperm, we were able to robustly pinpoint changes in transcriptional behavior within replicate-powered mutant cohorts. We then used this framework to evaluate the impact of nuclear regulators on mouse embryogenesis, including effects on differentiation, ChrX silencing in females, endogenous gene and repetitive element activities, and the evolutionary dynamics of gene regulatory networks over gastrulation.

**Figure 1.**
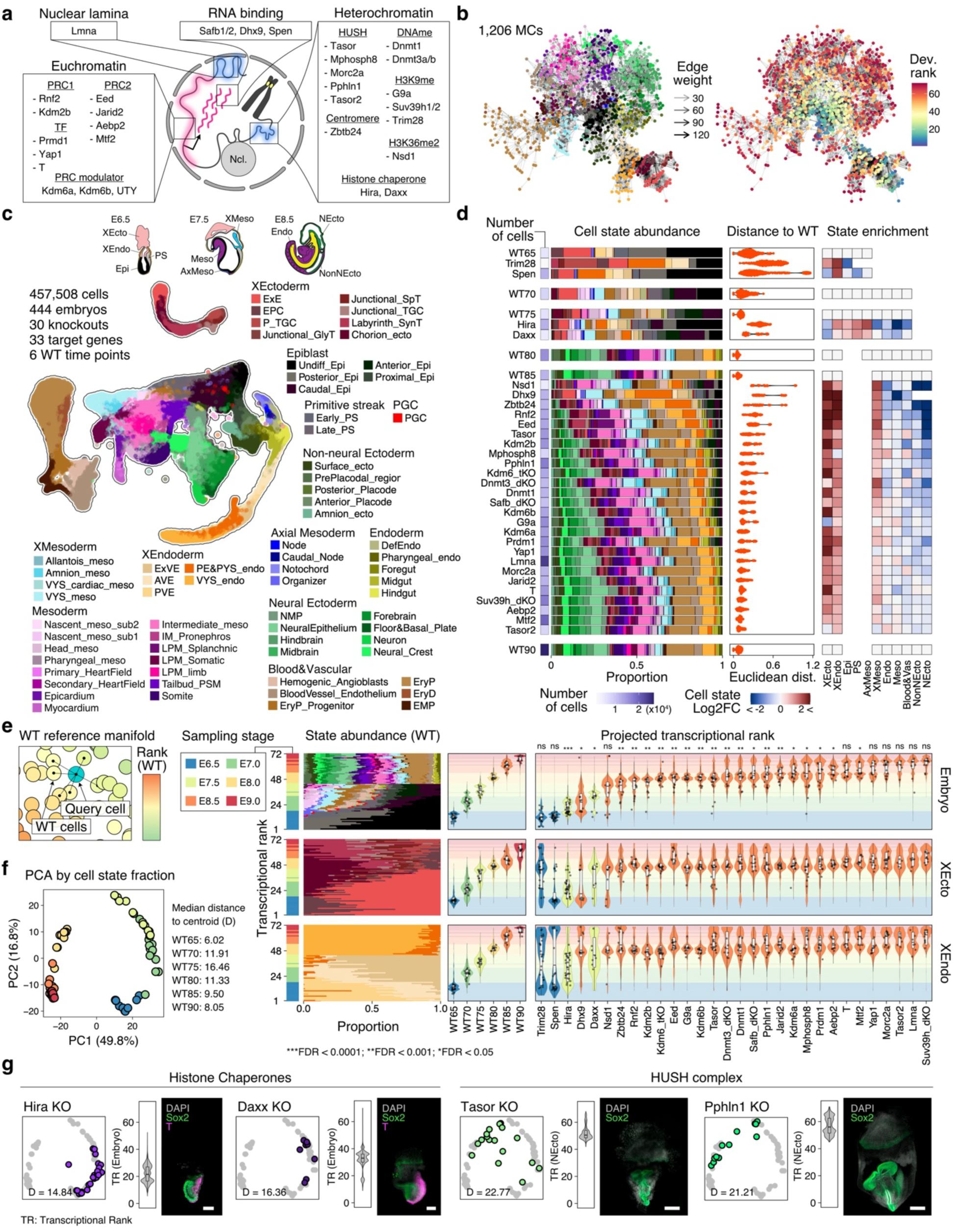
Developmental dependency maps of core nuclear regulator mutants. **a.** Schematic representing the forms of nuclear regulation explored in this study. Note that many regulators operate in multiple categories, such as the HUSH subunit Pphln1, which is also an RNA binding protein, or Spen, which is expected to function in the induction and maintenance of ChrX silencing. We prioritized informative candidates with well described biochemical functions in gene or genome regulation that had classically described mid-gestation phenotypes, under the logic that these likely impact developmental versus constitutive functions. We generated double or triple mutants in instances where individual genes are described as having highly similar or redundant functions. **b.** Metacell (MC) transition network constructed by the network flow model, colored by cell states (left) and transcriptional rank (right). Refer to Fig. 1c for the color legend of associated cell states. **c.** Projection of our full mutant data set onto a control Uniform Manifold Approximation (UMAP) of WT mouse embryogenesis, which includes 72 reference embryos from Embryonic Day (E)6.5 to E9.0 (36 females) as they proceed through 68 individual transcriptional states. Reference mapping of our mutant data provides information regarding the relative identity of recovered cells, as well as the overall cell state composition of individual replicate embryos. In total, our combined manifold consists of 457,508 confidently assigned cells, 444 embryos, 30 disrupted pathways, and 33 total mutated genes. Schematic at the top highlights the broad spatial positioning and timing of the described cell states as they relate to embryo morphology. **d.** Broad assessment of embryo cell state composition and developmental progression. **Left:** cell state proportions of individual mutant regulator cohorts alongside time-matched control samples. Purple heatmap refers to the overall number of cells recovered per data set. Regulator mutants are ranked according to the isolation time as well as the aggregate Euclidean distance of their cell state composition compared to time matched controls. **Middle:** Composition-based pair-wise Euclidean distance between each mutant embryo and its time-matched control. Epigenetic regulator mutants are frequently characterized by a larger degree of phenotypic variation, which scales with the overall severity. **Right:** Time-matched Log_2_-fold change of mean embryo composition according to major developmental lineages, including the XEcto and XEndo. Regulator mutants often show higher proportions of extraembryonic cell states compared to embryonic cell states. Within the embryo proper, the late-differentiating neural ectoderm tends to be underrepresented, with a corresponding increase in the proportion of early differentiating extraembryonic mesoderm cells. **e.** Developmental progression of mutant cells according to the transcriptional rank of control embryos, ordered by the median progression of embryonic cells. **Left:** individual query cell is projected onto the control manifold to determine its transcriptional rank according to its 20 nearest control neighbors. **Right:** Violin plots of the aggregate transcriptional rank of sampled cells as determined by the control manifold, split into embryonic (top), XEcto (middle), and XEndo (bottom) lineages. The overall composition of control samples are included as a reference to highlight the major developmental transitions that occur between each time point, including the initial induction and elaboration of the primitive streak from E6.5 to E7.5, followed by more complex processes related to elongation/somitogenesis and neural ectoderm differentiation from E8.0 to E9.0. Violin and inset colors highlight the time of sample isolation and the relative positioning of control samples, respectively. The black dot represents median transcriptional rank for each embryo, and the statistical test was performed for the median values against the stage-matched WT embryos using Wilcoxon’s two-sided rank-sum test with FDR adjustment. **f.** Replicate-level principal component (PC) projection of reference atlas according embryonic cell type composition. Replicate embryos sorted by their collection stage and can be utilized as a secondary method for assessing embryo progression. Compositional analysis also highlights an extended progression from E6.5-E7.5 and a large gap between E7.5 and E8.0-E9.0. These highlight areas of increasing growth and complexity associated with gastrulation (early) and axial elongation (late). **g.** Morphological validation of mutant embryos with early versus late phenotypes by immunofluorescence, including the inferred transcriptional rank (TR) of select tissues (embryonic or neural ectodermal) as well as the projection of individual replicates onto the cell state composition PC space. The histone chaperones Hira and Daxx both exhibit early gastrulation phenotypes that affect differentiation of the primitive streak (T, red), with Hira more severe than Daxx. The HUSH subunits Tasor and Pphln1 both exhibit complex later stage phenotypes that affect the neural ectoderm (SOX2+, green), with Tasor more severe than Pphln1. Scale bar: 100 µm for Hira and Daxx mutants; 200 µm for WT, Tasor, and Pphln1 mutants.

## Results

### An integrated atlas of genomic regulator mutants

To examine the roles of nuclear regulators during early mouse development, we leveraged a wild-type (WT) single-cell transcriptome reference atlas that samples 72 whole embryos from the onset of gastrulation (Embryonic Day (E)6.5) through anterior-posterior axis formation, neurulation, somitogenesis, and the production of early organ fields (E9.0) at 12h resolution (**Extended Data Fig. 1a-c; Methods**). Similar to previous work from our group, we employed Intracytoplasmic Sperm Injection (ICSI), here using banked female CAST/EiJ x male C57BL6/J crossed F1 (“CF1”) sperm into B6D2F1 oocytes to generate experimental and control embryos, which allows us to demultiplex and sex type individual sibling replicates according to the random transmission of dense CAST/EiJ polymorphisms (**Extended Data Fig. 1d-f**; **Methods**)^33^. We further resolved our reference into 68 distinct transcriptional cell states, and incorporated these into a metacell-based network flow model^27,39^ that captures the dynamic transitions from the pluripotent epiblast into regionalized germ layers and tissues (**Fig. 1b; Extended Data Fig. 1g-k**).

As our primary read-out involves the recovery of single cell transcriptomes, we selected candidate regulators whose genetic interactions could be read out by changes to gene expression and whose mutant phenotypes could be expected to emerge during this window (**Supplementary Table 1**): **general heterochromatin associated proteins and complexes**, including the H3K9 methyltransferases and their interactor G9a^40^, Suv39h1/2^41^, and Trim28^42–44^ as well as the DNA methyltransferases Dnmt1^45^ and Dnmt3a/b^46^, the intergenic H3K36 methyltransferase Nsd1^47^, the centromere-associated Zbtb24^48^ and the nuclear laminar-associated Lmna^49,50^; **RNA-based regulators** including core and auxiliary components of the Human Silencing Hub (HUSH complex^51^; Tasor^52,53^, Mphosph8^54^, Pphln1^55^ and Morc2a^54^) and its variant HUSH2 complex member Tasor2 ^56^, the ChrX inactivation-associated Spen^57^, as well as the co-transcriptional associated genes Dhx9^58,59^ and Safb1/2^60,61^; **replication-independent histone chaperones** (Hira^62^ and Daxx^63^); and finally, the **Polycomb Repressor Complex axis**, including core components Eed^64,65^ and Rnf2^66^ (for PRC2 and 1, respectively), recruitment-related cofactors (Aebp2^67^, Jarid2^68,69^, Mtf2^70,71^ and Kdm2b^72,73^), and the H3K27 demethylases (Kdm6a^74^, Kdm6b^75^, as well as their combined mutation, “Kdm6 TKO,” which includes the ChrY-ortholog “UTY”). For comparison, we also included three **transcription factors (TFs)** with encoded sequence specificity: the master regulators Brachyury T (T) and Prdm1 as key determinants of the embryonic mesoderm^76–78^ and primordial germ cell (PGC) lineages^79,80^, respectively, and the more broadly expressed mechanotransduction-related Yap1^81^. Notably, for regulator mutants for which standard knockout alleles have been reported, all produce viable and fertile heterozygous progeny (**Supplementary Table 1**).

To generate mutant embryos efficiently, we employed a previously described “Crispant” strategy whereby 2-5 high confidence, locus-spanning single guide RNAs (sgRNAs; **Supplementary Table 2**) are simultaneously injected into zygotes 6 hours after fertilization using donor-matched sperm^38^. In contrast to traditional breeding strategies, this method ensures high efficiency mutagenesis across the majority of cells within the majority of recovered embryos, each of which represents a distinct biological replicate; it also allows for phenotypic inspection of extraembryonic tissues, for which embryonic stem cell (ESC)-based methods do not^34,82^. While our Crispant approach provides these advantages, it does not allow for straightforward dissection of autonomous versus non-autonomous effects and likely increases the background variation. When possible, we surveyed these regulator mutants at E8.5 so that their developmental progression could be observed at a fixed time point, but proceeded to earlier timepoints when we found we could not reliably recover them later (E6.5: Trim28 and Spen; E7.5: Hira and Daxx). In general, these timepoints preceded historical reports for lethality using standard genetic approaches, minimizing the degree to which measurements would capture non-specific secondary consequences of embryo rejection (**Supplementary Table 1**). In total, we sequenced 444 embryos of cohort data (mean 12.3 embryo replicates per condition) and a total of 457,508 high confidence single cell transcriptomes, including reference embryos and the 30 regulator mutants described above (1,946-23,700 per condition, **Fig. 1c, d**). Consistent with previous reports of Crispant-based mutagenesis in mice^83^, sequencing-based analysis of editing sites across our cohort confirmed the complete absence of unedited reads in the majority of embryos samples (380 out of 386 embryos with high coverage) (**Extended Data Fig. 2a, b, Supplementary Table S1**). Further immunofluorescence-based validation on select targets further confirmed the rarity of obvious mosaicism, both at the level of the target protein or the associated epigenetic modification (**Extended Data Fig. 2c**).

### Mutant phenotypes affect distinct developmental phases

To gain a picture of our integrated atlas, we projected our mutant data onto the control manifold and measured the relative composition of cell states in each individual replicate, the inferred developmental age of nearby control cells, as well as the aggregate transcriptional profile of each assigned cell state (**Fig. 1c, d**; **Extended Data Fig. 3, 4**). Overall, mutant cell states were well matched to controls, both in terms of high cell state prediction scores and transcriptional correlation to their control states, suggesting that major features of cell identity are stable in the absence of any given epigenetic component (**Extended Data Fig. 4a-c**).

However, although cell states were on the whole “stable,” mutant embryos differed substantially in terms of their overall cell state composition (**Extended Data Fig. 4a**). Such compositional variability can be explained in part by developmental delay, which we inferred using the age of individual mutant cells by their projections to our WT manifold (which we term “Transcriptional Rank”; **Fig. 1e**). We also examined the overall projection of replicate cell state proportions to those of WT (“Cell state proportion rank”) (**Fig. 1f, Extended Data Fig. 4d-f**). Both approaches produced similar estimates of individual embryo progression (**Extended Data Fig. 4d, f**), and the average inferred delay was also consistent with previous reports from classic transgenic studies, all of which are otherwise viable as heterozygotes (**Supplementary Table 1**). Though some degree of experimental variation likely stems from our Crispant approach, we observed an association between the cell state composition variability and the phenotypic severity (**Extended Data Fig. 4g**) and low phenotypic variation in an E8.5 control sample generated by zygotic injection of Cas9 and three validated, non-targeting sgRNAs^84^ (**Extended Data Fig. 5a-e**). These data suggest minimal artifacts introduced by the injection process itself and confirm our data’s utility for integrated analysis of epigenetic regulator function within the context of the developing mouse embryo.

By cell state, regulator mutants tended to show diminished cell state proportions within the embryo proper and higher sampling of Extraembryonic Ectoderm and Endoderm (XEcto and XEndo), which undergo fewer sequential cell state transitions during this developmental window (**Fig. 1d**). Within the embryo itself, we additionally observed a trend towards the overproduction of Extraembryonic Mesoderm (XMeso) and away from the Neural Ectoderm (NEcto), which represent the earliest and latest components of germ layer induction, respectively (**Fig. 1d; Extended Data Fig. 5f-h**).

The inferred timing of developmental phenotypes spanned the full time series, with the overall severity tending to separate regulators into those with early roles in epiblast expansion and gastrulation (median rank <=E7.5) and those with later roles in the formation of a patterned, elongated body axis (neurulation/somitogenesis phenotypes, median rank between E7.5-E8.0). For example, mutants related to global histone replacement, heterochromatinization or double-stranded RNA processing (Trim28, Hira, Daxx, Zbtb24, Spen or Dhx9, respectively) arrested early, before accelerated growth or differentiation of the embryo proper, while many others associated with different aspects of DNA/Chromatin methylation generated relevant germ layers, but often with inappropriate proportions alongside disrupted morphologies and delay. It is worth highlighting that, in many cases, the overall distribution of individual replications tended to fall in between windows of increasing organizational complexity, namely the initial patterning of the primitive streak between ∼E7.0-E7.5, or in the rapid induction and patterning of axes between ∼E7.5-E8.0. In these cases, the failure to make these highly complex transitions underscore a frequent role of epigenetic regulators in facilitating major shifts in embryonic complexity, in particularly highly conserved aspects of gastrulation and axis formation. Within this framework, we also find that genes with related functions tended to produce similar phenotypes, but over ranges that appeared to reflect each component’s overall importance, including more severe phenotypes for the H3.3 chaperone Hira over Daxx, the HUSH components Tasor over Pphln1, Mphosph8, and Morc2a, and the core PRC subunits Rnf2 and Eed over their putative recruitment factors Kdm2b and Jarid2 (**Fig. 1g**; **Extended Data Fig. 6a, b**).

In comparison to these more complex responses, TF regulator mutants tended to have similar mean transcriptional ranks to those of E8.5-stage controls, with only one or several affected lineages (**Fig. 1e**). For example, T mutant embryos proceed through most of early post-implantation development but are depleted of tailbud mesoderm alongside a comparable enrichment of NeuroMesodermal Progenitors (NMPs) ^82^, whereas Prdm1 mutants are depleted for pharyngeal cell states, also as previously reported^80^ (**Fig. 1d; Extended Data Fig. 4a, 6c-f**). Prdm1 has also been reported to be an essential specifier of the primordial germ line^80,85^, but PGCs are poorly sampled in our data set, precluding statistical validation of this effect. Although Yap1 is more constitutively expressed and expected to play a role in translating local forces into transcriptional changes^86^, its mutation also did not affect the overall ability for embryos to generate cell types from all three germ layers, instead leading to gross morphological defects, delayed growth and stunted elongation, as previously reported^81,87^ (**Extended Data Fig. 6c, d, g**). It is worth noting that TF mutant responses also tend to show more global distortions in embryo progression and growth, albeit less dramatically than for many epigenetic regulators, which could also reflect secondary, non-autonomous effects downstream of lineage-specific defects.

### Distinguishing constitutive versus lineage-specific X inactivation

We next leveraged the high density of CAST/EiJ SNPS to investigate the effects of epigenetic disruption on Chr X inactivation in females (**Extended Data Fig. 7a, b**). Although the breadth of regulators tested precluded statistical testing due to each cohort being underpowered, we observed female-specific effects for the RNA binding protein Spen^20^, as well as the core PRC components Eed and Rnf2 (**Extended Data Fig. 7b**). At the level of individual state-level comparisons, Spen mutant females lacked nearly all embryonic or placental cell types and were composed almost entirely of parietal endoderm (an XEndo derivative; **Extended Data Fig. 7c**). In contrast, Eed mutant female embryos showed accelerated differentiation and depletion of placental cell types^88^ (**Extended Data Fig. 7d**).

Within single cells of our WT cohort, haplotype-resolved ChrX expression is bimodally distributed within embryonic lineages and heavily skewed to the maternal X in yolk sac and placenta, where X inactivation (XCI) is imprinted in mice^89^ (**Extended Data Fig. 1f; Extended Data Fig. 7e, f**). To more quantitatively assess subtle changes in allelic usage, we computed single cell-level XCI integrity scores as the per-cell fraction of transcripts derived from the active versus inactive ChrX and examined these ratios across cell types. Overall, we found that Spen showed constitutive biallelic ChrX expression, while the developing placenta exhibited notably leaky transcription from the inactive ChrX, even within control samples (**Extended Data Fig. 7g**). However, inactive X expression was amplified across a number of epigenetic regulator conditions, most extremely in Eed and Rnf2 mutants, but also more subtly by multiple heterochromatin-associated regulators, such as Trim28, Dnmt1, Dnmt3a/b, and Nsd1 (**Extended Data Fig. 7h; Methods**). The overall sensitivity of placental X inactivation was strikingly specific, suggesting that this tissue maintains a more dynamic form of heterochromatin with a higher degree of latent transcriptional potential.

### Epigenetic control of developmental gene expression

Our composition-based analyses confirmed that epigenetic regulators tend to produce mutant phenotypes that emerge during the early stages of embryonic differentiation, including graded phenotypes within regulators with overlapping biochemical functions. To assess how these phenotypes reflect changes in gene expression, we employed Non-Negative Matrix Factorization (NMF^90,91^) to decompose the full matrix of mutant and control transcriptional states (**Fig. 2a**; **Extended Data Fig. 8-10**; **Methods**). Briefly, NMF provides a unified framework to decompose complex gene expression signatures into coherent subgroups (Gene Expression Programs or GEPs), alongside a usage matrix that can act as a cell- or state-level activity score. For more refined analysis, we additionally performed replicate-informed state-level DESeq2 analysis against matched control data for each cell state (**Supplementary Table 3**). Notably, the resolution of our cell state assignments effectively controls for differences in developmental progression, as most individual cell states are only present for a relatively short duration (Average transcriptional rank IQR of 12.4 (SD=6.2) for 55 states recovered).

**Figure 2.**
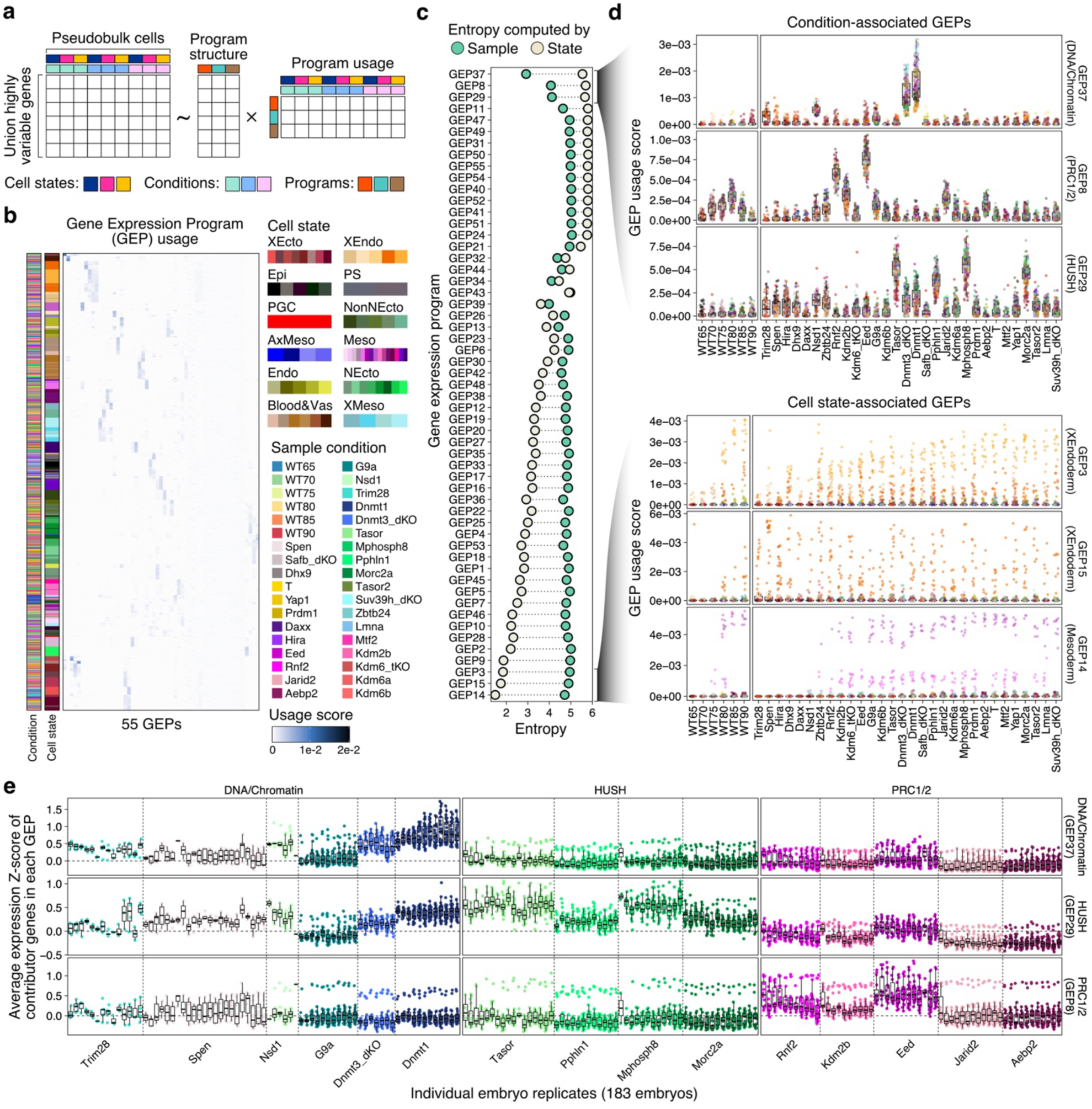
Identifying transcriptional domains of epigenetic regulation. **a.** Schematic describing our Non-negative Matrix Factorization (NMF) approach for deconvolving developmental and regulator-associated transcriptional programs. Briefly, NMF decomposes matrices into the product of two simpler matrices, which describe the overall structure of individual “Gene Expression Programs” (GEP, the weighted contribution of individual genes to a given program) as well as a “Program Usage Score,” or loading value (the relative usage of that program within a state). Here, we constructed a matrix composed of pseudobulk WT and mutant cell states by the union of 2,261 most highly variable genes calculated for each state. Descriptions of NMF optimization and validation are presented in **Extended Data Fig. 8**. **b.** Hierarchical clustering of program usage scores (loadings) for each of 55 GEPs identified using our pseudobulk gene expression matrix. Sample and state identities are highlighted on the left. In general, the overall strength of developmental GEPs drives the clustering into individual cell states rather than by individual regulator mutants. **c.** GEP prioritization driven by regulators versus cell states. Briefly, for each GEP, we calculated the entropy for program usage scores as grouped by samples (i.e., regulator mutant identity) versus cell states. GEPs are ranked by the difference between comparisons, with more regulator-specific GEPs at the top and more state-specific GEPs at the bottom. GEPs 37, 8, and 29 show lower sample-level entropy than state-level entropy, suggesting more structured program usage across combinations of regulators than by cell states. **d.** Program usage scores for the top and bottom three regulators by the entropy sorting in Fig. 2c. State-specific GEPs (bottom) showcase the dynamic interactions across different regulator classes or cell states. Each dot represents a cell state recovered from a single embryo replicate. GEP37 (DNA/Chromatin) shows significant program usage for Dnmt1, Dnmt3a/b, Nsd1, Trim28, and G9a in that order. GEP29 (HUSH) is mostly specific for the HUSH complex, with high usage scores for Tasor, Mphosph8, Morc2a, and Pphln1 in that order. Dnmt1 and Nsd1 also appear to interact with this program, but to a lesser degree. Finally, GEP8 (PRC) is descriptive of the Polycomb Repressive Complexes, ranked from core complexes Eed and Rnf2 to Kdm2b and Jarid2. G9a and Aebp2 also show significant use of this program. Finally, unlike the DNA/Chromatin and HUSH programs, the PRC GEP shows dynamic transcriptional activity over the control reference, rising from E6.5 to E8.0 before decaying at later time points. Number of state-by-replicate pseudobulk cells per condition= 71-391 (mean = 248.9; s.d. = 81.7). **e.** State-level expression summary for each of our three major regulator specific GEPs, including DNA/Chromatin (GEP37), HUSH (GEP29) and PRC (GEP8). Individual dots represent individual cell states for each replicate embryo. Replicates within their respective cohorts exhibit heightened expression of these targets, with differences between regulators reflecting either primary versus auxiliary regulatory roles (for PRC and HUSH) or subnetworks of additional regulation (DNA/Chromatin, see below). Note that the PRC program exhibits higher expression within two cell states across the mutant cohort and in wild type, which appears to represent an interaction between genes in this cohort and extraembryonic endoderm (see **Extended Data Fig. 10a**).

NMF resolved 55 individual GEPs, the majority of which described normal developmental processes and clustered control and mutant cell states together (**Fig. 2b; Extended Data Fig. 8**). To determine which programs are more descriptive of epigenetic regulation versus native development, we evaluated the overall dispersion (entropy) of each GEP’s usage scores by both regulators and cell states, reasoning that GEPs with lower regulator-level entropy would indicate greater epigenetic control over gene expression (**Fig. 2c**). There, we observed ∼19 GEPs that showed a higher degree of regulator-specific activity, the top three of which grouped transcriptional responses into clear regimes: DNA/Chromatin methylation (GEP37: including interactions with Dnmt1, Dnmt3a/b, Nsd1, Trim28, and G9a), PRC function (GEP8: Eed, Rnf2, Kdm2b, Jarid2, Aebp2, and G9a), and HUSH/RNA binding (GEP29: Tasor, Mphosph8, Pphln1, Morc2a, Nsd1, and Dnmt1; Adjusted Wilcoxon’s one-sided rank-sum test *P*-value <= 0.01; **Fig. 2d**). Each showed comparatively high enrichment for genes with previously described biological roles, genetic architectures, or chromatin modification states for each class as found in stem cells (**Extended Data Fig. 9, Supplementary Table 4**). For example, the PRC program (GEP8) shows high enrichment for genes with Suz12 binding and H3K27me3-modified bivalent genes (**Extended Data Fig. 9a-d**), while the HUSH program (GEP29) is enriched for shorter, A-rich pseudogenes, consistent with the distinct biochemical and genetic specificities of their associated complexes (**Extended Data Fig. 9e,f**)^92,93^. In contrast, the DNA/Chromatin methylation program (GEP37) is enriched for so-called “genomic defense” factors^94^ associated with male germline development (**Extended Data Fig. 9g,h**).

The overall consistency of mutant effects was also apparent when looking at replicate level GEP activity, which was highly reproducible across the DNA/Chromatin (GEP37), HUSH (GEP29), and PRC (GEP8) programs. In all three cases, the level of expression was highly similar for individual regulators and across the majority of states (**Fig. 2e**, **Extended Data Fig. 10a**). Notably, within this framework, we observed that the DNA/Chromatin program was partially activated with Spen mutant females, which we traced to failure to deactivate X-linked genes that are otherwise regulated by DNA methylation (**Extended Data Fig. 10b-d**). We recapitulated these results when using T and Dnmt1 mutant data sets^32,82^ from prior studies, which utilized mESC-based chimerism or traditional breeding, respectively (**Extended Data Fig. 10e-k**). Combined, these results show that many individual epigenetic regulators constrain development in set ways, either by blocking or augmenting differentiation. Furthermore, many regulators associated with early arrest, such as Hira, Daxx, Dhx9 or Zbtb24, do not provide clear transcriptional signatures despite major effects on developmental progression, suggesting that other mechanisms may be necessary to support embryonic viability after implantation.

### A developmentally resolved epigenetic network

A key feature of our NMF-based analysis is the ability to determine the genetic interactions between different regulators over the same genes and cell states. To do so, we assembled lineage-level interaction networks of GEP program usage across our reference and mutant data (**Fig. 3a**). Within this framework, we see clear separation of transcriptional interactions associated with constitutive and facultative heterochromatin, including subnetworks of DNA/Chromatin-based heterochromatin and the HUSH complex members. Notably, these mutant responses were largely disconnected from those of the PRC subunits, which are more correlated to embryonic time points associated with body axis formation, neural differentiation, and elongation (**Fig. 3b; Extended Data Fig. 11a-c; Methods).** From this point of view, the repressive roles of epigenetic regulators within this window appears to be partitioned between those that oversee embryo patterning itself (such as the PRCs) and those that consolidate programs associated with other developmental windows, namely the germline and early embryo.

**Figure 3.**
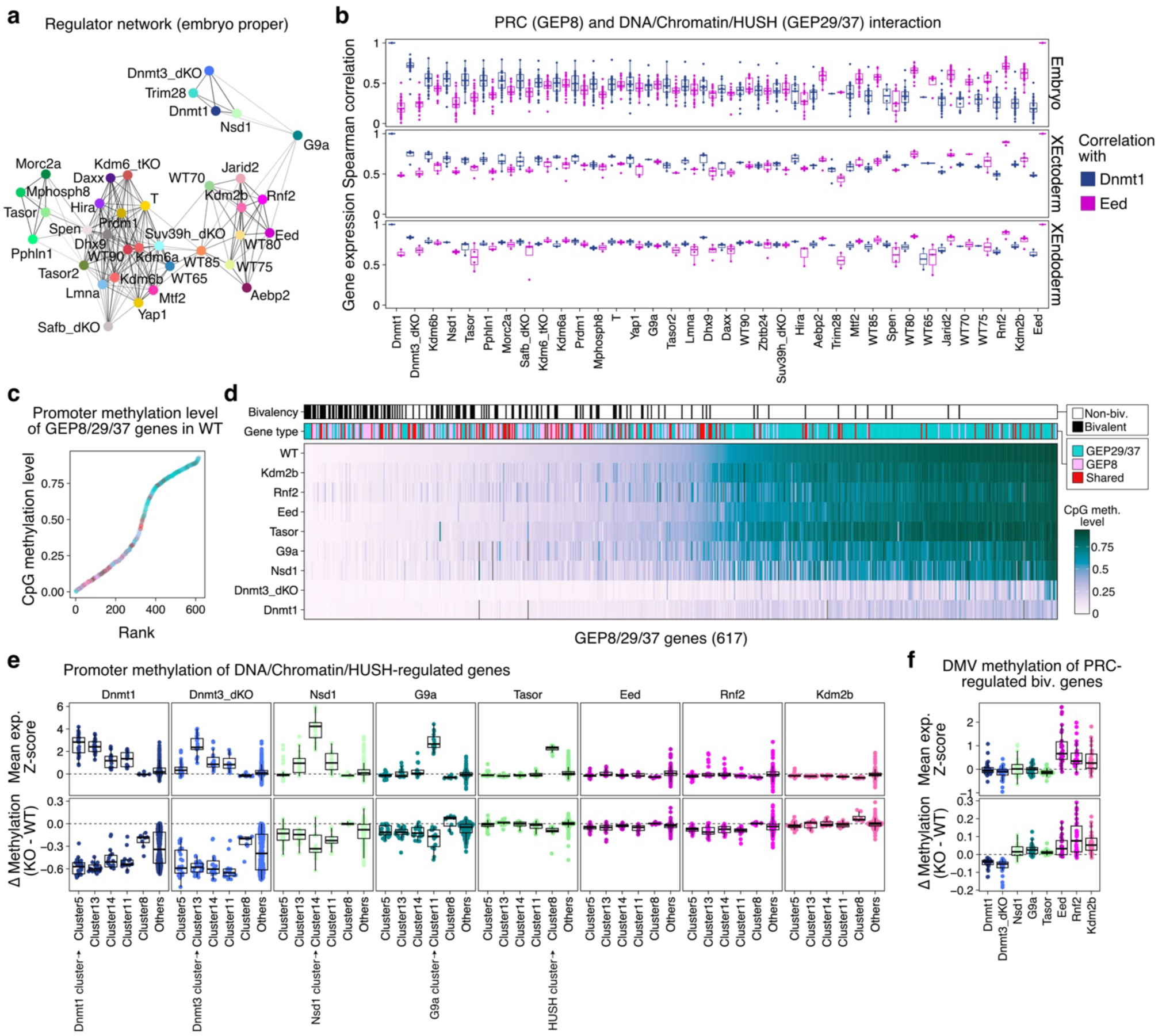
Orthogonal subnetworks oversee germline versus developmental gene regulation. **a.** An interaction network of epigenetic regulators within embryonic cell types. The top 10 low-entropy GEPs by condition (Fig. 2c) were selected and their usage score in embryonic cells were used to construct the network. Overall, regulator level transcriptional responses segregate into three obvious groups associated with DNA methylation/heterochromatin, the HUSH complex, and the PRCs. Of these, the PRCs interact heavily with select embryonic timepoints associated with axis formation and patterning (E7.5 and E8.0). **b.** Regulator-state level correlations between PRC (GEP8) and DNA/Chromatin (GEP37) expression. Here, each individual mutant cell state was tested for the degree of transcriptional correlation against the Eed or Dnmt1 mutant cohort. Pairwise correlation coefficients were computed for each cell state by replicate-average Z-score of the union of GEP8/29/37 contributor gene sets for each respective program, and this process was iterated over all recovered cell states. Genes associated with PRC recruitment show notably high correlation to the Eed regulator for the GEP8 program, and correspondingly low correlation to the Dnmt1 mutant cohort for the GEP37 program. The converse is true for GEP37 and DNA methylation/chromatin regulators, which show high correlation to Dnmt1 but low correlation to Eed. Genes such as G9a, which shows positive program usage for both sets, exhibit middling correlations to both Eed and Dnmt1, suggesting a degree of interaction in both cases. Number of detected cell states = 3-72 (mean = 49.7; s.d. = 18.3) for Embryo; 2-11 (mean = 8.7; s.d. = 2.4) for XEctoderm; 3-7 (mean = 5.3; s.d. = 1.4) for XEndoderm. **c.** Ranked promoter methylation levels of the union set of PRC, DNA/Chromatin, and HUSH target genes in E6.5 WT Epiblast. Gene promoters are colored by GEP occupancy into either GEP29/37 (HUSH and DNA/Chromatin) or GEP8 (PRC). Overall, target genes strongly separate by their mode of epigenetic regulation. **d.** Heatmap of the same gene set in **c**, with promoters ranked by methylation status in E6.5 Epiblast and mutant samples ranked by mean methylation of the feature set. Variable methylation level is visible in the mutants, including increased methylation of PRC targets (GEP8) within the left portion of the plot. Dnmt1 and Dnmt3a/b mutants show global loss of most methylation across promoters and the genome overall. **e.** Combined per gene expression (top) and promoter methylation (bottom) status for GEP29/37 subclusters highlights substantial specificity for individual co-factors within the larger umbrella of Dnmt1-based regulation, including Dnmt3a/b repressed (Cluster 13), Nsd1 repressed (Cluster 14), G9a suppressed (Cluster 11), and Tasor suppressed (Cluster 8). Each of these dynamics show statistically significant changes in DNA methylation versus the corresponding WT sample (see **Extended** Data Fig. 12a). **f.** Combined per gene expression (top) and promoter methylation (bottom) status for DMVs that intersect with GEP8 target bivalent genes (n=30 genes and 34 DMVs). In general, all three PRC subunits tested show statistically significant increases in DMV methylation, which corresponds to a loss of hypomethylation surrounding CGIs (**Extended Data Fig. 12d, e** and as previously reported^33,101^)

Beyond this high level separation, transcriptional responses within individual subnetworks also showed distinct hierarchies, with the PRC and HUSH complexes showing a high degree of synergy and the DNA/Chromatin regulators showing more heterogeneity. For the PRCs, core complex mutants exhibited consistent derepression across states, while biochemically-validated recruitment partners such as Kdm2b and Jarid2 exhibited weaker effects on the same targets in fewer cell states, a phenomenon we also observed for the HUSH subunits for their respective program^19,95,96^ (see **Extended Data Fig. 9c-e**). In contrast, individual mutant responses within the DNA/Chromatin program showed a larger variety of factor-specific roles, with individual recruitment factors falling under the shared umbrella of the Dnmt1 mutant response (**Extended Data Fig. 11d-f**). Notably, while this full gene set was enriched for germline-specific functions, particularly regulation of so-called “genomic defense” factors^94^, regulator-specific subsets appeared more refined: Dnmt3a/b, Nsd1 and G9a all exhibited unique gene subsets that shared mis-expression with Dnmt1. For example, Nsd1-dominant genes include major piRNA processing components, such as Piwil2, while genes like Asz1 receive additional support from G9a^97^ (**Extended Data Fig. 11g-j**). When examining the native expression of these genes over male germ cell development, we found co-factor-specific genes tended to be enriched for earlier stages of meiosis (early and mid-pachytene), while Dnmt1-only genes were distributed throughout the full developmental time course (**Extended Data Fig. 11j**). These expression dynamics would be consistent with a model where individual co-factors support the suppression of multiple meiosis-associated processes, while DNA methylation oversees male germ cell differentiation more globally.

### Epigenomic changes precede gastrulation

Our integrated analysis of epigenetic regulator mutants revealed an interaction network where canonical heterochromatin machinery (DNA and H3K9 methylation, as well as HUSH) encompasses diverse roles that act independently of the Polycomb Repressors, with each subnetwork showing differing degrees of specificity and cooperation between individual regulators. To determine the degree to which these phenotypic and transcriptional responses reflect corresponding changes to the epigenome, we integrated Whole Genome Bisulfite Sequencing (WGBS) data of undifferentiated Tasor and Nsd1 mutant epiblast cells alongside those from two of our previous studies, including for Dnmt1, Dnmt3a/b, G9a, Eed, Rnf2, and Kdm2b^33,98^ (**Fig. 3c-f; Methods**). Investigation of promoter methylation status in wild type immediately emphasized the degree to which the PRC and DNA/Chromatin/HUSH programs operate within distinct genomic compartments: PRC target gene promoters are largely unmethylated in WT epiblast, while the others are heavily methylated, with clear loss of global methylation signal present in our DNMT mutants (**Fig. 3c, d**).

Focused investigation of HUSH and DNA/Chromatin methylation programs confirmed that their general regulatory architecture are further encoded within the epigenome, including mutant-specific loss of promoter methylation for G9a, Nsd1, and Tasor target genes compared to Dnmt1 and Dnmt3a/b (**Fig. 3e; Extended Data Fig. 12a**). In contrast, PRC mutants showed increased methylation within larger unmethylated territories termed DNA methylation valleys (DMVs, as previously reported^99–101)^ in a manner consistent with the transcriptional activation of a subset of Polycomb-targeted genes (**Fig. 3f, Extended Data Fig. 12b,c**). In both cases, these methylation changes highlight that the epigenome can be altered prior to the emergence of complex embryonic phenotypes, and are consistent with our transcriptionally resolved interaction network, including focused PRC-based regulation of developmental gene promoters and more diverse forms of DNA methylation-based suppression (**Extended Data Fig. 12d,e**).

### Early consolidation of lineage-specific repetitive elements

Mammalian genomes have notably high repetitive element (RE) content, including concentrated satellite sequences within centromeric/pericentromeric regions as well as Long Interspersed Nuclear Elements (LINEs), Short Interspersed Nuclear Elements (SINEs), Endogenous Retroviruses (ERVs), and DNA transposons throughout the rest of the genome^102,103^. Although many of these genetic elements exhibit selfish behavior, there are countless additional examples of their co-option into host gene regulatory programs^104^. To gain deeper insights into the regulation of the repetitive genome during gastrulation, we used STARsolo^105,106^ to prioritize uniquely mapped “autonomously expressed REs,” which we can confidently distinguish from likely passive “bystander” transcription (**Extended Data Fig. 13a-d; Methods**). As with gene expression, we additionally applied a joint NMF on the full set of control and mutant cell states, enabling us to identify tissue-specific RE programs as well as those that correspond to mutant signatures. This approach further allows us to infer state-by-regulator interactions as part of the same analysis.

To overcome the sparsity of the data, we began with RE family-level analysis by aggregating mapped reads from different copies for the same family. Principal component (PC) analysis of our control reference data set separated three major groups, including clear XEndo and XEcto clusters (**Fig. 4a**). NMF of our full data set confirmed these tissue-specific signatures, including two Repeat Expression Programs (REPs 3 and 5) that captured endogenous extraembryonic development, alongside two REPs associated with HUSH- and Dnmt1/Trim28-based suppression (REPs 6 and 8, respectively, **Fig. 4b; Extended Data Fig. 13e, f**). Notably, the XEndo program REP3 also showed consistently higher usage in definitive endoderm, germ cell, and epiblast states, analogous to what has been described for these early states in endogenous gene expression space^107^ (**Extended Data Fig. 13g**).

**Figure 4.**
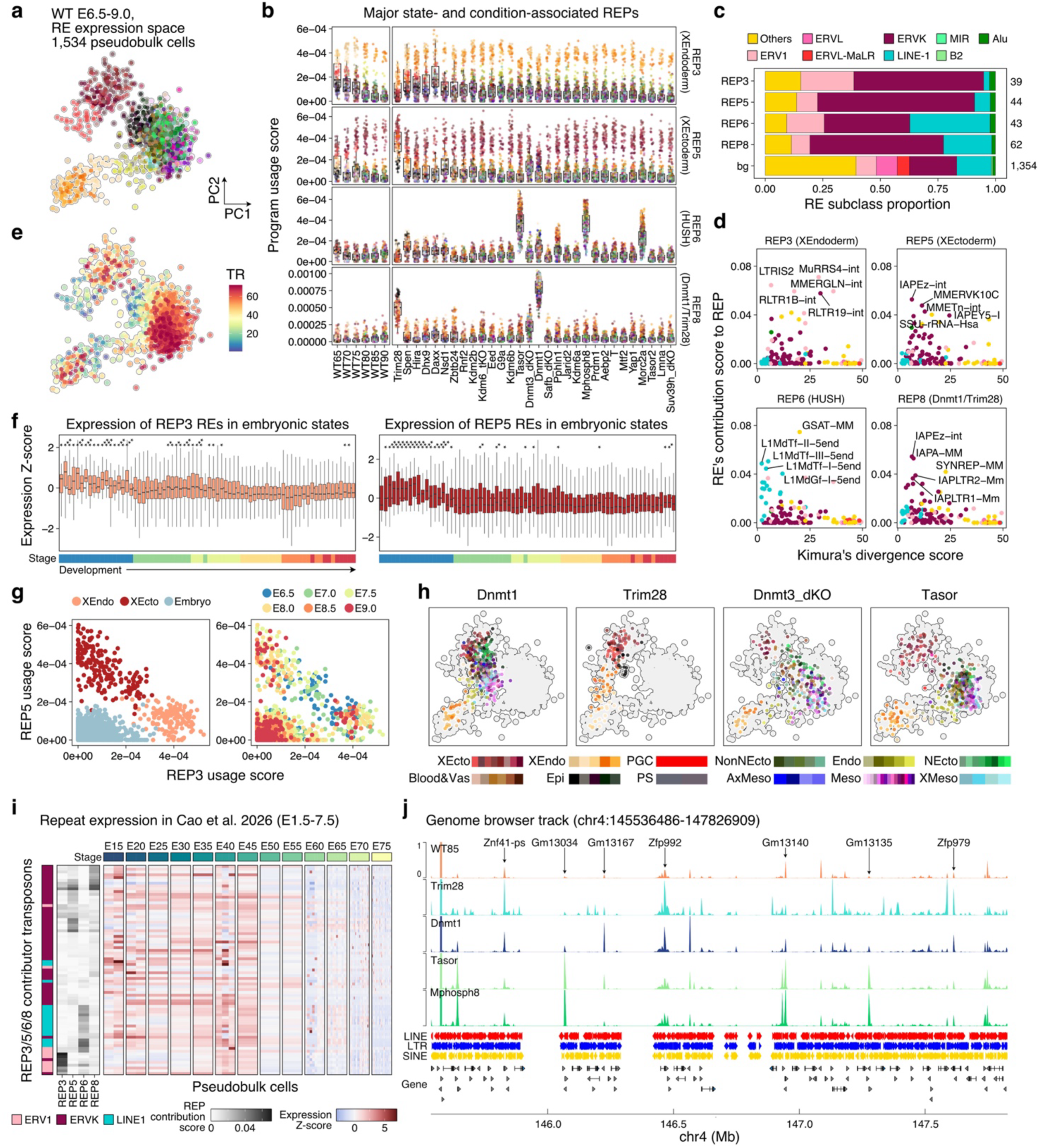
Dynamic control of endogenous and selfish repetitive DNA. **a.** Principal Component Analysis (PCA) of highly variable repetitive element families within our WT reference data, where each dot represents a given replicate transcriptional state. PC1 and 2 separate RE expression programs between placental, yolk sac, and embryonic states. Refer to Fig. 4h for the color code. **b.** Major endogenous and regulator-specific Repeat Expression Programs (REPs). Each dot represents the usage score for a single replicate-level transcriptional state colored by its cell state. REP3 and REP5 are reflective of the XEndo and XEcto (orange and red hues), respectively, and show high usage scores across the majority of regulator cohorts. REP6 and REP8 reflect dominant regulation by the HUSH complex (Tasor, Mphosph8 and Morc2a) and Dnmt1/Trim28. State color code provided in Fig. 4h. Number of state-by-replicate pseudobulk cells per condition= 117-474 (mean = 304.2; s.d. = 101.2). **c.** Broad family-level description for major repeat expression programs identified within our cohort. REP3 (XEndo) is enriched for the ERV1 family of endogenous retroviruses, and REP5 (XEcto) is enriched for ERVK family members. In contrast, REP6 (HUSH) is enriched for LINE-1s, and REP8 (Dnmt1/Trim28) is also enriched for ERVKs. Number of repeat families per program is shown on the right. bg, background. **d.** Scatterplots comparing REP contribution scores for individual RE families according to their evolutionary age (as calculated using Kimura’s divergence score^164^). REs associated with the yolk sac program (REP3) exhibit higher divergence scores than the placental one (REP5), implying that the evolutionary recency of placenta may be involved in this regulation. By comparison, RE families contributing to REP6 (HUSH) and REP8 (Dnmt1/Trim28) are evolutionarily young and enriched for extant LINE-1 and ERV-K elements such as L1MdTf and IAPs, respectively. **e.** PCA as shown in **a** but colored by transcriptional rank (TR). In general, developmentally younger states preserve low levels of extraembryonic programs before resolving towards their own. **f.** Expression Z-scores of unique REP3- and REP5-contributing RE families (14 and 19 families, respectively) within embryonic states, shown for individual embryos ordered by transcriptional rank. Dynamics confirm stable downregulation of endogenous placental and yolk sac RE programs within the embryo proper as it transitions through the initial stages of gastrulation. Each boxplot represents the expression Z-score of RE family members for individual pseudobulk cells taken at individual transcriptional ranks (14-589 RE-by-embryo-by-state combinations; mean = 248.4; s.d. = 170.3). Endoderm cells were omitted from this analysis as they show a high degree of extraembryonic program expression (see **Extended Data** Fig. 13g). FDR-adjusted one-sided Wilcoxon rank-sum test was applied for each embryo versus all the other embryos. ***P < 0.0001; **P < 0.001; *P < 0.05. **g.** Scatterplot comparing WT cell states for their respective REP3 (XEndo) and REP5 (XEcto) usage, colored by embryonic versus extraembryonic identity as well as by their developmental time point. In general, younger XEcto and XEndo states co-express REs associated with both programs, while the embryo also shows a low level of usage. **h.** Projection of mutant RE expression data onto the reference PC highlights the degree to which lineage-specific programs can become disconnected from the underlying cell state. Each dot represents a replicate level cell state. In particular, Dnmt1 and Trim28 mutants shift along both axes towards the XEcto space. By comparison, HUSH mutant regulators appear to disrupt an independent repetitive element axis that is not observed over mouse gastrulation, with less distortion of cell identity within this reference-defined space. **i.** Select RE expression over mouse preimplanation and early implantation, taken from Cao et al., 2026^115^. The expression Z-score of LTRs and LINE-1s from four major REPs in the embryo proper lineage are shown for the window corresponding to fertilization through early gastrulation (E7.5). Columns correspond to replicate-resolved cell states. Epigenetic regulator clusters exhibit transcriptional dynamics during the preimplantation window that get silenced over implantation and gastrulation. Of these REPs, the HUSH program seems skewed towards fertilization and early cleavage divisions. REPs included: Placental (REP5), Yolk Sac (REP3), Dnmt1/Trim28 (REP8) and HUSH (REP6). **j.** Genome browser track highlighting an expanded, repetitive element-rich KRAB Zinc Finger Protein cluster on Chromosome 4. In general, HUSH and Dnmt1/Trim28 mutant embryos show global destabilization of this region through activating different subsets of repetitive elements and genes. As such, it would appear that the shared transcriptional activity of these types of regions reflects their overall enrichment for autonomous REs, likely a feature of their evolutionary recency. A CPM threshold of 1 was applied to all the tracks to better visualize pervasive signals within the region.

The overall spectrum of repetitive elements that contributed to each program was notably distinct, both at the level of their specific biology as well as their relative evolutionary age (**Fig. 4c, d**). For example, the native XEndo and XEcto programs (REP3 and 5) are enriched for different ERV subfamilies: the XEndo program is enriched for ERV-1s (e.g. MuRRS4, LTRIS2, and MMERGLN), while the XEcto program is enriched for ERV-Ks (such as IAPEz, MMERVK10C, and MMETn). Interestingly, we found that XEndo REs showed higher sequence divergence scores than XEcto REs, suggesting that tissue-specific transcription of these elements perhaps mirrors each tissue’s relative evolutionary and developmental timing^108^ (**Fig. 4d**). By comparison, the HUSH subunit and Dnmt1/Trim28 programs primarily oversee suppression of active elements with very low divergence scores, with REP6 (HUSH) enriched for autonomous LINE-1 subfamilies, such as L1Md-Tf and Gf, and REP8 (Dnmt1/Trim28) primarily enriched for multiple ERV-K subclasses, including IAPA-MM and IAPEz^109–111^ (**Fig. 4d; Extended Data Fig. 13i, j**). Notably, both programs also targeted satellite repeat families, with REP6 (HUSH) biased towards pericentromeric GSAT-MM repeats and REP8 (Dnmt1/Trim28) primarily targeting the centromeric SYNREP-MM^109,112,113^.

We further observed that more differentiated cell states tended to acquire more strict RE expression patterns as they differentiate, including leaky expression of extraembryonic REs within the epiblast prior to gastrulation (**Fig. 4e-g**). To better understand how these programs may influence development, we projected each mutant data set onto our RE-based reference. We found that embryonic Dnmt1 and Trim28 KO cell states projected into the extraembryonic space, particularly in the direction of the extraembryonic ectoderm, while those in HUSH KO samples retained their overall embryonic identity (**Fig. 4h; Extended Data Fig. 13h**). This would imply a connection between Dnmt1/Trim28-based RE regulation and regulation of placental programs, as reported previously^114^, while Tasor regulates targets whose native expression may fall outside this developmental window. In keeping with this observation, REP expression over a recently generated atlas of mouse ontogeny^115^ showed that extraembryonic, Dnmt1/Trim28, and HUSH-specific target REs are all endogenously expressed during fertilization and the preimplantation (**Fig 4i**). Collectively, these results showcase diversity of element-regulator specific interactions that act to suppress these elements prior to the onset of gastrulation.

### Gastrulation suppresses nearly all autonomous RE transcription

We next analyzed RE expression at single copy resolution to better understand their developmental dynamics and potential effects on nearby genes. First, we asked if the observed RE expression signals are driven by derepression of the elements themselves. To do so, we examined the “autonomous” versus “bystander” nature of each recovered RE transcript^116^: here, “autonomous” refers to the transcripts derived from the element themselves, such that they generally align to the sense direction and are similarly active inside or outside of host genes, while “bystander” refers to carry-over of elements as part of pre-processed host mRNAs (**Extended Data Fig. 13a**). Comparative embedding of sense and antisense-only UMAPs by single-copy RE count matrices provided examples of both scenarios: the sense RE alignment showed greater separation of extra-embryonic and regulator-associated states from the gastrulating embryo, while bystander alignments were heavily skewed to transcribed introns (**Extended Data Fig. 13k-m**). Here, we find that extraembryonic and mutant RE expression patterns predominantly reflect autonomously expressed RE activity, while the majority of post-gastrulation RE transcripts within the embryonic lineage reflect bystander REs captured within introns **(Extended Data Fig. 13n**). In this manner, the developing embryo proper can be distinguished from the extraembryonic tissues by the robust suppression of autonomous RE activity.

We next investigated the potential of deregulated REs to impact local gene expression^104,117^ by exploring the effects of repetitive element activity as a function of linear distance – including REs embedded within introns (**Extended Data Fig. 13k-n**). Overall, we observed minimal proximity-based relationships for most RE-by-gene comparisons, with the majority of nearby genes maintaining endogenous transcriptional levels despite the nearby presence of an activated RE (**Extended Data Fig. 14a-d**). Although this finding highlights the exceptional and comparatively rare nature of RE-based co-option, we found that condition-specific REs and highly variable genes still tended to co-occur in proximity to one another, particularly for Dnmt1 and HUSH reactivated LINEs and Trim28 reactivated ERVKs (**Extended Data Fig. 14a-d**). Closer inspection showed that the majority of these responses were unique to different regulators and elements, such that Dnmt1/Trim28 and HUSH-specific responses minimally overlapped (**Extended Data Fig. 14a-d**). Nonetheless, we found multiple instances where independently reactivated retrotransposons and genes appeared to converge on the same genomic region, particularly for highly repetitive gene clusters such as those associated with KRAB Zinc Finger Proteins (KZFPs) (**Fig. 4j**). This tendency for independent elements to affect nearby genes within the same loci highlights the natural opportunity for local reinforcement between parasitic elements and host-suppressive mechanisms within comparatively unconstrained regions of the genome^118–120^.

### Deactivating multiple programs compounds embryonic phenotypes

Many nuclear regulator mutants within our dataset induce germline programs, processed pseudogenes, and/or autonomously expressed retrotransposons, raising the question as to how or why activation of these seemingly unrelated genomic compartments is deleterious to embryo patterning. Notably, although key REP usages in Dnmt1/Trim28 and HUSH mutants first appear to be largely independent, we observed several lineage-specific interactions (**Fig. 5a, b**). In particular, HUSH mutants exhibit substantially higher REP6 transcription within the placenta and yolk sac compared to matched embryonic cell types, while the same targets are more consistently and weakly deregulated across all three lineages by Dnmt1 and Trim28 mutants as well (**Fig. 5b; Extended Data Fig. 15a**).

**Figure 5.**
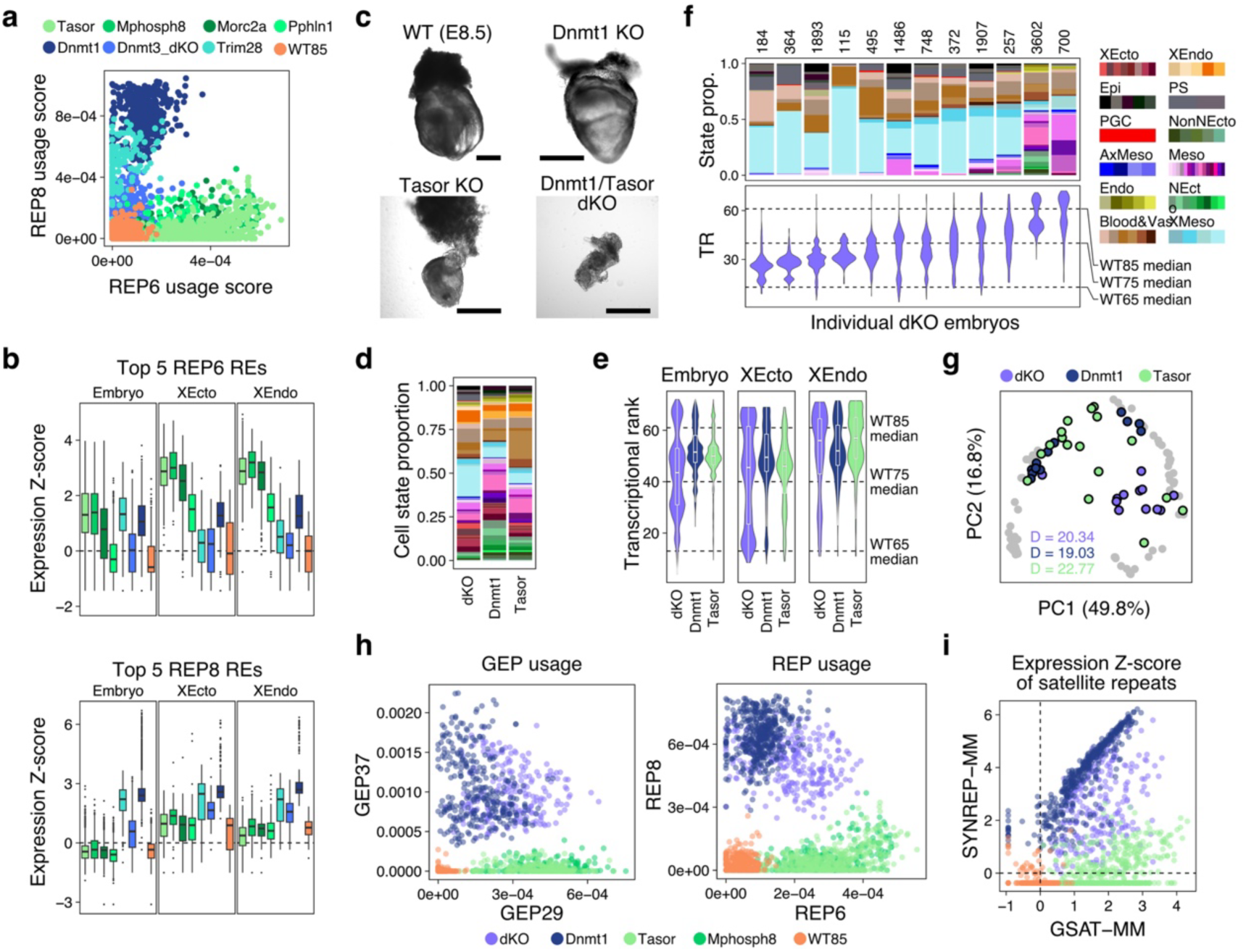
Heterochromatin-associated programs collectively support embryonic progression. **a.** Scatterplot showing REP6 (HUSH) and REP8 (Dnmt1/Trim28) usage in individual replicate states. In general, these two programs are independent, with Dnmt1 showing weak correlation and interactions with both programs. **b.** Expression Z-score within pseudobulk cells for the top 5 most contributing RE families to the HUSH (REP6) and Dnmt1/Trim28 (REP8) programs. These REs are expressed to near equivalent levels between HUSH core subunit and Dnmt1 mutants within the embryo, but are HUSH-exclusive targets within the XEcto and XEndo. By comparison, HUSH subunits do not interact with the top REs within the Dnmt1/Trim28 program, which instead show latent transcriptional activity with extraembryonic tissues (see expression of wild type in orange). Number of combinations of cell state, embryo replicate, and RE family per condition in (Embryo, XEcto, XEndo) = (1,805, 315, 225)for Tasor; (1,110, 180, 100) for Mphosph8; (1,765, 285, 135) for Morc2a; (1,635, 235, 160)for Pphln1; (115, 365, 270) for Trim28; (1,065, 100, 100) for Dnmt3a/b; (1,765, 295, 200) for Dnmt1; (1,460, 100, 75) for WT85. **c.** Transmitted light images of single Dnmt1 and Tasor mutants in comparison to WT and the combined mutant (DKO). Scale bar, 500 µm. **d.** Stacked bar plots showing the overall cell state composition of combined Tasor, Dnmt1 mutated (DKO) embryos compared to single mutants (single mutants also shown in Fig. 1d). In general, double mutant embryos show diminished embryonic progression, leading to the overproduction of extraembryonic mesoderm states (blue) and blood (brown) and minimal differentiation of neural ectoderm states (green). Refer to Fig. 5e for the color code. n = 16,514 cells for dKO; 11,852 cells for Dnmt1; 14,879 cells for Tasor. **e.** Predicted transcriptional rank of mutant cells, separated by embryonic, placental, and yolk sac identity. In general, double mutant cells show disrupted embryonic progression and differentiation defects, leading to the retention of less differentiated cells associated with earlier transcriptional ranks. n (Embryo, XEcto, XEndo) = 12,123, 2,714, 1,677 cells for dKO; 9,529, 1,183, 1,140 cells for Dnmt1; 12,461, 1,176, 1,242 cells for Tasor. **f.** Replicate-level stacked barplots (top) and transcriptional ranks (bottom; TR) of embryonic lineage for our 12 combined double mutants. 10 of the 12 mutant embryos show severe delays. The number of cells per embryo is noted on top. **g.** PCA-based based projection of cell state composition, with distance to centroid (D) included (inset). Previously shown E8.5 Dnmt1 (dark blue) and Tasor (green) projections are included for reference. Together with **f**, these plots highlight the increased phenotypic severity of the delay phenotype when two weakly interacting genomic suppression pathways are simultaneously disrupted. **h.** Replicate state-level scatterplots of the Dnmt1/Chromatin programs (GEP37 and REP8) and HUSH programs (GEP29 and REP6) as they compare between individual and double mutants. In general, the effects of double mutations lead to combined usages of programs associated with each individual regulator. **i.** Replicate state-level scatterplots comparing minor/centromeric SYNREP and major/pericentromeric GSAT expression within single and double mutants. Dnmt1 and Tasor mutant states show that expression of these two centromeric compartments is linearly correlated, with dominant Dnmt1-SYNREP and Tasor-GSAT regulation. Combined mutation of these pathways leads to combined derepression of both compartments to near equivalent levels.

The weak interactions between Dnmt1/Trim28 and HUSH regulation prompted us to investigate their collective effect on embryonic development, as well as the degree to which each regulator may compensate for the other’s endogenous functions. To do so, we disrupted both Tasor and Dnmt1 simultaneously and sampled a cohort of 12 E8.5 embryos by scRNA-seq **(Fig. 5c, Extended Data Fig. 15b)**. Compared to either single mutant, double-knockout (dKO) embryos were severely arrested, displaying a highly disorganized morphology that skewed heavily towards the XMeso lineage, with very little progression of the embryonic germ layers (**Fig. 5d-g; Extended Data Fig. 15c**). When individual embryos were sorted by their transcriptional ranks, the most arrested embryos consisted of undifferentiated primitive streak, blood/vascular, and XMeso cell types, suggesting that the loss of two suppressive regulatory pathways substantially enhances the severity of the developmental phenotype (**Fig. 5f**). Transcriptionally, projection of mutant gene expression data onto our pre-built NMF models found that both regulator programs were simultaneously activated, with minimal additional contributions beyond their individual targets **(Fig. 5h; Methods**).

Even as purely additive responses, simultaneous Dnmt1/Tasor disruption did lead to some previously unobserved phenomena. For example, we see a clear partitioning of primary repression between minor, centromeric repeats (SYNREP) and major, pericentromeric transcription (GSAT) as part of Dnmt1- and HUSH-specific REPs (**Fig 5i; Extended Data Fig. 15d**). Within the dKO, both compartments are induced to roughly equal levels, suggesting more broad loss of epigenetic control over these key features of mammalian chromosomes (**Fig. 5h**). Notably, this additive effect also holds for RE-gene pairs, which act independently within single mutants but are both reactivated in DKOs (**Extended Data Fig. 15e**).

### Genome suppression licenses entry into the phylotypic period

Biochemically, the HUSH complex is reported to have specific roles in regulating A-rich and/or monoexonic transcripts, which we observe in both gene and repeat expression spaces^92^, and lacks obvious connectivity to gastrulation-associated programs. Nonetheless, Tasor disruption causes a complex embryonic phenotype, including the underproduction of neural epithelium and loss of a robust anterior-posterior morphology^53^ (**Fig. 6a**). To interrogate how differentiation breaks down over the course of gastrulation, we generated a denser time series of HUSH mutant differentiation, including additional 10 and 14 embryos at E6.5 and 7.5, respectively (**Extended Data Fig. 16a**). Projection of our KO time series onto the control reference revealed a continuous transition from pluripotent to differentiated cell states, but progressive build up of earlier transcriptional ranks beginning around E7.5 that includes prolonged retention of epiblast and primitive streak cells (**Fig. 6b-d, Extended Data Fig. 16b**). Despite the progressive nature of this phenotype, our gene and repeat NMF models suggest that both target genes and REs remain constitutively active throughout the sampled timepoints, representing a continuous and persistent transcriptional noise rather than the accumulation of obvious cell type specific responses (**Fig. 6e**).

**Figure 6.**
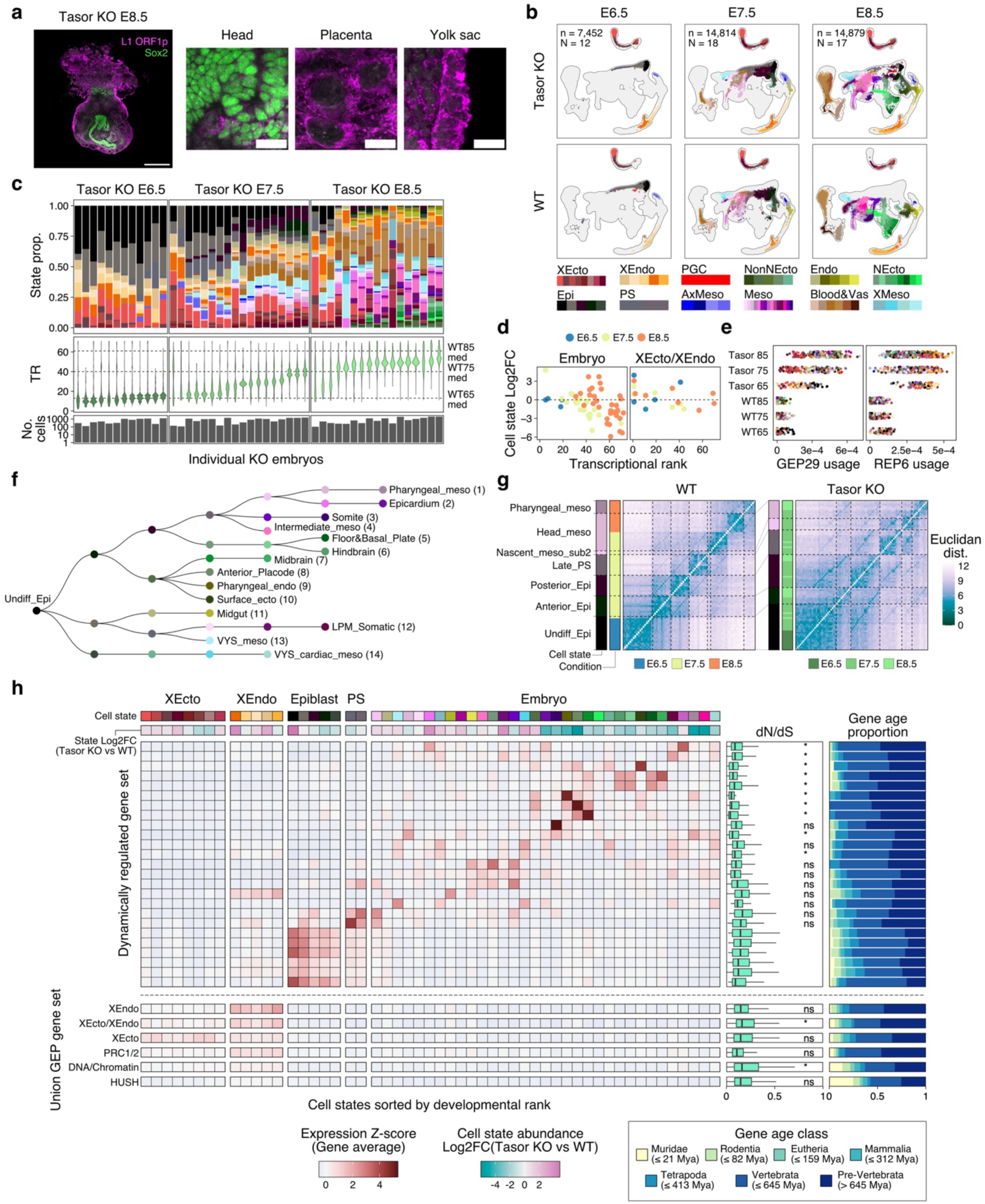
Genomic silencing licenses entry into conserved transcriptional programs. **a.** Morphology of a Tasor mutant embryo isolated at E8.5, whole mounted and stained for the Neural Ectodermal marker Sox2 and for LINE-1 ORF1p. Even though the embryo proper shows comparatively lower levels of LINE-1 dysregulation compared to the placenta and yolk sac, it shows substantially disrupted morphology, including an unclear anterior-posterior axis and a disrupted proportion of mesodermal and neural states. Scale bars = 200 µm for the whole embryo images; 20 µm for the zoom-in images. **b.** Projections of our full Tasor mutant and WT timeseries onto our reference manifold, separated by isolation time and colored by cell state. While reference embryos proceed from early differentiation states such as the primitive streak to more established germ layers, Tasor mutants maintain undifferentiated pluripotent epiblast cells and skew more heavily towards mesodermal and extraembryonic mesodermal states. By comparison, states associated with the neural ectoderm are depleted and delayed. The Tasor E8.5 data is also presented in **Extended Data** Fig. 3. **c.** Per replicate stacked bar plots of cell state, ranked by each replicate’s respective transcriptional rank (TR). It highlights the gradual accumulation of delay phenotypes through gastrulation. The Tasor E8.5 data is also presented in **Extended Data** Fig. 4a. **d.** Stage-matched Log2-fold change in cell state abundance, colored by isolation time point and split by embryonic versus extraembryonic. As development proceeds, there is an increasing accumulation of earlier cell states and underproduction of later ones. This phenomenon is not apparent within the extraembryonic lineages. **e.** Per replicate state-level usage plots for GEP29 and REP6, gene and RE-associated programs regulated by HUSH. While GEP29 shows subtly increased usage over time, REP6 is stable from E6.5 onwards. **f.** Summary tree of the 14 non-redundant reference trajectories as the shortest paths from undifferentiated epiblast to E8.5 cell states, as predicted by our network flow model (see Fig. 1). This tree was subsequently utilized to infer time-resolved transcriptional dynamics as they compare between the control reference and our Tasor mutant time series (see **Extended Data Fig. 16c-e**). **g.** Euclidean distance matrices of normalized gene expression counts along the path from undifferentiated epiblast to pharyngeal mesoderm, comparing the transition profiles between control and Tasor KO embryos. Tasor KO embryos differentiate with less clear transcriptional boundaries, as evident by the lower state level distances between the terminal mesodermal state and the initial epiblast. Off-diagonal stripe patterns derive from cell state comparisons within the same embryos, but trends hold even when these sample-matched state pairs are removed (see **Extended Data Fig. 16f**). **h.** Summary heatmap of mean gene expression Z-scores describing the relationship between differentiation and gene regulatory constraint over the course of Tasor mutant embryogenesis. At the top rows, cell states are ordered by their mean transcriptional ranks, as well as the Log2FC in state occupancy between Tasor mutant embryos and the control reference. For the cell states that span multiple time points, the lowest FC values were selected. Y-axis highlights dynamically regulated gene sets as identified by trajectory inference (**Extended Data** Fig. 16c**-e**). Right: The overall constraint (measured as the mean dN/dS ratio) and relative gene age of each set. Significance of the dN/dS scores is tested by FDR-adjusted two-sided Wilcoxon rank-sum test compared to the union gene set for the six earliest epiblast-enriched gene sets. In general, earlier differentiation states associated with the primitive streak, definitive endoderm, and other initial products show comparably lower constraint (higher dN/dS ratios) compared to underproduced later states, where the gene ages rise and dN/dS ratios fall. At the bottom, GEPs selected from Fig. 2 are provided for comparison, and highlight the relatively higher dN/dS ratios and enrichment of evolutionarily young genes of the HUSH and DNA/Chromatin programs, particularly in comparison to the PRC1/2 program. ns, not significant; \**P* <= 0.05 (Wilcoxon’s two-sided rank-sum test, FDR adjusted).

We next leveraged our metacell flow model to assemble a graph of cell state transitions from the pluripotent epiblast forward in embryonic WT cells and obtained dynamically regulated gene sets along each developmental graph path (**Fig. 6f; Extended Data Fig. 16c-e; Methods**). There, we observe that mutant cells tend to maintain higher transcriptional similarities to earlier states, weakening the stability of multiple cell state transitions and preserving a higher similarity to the undifferentiated epiblast cell (**Fig. 6g; Extended Data Fig. 16f, g**). This lack of robust differentiation appears to reflect two co-occurring processes: incomplete suppression of transcripts associated with earlier states (including REs), as well as a failure to stably induce lineage-specific transcripts (**Extended Data Fig. 16g**). Using RNA velocity^121^ to infer transcriptional flux also supported this property, with Tasor mutant embryos exhibiting diminished separation between distinct lineages, leading to less discrete cell state transitions that compound over developmental time (**Extended Data Fig. 16h**).

Notably, although our other regulator mutants were only sampled at single time points, we find that transcriptional bleedthrough between progenitor and differentiated states is a common feature of epigenetic disruption, particularly for those with aggressive developmental phenotypes (**Extended Data Fig. 17a**). Closer examination of each progenitor cluster according to the evolutionary character of its expressed genes revealed a stepwise increase in genetic constraint over time, which recapitulates a range of studies that support the developmental hourglass model^122–125^ (**Fig. 6h**). Intriguingly, the break point between overrepresented and underrepresented cell states within our Tasor mutant data appears to directly mirror this transition, leading to the retention of earlier, comparatively unconstrained states and a failure to stably differentiate into more conserved ones (**Fig. 6h**). These evolutionary behaviors were also consistent with our GEPs, where extraembryonic cell types as well as our DNA/Chromatin and HUSH associated modules are all enriched for newer, less constrained genes, while PRC-regulated genes are relatively more conserved and contribute to cell states that emerge around the phylotypic period (**Fig. 6h; Extended Data Fig. 17b**). As such, epigenetic suppression mechanisms tend to separate into those that suppress fast-evolving, evolutionary new genes and REs (such as Dnmt1s, HUSH, and other chromatin regulators) and those that mainly function to maintain body axis patterning (such as the PRCs).

## Discussion

We present a unified framework for evaluating the developmental roles of nuclear regulators, including broad classes whose molecular and biochemical functions are well understood but challenging to connect to developmental phenotypes. Overall, we find that many of these play critical roles in aspects of embryo implantation, differentiation, or gastrulation, including frequently observed progression defects that lead to disordered patterning. With our highly controlled perturbation pipeline, we re-establish the value of developmental genetics as a core component for evaluating each regulator’s relative importance in maintaining transcriptional control and cell identity. Our ability to evaluate genes, repetitive DNA, and processes such as XCI simultaneously also provides helpful context for prior work conducted *in vitro*, where evaluating the relative effects between different classes of regulators can be harder to compare among systems and studies.

More generally, our analysis highlights the dynamic interactions between canonically epigenetic regulators. Across multiple levels of analysis, we identify a striking phenotypic partitioning between regulators associated with pre-implantation and germline regulation (pre-phylotypic) – such as Dnmt1, Dnmt3a/b, Trim28, G9a, Nsd1, and the HUSH complex – and those that maintain dynamic control over embryo patterning (phylotypic), namely those associated with the Polycomb Repressive Complexes. Ultimately, these relationships imply a developmental logic to the organization of the mammalian genome, with the gastrulating embryo representing a pivot from comparatively plastic and diverse regulatory processes to the highly conserved induction of organizational programs^126,127^. This phenomenon would explain why many distinct forms of misregulation nonetheless manifest as the same generic phenotypes, namely arrest points in windows of otherwise rapidly increasing cellular complexity, as well as the tendency to impact the late emerging, highly constrained neural lineage.

In the absence of detailed time series data, we cannot discriminate the primary response to epigenetic regulator loss from its secondary effects for the majority of our target genes, and many of our observations reflect correlations between modes of transcriptional dysregulation and the relative timing of emergent developmental defects. It is also very possible that these disruptions in developmental progression speak to dedicated pathways, such as those associated with viral RNA sensing, that may become activated during this window and impact developmental progression^112,128,129^. Overall, our temporal analysis of the Tasor mutant phenotype does provide a framework for considering these trends, including gene expression changes within evolutionarily young regions of the genome that precede the gross morphological disparities that emerge upon entry into the phylotypic period. Our deeper investigation of Tasor could also explain how developmental responses could be feasibly shared across species even if the targets themselves change. For example, within our Tasor mutant time series, we find that a comparatively limited, yet highly species-specific transcriptional signature can disrupt differentiation without directly regulating developmental genes. Both LINE-1s and pericentromeric repeats also represent conserved HUSH targets between mouse and human despite substantial sequence divergence^112,113,117^, representing yet another intriguing dynamic between selfish elements and an essential host function that likely reflects a key feature of their evolutionary interplay. Collectively, these observations offer the rising opportunity to connect what we know about epigenetic regulation to development, including how evolution and genome organization may constrain their relationship.

The interconnectivity between gene expression programs, developmental processes, and phenotypic response is further emphasized by the comparative resilience of the extraembryonic tissues, which maintain evolutionarily plastic features during the same window as the embryo becomes genetically and morphologically constrained^130–132^. Our analysis shows that the placenta and yolk sac exhibit multiple forms of permissive transcription, including endogenous RE programs and within the inactive X chromosome, as well as more extensive HUSH-mediated suppression. These surprising examples of extended transcription-based regulation, endogenized RE programs, and diminished developmental constraint appear to represent intrinsic properties of these tissues that would naturally confer robustness against epigenetic perturbation, underscoring the natural trade-off between evolutionary canalization and the potential for innovation^37,132^. They may also reflect the unique epigenomic status of these tissues prior to their differentiation, as well as their comparatively convergent evolutionary behavior across species^38,131,133^. Focused investigation into the epigenetic regulation of extraembryonic cell types, both further into their development and in other species, will better clarify the contributions of these novel reproductive tissues to mammalian genome evolution.

Although our survey identified several key hubs of epigenetic control, we do not address the precise molecular mechanisms through which different forms of genome regulation communicate to support development, including many regulators that are required for the earliest stages of gastrulation but exhibit few obvious transcriptional changes. Multiple in vitro and in vivo studies have highlighted the degree to which the transition from pre-to post-implantation embryogenesis involves genome-wide changes in nuclear morphology and chromatin state, including major swings in Polycomb activity from globally to locally distributed^134–137^. Still other findings have observed key interactions between Polycomb and other forms of heterochromatin at the level of parental imprinting or germline gene control that are also likely present in our data as more subtle transcriptional or allele-specific signatures^98,130,138,139^. Nonetheless, the nature and timing of this transition in mammals may represent a defining molecular feature that partitions developmental programs and determines their distinct evolutionary constraints. Future work that more directly seeks to translate the molecular organization of genetic elements towards the overall phenotype may provide a unifying explanation, as will efforts that evaluate the diversity of these dynamics across species.

## Acknowledgements

We thank all members of the Smith, Aktas, Sumigray and Kretzmer labs for their intellectual and technical support of this manuscript, as well as our colleagues at the Yale School of Medicine and MPIMG for their supportive feedback. We also thank Dr. Yui Uchida for valuable feedback about the manuscript. We thank the animal caretakers and technical staff of the Yale Animal Research Center, the MPIMG Animal Facility, and the MPIMG Transgene Unit for overseeing the maintenance and wellbeing of experimental animals. Research in the Smith Lab is supported by the NIH Director’s New Innovator Award (DP2HD108774), the Mathers Foundation, the Chen Innovation Award, and the Max Planck Society. Research in Aktas lab is supported by Max Planck Society and the Dhx9 KO embryo sequencing costs were partially covered by the DFG grant (Project number: 456668871) awarded to T.A. Y.K is supported by EMBO postdoctoral fellowship (ALTF 315-2024).

## Methods

### Animal experiments

CAST/B6 F1 mice were generated in-house by breeding CAST/EiJ strain female mice with C57BL/6J strain males. Swimming sperm were isolated from the caudal epididymis for males (>2 months of age) in M2 media (Millipore), decapitated by brief pulse sonication in a Branson Sonifier with double stepped tip (Branson), and stored at −40°C in 25 μl aliquots for use within 6 months to a year of collection. We used 8-week-old male CAST/B6 F1 as sperm donors for ICSI and 6–12-week-old B6D2F1/J (BDF1) female mice to provide oocytes (both from the Jackson Laboratory and/or Janvier). 22–24g female CD-1 mice were purchased from Charles River Laboratory and used as pseudopregnant donors, and Swiss-Webster vasectomized males (9 weeks old) were mated to CD-1 females. All procedures were performed in specialized specific-pathogen-free facilities, followed all relevant animal welfare guidelines and regulations, kept on a 12 h light-dark cycle from 6 am to 6 pm, and provided a standard diet ad libitum. Embryos were collected without considering sex, but individual sexes were analyzed after sequencing (not shown). Interventional procedures necessary to conduct this work were approved by the Yale University Committee on the Use of Animals in Research and Teaching (IACUC, Protocol #2026-20357) or by the LaGeSo Berlin (Protocol #G0098/23).

### Generation of wild type and mutant mouse embryos

Experimental embryos were generated as previously described^33,34,98^. Briefly, BDF1 strain females (Charles River Laboratory strain code 099) were superovulated using ∼100 µl of CARD HyperOva FD (Cat. #KYD-015-EX, Cosmos Bio) and 5 IU/mouse of human chorionic gonadotropin (Cat. #HOR-250, Prospect Protein Specialists) with injections separated by ∼46 hours. MII stage oocytes are then isolated, cleaned of cumulus cells using M2 media with hyaluronidase (Cat. #MR-051-F, Millipore sigma), and stored on a petri dish in 25 µl drops of advanced KSOM (Cat. #MR-101-D, Millipore sigma) with mineral oil. Oocytes were then fertilized with thawed CAST/B6 sperm using piezo-assisted intracytoplasmic sperm injection (ICSI). Once zygotes reached pronuclear stage 3 (PN3), 200 ng/ml of Cas9 mRNA and 100 ng/ml of 3-4x sgRNA pools (1ug/sgRNA) targeting different exons spanning the gene of interest were microinjected into the zygote cytoplasm. Cas9 mRNA and sgRNAs were generated via PCR using T7-promoter containing primers and 2x Phusion Master Mix (Cat. # F531L, Thermo Scientific) to create a template for in vitro transcription, followed by the generation of RNA using either the MEGAshortscript T7 Transcription Kit (Cat. # AM1354, Invitrogen) or HiScribe T7 High Yield RNA Synthesis Kit (Cat. #E2050S, NEB). IVT reactions were purified using the Monarch RNA Clean up Kit (Cat. #T2040L, NEB), and resuspended in Injection Buffer (5 mM Tris buffer, 0.1 mM EDTA, pH 7.4). At E3.5, cavitated blastocysts were transferred into CD-1 strain pseudopregnant females (E2.5) by laparotomy, followed by embryo isolation at the desired time point.

### scRNA-seq of whole embryos

Whole mutant embryos were isolated from surrogate female mice and cleaned of maternal tissue at the desired time point. Preliminary deciduae were collected in a petri dish with 1x PBS, maternal tissue was carefully removed without damaging the extraembryonic tissue, and then serially washed through several drops to remove maternal tissue. To eliminate bias in sample selection, we aimed to sample 10 or more embryos per time point, sorting the full clutch of embryos and prioritizing the largest and least damaged candidates to avoid over-estimating developmental delay phenotypes. Otherwise, we employed no additional selection criteria.

Dissected and cleaned embryos were transferred as a clutch into a 200 µl drop of 0.2% BSA/PBS and dissociated into single cells by replacing the 0.2% BSA/PBS with TrypLE (Cat. #12604013, Invitrogen) and incubated for 10 minutes at 37 °C. After the incubation, embryos were mechanically dissociated using a P200 pipette and incubated >2x for 5 minutes. The cell suspension was quenched by serially adding and collecting 200 µl of 0.2% BSA/PBS and carefully resuspending cells to collect into a LoBind Eppendorf tube until the final volume was ∼1 mL (Cat. #22431021, Eppendorf). Cells were filtered into a new LoBind Eppendorf tube using a Scienceware Flowmi cell strainer, 40 µm (Cat. #10032-802, VWR) and centrifuged for 5 min at 1000 rpm at 4 °C. Following centrifugation, ∼800 µl of supernatant was removed and replaced with 800 µl of 0.2% BSA/PBS and centrifuged once more for 5 min at 1000 rpm at 4 °C. Once again ∼800 µl of supernatant was removed, and cells were gently resuspended in ∼200 µl and centrifuged for 5 min at 1000 rpm at 4 °C to concentrate the pellet. An additional 100 µl was removed for a final 100 µl cell suspension used to count on a hemocytometer. Cells were then subjected to single-cell RNA sequencing (10x Genomics, Chromium Single Cell NextGem 3’ v3) with a target cell recovery of up to 12,000 sequenced cells per emulsion reaction. Single cell libraries were prepared following the protocol with a cDNA and PCR indexing amplification cycle of 10 and examined on a D5000 and D1000 TapeStation System, respectively (Agilent Technologies). Libraries were sequenced with a minimum of 350 million paired-end reads according to parameters described in the manufacturer’s protocol.

Our full reference was supplemented with previously collected samples, which employed an identical perturbation and sampling strategy but for which the sperm strain was reversed (male donors were an F1 hybrid of C57Bl6/J females and CAST/EiJ males). E8.5 samples included the H3K9 methyltransferase G9a and the H3K36 demethylase Kdm2b (both taken from ref ^33^). E6.5 Trim28 samples were taken from ref ^21^.

### Whole mount immunofluorescence and imaging

Whole embryo immunostaining was performed on a rotating shaker at specified temperatures and embryos were transferred using widened P200 plastic pipette tips. Briefly, embryos were fixed overnight at 4℃ (4% PFA, sterile 1x PBS) and washed 3x for 10 minutes the next day with PBST wash buffer (PBSTw, 0.2% TritonX-100, sterile 1x PBS). They were then permeabilized for at least 1 hr (PBSTr, 0.5% TritonX-100, sterile 1x PBS) and then incubated for at least 4 hours at room temperature in donkey block solution (3% BSA, 5% donkey serum, PBSTw). Embryos were then incubated in a primary antibody solution (Anti-ORFp1 Rabbit, Abcam Cat. #Ab216324; Anti-Sox2 Rat, Invitrogen Cat. #14-9811-82; Anti-Dnmt1 Rabbit, Abcam Cat. #Ab188453; Anti-Tasor, Rabbit Proteintech Cat. #30188-1-AP; Brachyury Rabbit, CST Cat #81694S, H3K27me3 Rabbit, Thermo Fisher Cat.#MA5-11198, donkey block solution) for 2 nights at 4℃. After the two nights, they were washed with PBSTw 3x for 10 min and incubated in a secondary antibody solution with 1ug/ml of DAPI (Anti-Rabbit AF555, Invitrogen Cat. #A-31572; Anti-Rat AF488, Invitrogen Cat.A48269, donkey block solution) overnight in the dark at room temperature. Embryos were then washed 3x for 10 min with PBSTw and stored at 4℃ to image within a week. The remaining PBSTw was removed and quickly replaced with Ce3D clearing mounting medium (86% Histodenz dissolved in 40% N-methylacetamide/sterile 1x PBS, sterile 1x PBS, 0.1% Triton X-100, 0.5% 1-thioglycerol) for imaging. Taser mutant embryos and time-matched controls were embedded post-staining and cryosectioned prior to imaging to confirm lack of positive signal within the nuclei.

### Imaging

All brightfield images were performed using a Leica DMi8 microscope system with a 10x air objective. All confocal images were performed using a Leica STELLARIS 5 microscope with either a 10x air objective or 40x oil objective. Images were taken with the appropriate laser and filters for Alexa 405, Alexa 488, Alexa 555, Alexa 546, and Alexa 647 and processed using Fiji ImageJ2 software (v2.9.0/1.53t).

### Whole Genome Bisulfite Sequencing

Nsd1 and Tasor mutant embryos were isolated from surrogate female mice in a petri dish with 1x PBS. Embryos were then transferred into 200 *μ*l drops of 0.2% BSA/PBS, further cleaned of Reichart’s membrane and the ectoplacental cone and microdissected using pulled glass capillaries to isolate the extraembryonic and epiblast regions independently of one another by gross morphology. For Tasor mutant data, least 5 matched epiblast tissues were collected and pooled into independent 200 *μ*l drops of 0.2% BSA/PBS to avoid cross-contamination of cells and each was incubated in a dissociation medium comprised of 2.5% pancreatin and 0.5% trypsin dissolved in PBS65 for 10 min at 4°C, followed by removal of the primitive endoderm using a narrow glass capillary in KSOM. The samples were transferred into LoBind Eppendorf tubes, snap frozen, and stored in -80°C until time for library preparation. For Nsd1, mutant embryos were isolated at E8.5, dissected independently, and submitted to individual WGBS library preparation.

Genomic DNA was extracted by incubating cells overnight at 55°C in 200 *μ*l of lysis buffer (lysis solution, 10mM Tris-HCL at pH 8.0, 10 mM NaCl, 10 mM EDTA, 0.5% SDS; plus 15 *μ*l of 20 mg/ml proteinase K per 985 *μ*l of lysis solution, Cat. #P8107S, NEB). The following day overnight digestions were spun down and 200 *μ*l of phenol chloroform was added, and samples were shaken prior to centrifugation for 5 min at 10,000 rpm at 4°C. The aqueous layer was transferred into new LoBind Eppendorf tubes and ethanol precipitation began by adding 10 *μ*l of 5M NaCl and 2 *μ*l of glycogen (Cat. #10901393001, Roche Life Science Products) to each sample and vortexed. After samples were vortexed, 500 *μ*l of cold 100% ethanol was added to and incubated for 1 hr in -80°C followed by 1 hr in -20°C. Ethanol precipitation was then performed by centrifuging 2x for 45 min at max speed (15,060 rpm) at 4°C with a 1 ml of cold 70% ethanol wash. Samples were then resuspended in 21 *μ*l of elution buffer (Qiagen). DNA concentrations were measured using Qubit Fluorometric Quantification, normalized to 0.5-5ng/*μ*l across samples, and 110 *μ*l of elution buffer was added to 20 *μ*l of sample to enable sonication using Covaris tubes (Cat. #NC9871616, Covaris Inc) and the Covaris S220 Focused-ultrasonicator with a treatment time of 1 min and 15 sec (3x with peak power 140 W, duty factor 10%, cycles per burst 200). After a second round of ethanol precipitation, samples were resuspended in 20 *μ*l of elution buffer. Bisulfite conversion and library preparation was performed the same day according to the EZ Bisulfite Conversion Methylation Gold Kit (Cat. #D5006, Zymo Research) and xGen Methyl-Seq Library Preparation (Cat. #10009824, Integrated DNA Technologies), respectively according to the manufacturer’s protocol. Libraries were then sequenced on an Illumina Novaseq 6000 or the Element Biosciences Aviti systems using 2x150 paired end sequencing.

### Computational analysis

#### General read processing and data quality filtering

A schematic diagram of the general data handling is shown in **Extended Data Fig. 1b**. Sequencing FASTQ files were first processed with 10x Genomics Cell Ranger v6.0.2^140^ for genome alignment and gene expression quantification. The unfiltered gene expression matrix was then passed to CellBender v0.3.0^141^ to remove the ambient RNA effect from burst cells in the input cell suspension for the 10x Chromium run. The output h5ad file was converted into a Seurat object with SeuratDisk v0.0.0.9021 and Seurat v5.1.0^142,143^. To remove low-quality cells, we calculated the median absolute deviation (MAD) of log-transformed UMI counts of single cells and set the minimum UMI count threshold as:

median(UMI count) - 1.5*MAD

If this threshold was lower than 2000, we used the UMI count of 2000 as the threshold. After this process, we found that the first and second GEM wells of Mphosph8 and Daxx, respectively, had a high number of low-count cells. Therefore, we removed these two libraries from the following analyses. We next sought to remove multiplet cell barcodes (more than one cell encapsulated in a single droplet during the library preparation step). We adopted Scrublet v0.2.3 ^144^, which computes the likelihood of multiplets for each cell barcode by simulating synthetic doublets in a given dataset. We set the expected doublet parameter as 7.5% and removed putative multiplet cell barcodes with parameters min_gene_variability_pctl=85 and n_prin_comps=20 for the scrub_doublets function. We inspected the doublet score distribution, and if the threshold doesn’t split the peaks of a bimodal distribution, we used a hard-coded doublet score threshold of 0.25 to extract singlet cells.

#### Embryo demultiplexing

Embryo demultiplexing was performed using the same workflow as published previously to deconvolute random CAST haplotypes^33^ with minor modifications. In brief, the cDNA reads in Read 2 were aligned to the mouse reference genome mm10 using STAR v2.5.3^105^, and the alignment BAM file was split into B6 (reference) and CAST (alternative) using SNPsplit v0.3.4^145^. We then counted the read counts assigned to two genotypes for each chromosome for a single cell, which was transformed into a cell-by-chromosome allelic read fraction matrix. Using only autosomes, we first filtered cells with at least 1,000 SNP-typable read counts. Then, we performed k-means clustering of the allele fraction matrix with k = the number of pooled embryos. For each k-means cluster, we calculated the Mahalanobis distance from each cell in the cluster to the centroid and filtered out cells that are more distant than

median(Mahalanobis distance) + MAD

to focus on confidently demultiplexed cells. To visualize the demultiplexing result, we applied UMAP^146^ to the allelic count matrix with parameters n_neighbors = 15, n_components = 2, metric = “cosine”, and min_dist = 0.1 using the R uwot package v0.1.11^147^. The successfully demultiplexed cells were regarded as high-quality, confident cells and used for downstream analyses. Note that we called embryos by assembling datasets from multiple GEM wells in case a single pool of cells was split into two wells; if each cell suspension was assigned to an independent GEM well, we applied this pipeline to sequencing data from each well.

Sex-typing of embryos was performed by computing the linear-normalized UMI counts of Xist and ChrY genes (average), and count thresholds of 1e-4 and 1e-7 were applied to the respective sex markers.

#### Cell state labelling

The WT data from multiple time points were merged, and the standard Seurat pipeline was adopted, where UMI count log-normalization, standardization, identification of the top 2000 highly variable genes, and PCA were performed. We selected the top 16 PCs for UMAP embedding and unsupervised clustering, according to the standard deviation of each PC. To identify clusters of cells with the same identities, we used iterative unsupervised clustering of the WT reference data as follows. First, we generated a preliminary set of clusters by jointly clustering the merged datasets, which allowed us to separate the major germ layers and designate low-quality cells (characterized by low UMI counts) as “shadow” clusters. Next, stage-specific clusters were derived based on this initial assignment and annotated according to the enrichment of manually curated stage-specific marker genes, as well as the developmental stage from which the samples were collected. Finally, clusters corresponding to the same biological state were merged, ensuring coherent and accurate identification of stage-specific cell populations. Cell state assignments were conducted as part of a companion manuscript based upon an extensive literature search (Wang *et al.*, in submission). A full online webtool will be provided at the time of publication to provide detailed justification for each assigned state.

#### Metacell analysis and Network Flow inference

For feature gene selection in the Metacell (MC) analysis, genes related to cell cycle and stress were first excluded by manual annotation. Following established procedures using metacell v0.3.7^39^, we identified 3,037 highly variable genes. MCs were then generated with K = 100, using standard bootstrapping, MC splitting, and outlier filtering as described by Baran et al.^39^. The resulting reference model comprised 1,206 MCs representing 90,886 cells, after excluding outlier cells from the analysis.

MC annotation was performed using Seurat’s label transfer function, with minor manual adjustments based on the predominant cell type within each MC. To evaluate robustness, we followed the strategy employed by Mittnenzweig et al.^27^. Specifically, we generated 72 replica MC sets, each time leaving out cells from one of the 72 reference embryos. For each replica, MC partitions, manifold distances, and flows were recomputed using the same parameters as in the reference analysis. MCs were annotated by projecting each replica MC to the most correlated original MC, based on log-transformed mean gene UMI fractions of the feature genes. Intrinsic embryo ranks and age-group partitioning were preserved from the representative MC sets. Finally, the network flow was aggregated across the 72 replicas.

For the unidirectional graph representation in **Fig. 1b**, we summed up all the MC-by-MC transition score matrices between age groups to obtain the overall MC transition weights. We then subtracted the weights of edges connecting the same pairs of nodes but with opposite directions and retained edges with a minimum delta-weight of 5.

#### Computational sorting of developmental stages of WT embryos

In this study, we adapted the embryo ordering algorithms of Mittnenzweig et al.^27^ with modifications. We inferred the intrinsic temporal ordering of embryos by analyzing embryo–embryo similarities and optimizing a linear ordering that minimized the temporal distance between similar embryos. To avoid potential biases from unstable sampling of extraembryonic tissues and the hematopoietic lineage, we removed all residual cells corresponding to the extraembryonic endoderm, extraembryonic ectoderm, and hematopoietic lineage prior to further analysis. Following this filtering, we aggregated the unique molecular identifier (UMI) matrix by embryo and performed principal component analysis (PCA) using the AggregateExpression and RunPCA functions in the Seurat package. Embryo–embryo distance was then defined as the Euclidean distance in PCA space, and similarity as the inverse of this distance. For ordering, we implemented an algorithm resembling the bubble-sort technique, iteratively adjusting the positions of adjacent embryos whenever doing so reduced the goal function, continuing until a local minimum was reached. After optimization, age group assignments were manually defined, ensuring that each group contained at least three embryos.

#### CRISPR cut site read counting

For quantifying reads overlapping CRISPR-target cut sites, we extracted reads mapped through the cut site of each sgRNA, which is between 3- and 4-bp upstream of the protospacer adjacent motif (PAM), from an embryo-level BAM file produced by CellRanger. Using CIGAR strings, we went over each read by nucleotide and saw if the cut site overlapped with ‘M’ (Match), ‘I’ (Insertion), ‘D’ (Deletion), ‘S’ (Soft clipping), or ‘N’ (Skipped region). Given that the majority of CRISPR-Cas9 edit outcomes are deletions, we counted the number of reads with ‘M’ mapping at both the third and fourth base pairs from the PAM as the unedited read count. The unedited read count was measured for all 114 sgRNA cut sites across 33 target genes and normalized by the total read count for each embryo. Embryos with less than 500 total reads supporting the cut sites were filtered out. We applied the same workflow to the bisulfite sequencing data using deduplicated BAM files using BSMAP^148^.

#### Reference embedding of mutant cells

To assign cell states to the regulator mutant datasets, we applied reference-based projection rather than *de novo* cell state identification. For each query mutant dataset, we first identified shared anchor points by Seurat’s FindTransferAnchors using the top 30 PCs of the WT reference data. This anchor data was passed to the MapQuery function to embed the mutant cells onto the reference PC space, followed by dimension reduction by UMAP. This also allows for assigning a cell state to every single cell, and we applied the transfer score threshold of 0.5 to filter out cells with ambiguous cell identity. To infer the transcriptional ranks of mutant single cells, we collected the top 20 nearest-neighbouring WT cells for each mutant cell and took the median rank.

#### Cell state proportion-based rank estimation

Per-replicate cell state proportion data including WT and mutant embryos (50 >= cells per embryo) were converted into centered log-ratio (CLR), which is a log-ratio of a specific cell state fraction to the same embryo’s geometric mean fraction. The CLR matrix was embedded in the PC space. To order the WT embryos by the developmental progression levels, we used two embryos as anchors, the earliest one at E6.5 and the latest at E9.0, predicted by the transcriptional rank approach described above. Using these start and end embryos, we ordered the remaining 70 embryos by connecting them through the shortest path within the PC space using the R library TSP^149^. We then computed the nearest five WT embryos within the PC space from each mutant embryo and took the median rank as the mutant’s estimated rank. The distance to the centroid was taken among replicates in the PC space to represent the phenotypic variability.

#### Non-negative matrix factorization (NMF)

To streamline NMF analysis, we first created pseudobulk data for cell state-by-embryo replicate combinations for each condition, with a minimum threshold of 10 cells. Since we could not sample a large number of PGCs, we merged PGCs by condition without considering the embryo replicates. We then performed log-normalization of the UMI counts.

To focus on highly informative genes dictating sample conditions, we selected the top 500 highly variable genes across pseudobulk cells within the same cell state, and iterated this analysis over the 68 reference cell states. This resulted in 8,487 union genes, with 508 genes overlapping among at least 20% of the cell state comparisons. At the same time, we obtained the top 2,000 highly variable genes in this pseudobulk data to recover genes that are variable regardless of the sample conditions. We took the union of these gene sets, which resulted in 2,261 genes as the NMF-target gene set. For the NMF of REs, we followed the same criteria with minor modifications: we filtered out pseudobulk cells with fewer than 1000 UMIs, and we selected the NMF-target RE family set by taking the union of the top 100 within-state highly variable RE families (HVREs) and the top 100 global HVREs, resulting in total 141 RE families.

For gene-level NMF, we first performed a coarse sweep by setting the number of programs (k) to 20, 40, 60, 80, and 100 with triplicates using the crossValidate function of the RcppML package v0.3.7^150^. We then performed an extensive sweep by setting k from 40 to 70 and decided to use k=55 as the representative number of programs (see **Extended Data Fig. 8** for validation analyses).

For REs, we first swept parameters by k = 5, 10, 15, 20, 25, and 30, and then performed a fine sweep of k from 2 to 15, where we took k = 9 as the number of programs (see **Extended Data Fig. 13** for validation analyses). To compute the GEP usage in Dnmt1-/- embryos from Clark *et al.*^32^, Tasor/Dnmt1 double mutant, and time-course Tasor KO embryos, we used the pre-built NMF model as a reference and applied the project_model function to a combined pseudobulk gene expression matrix of all embryos, including double mutants, time-course mutants, and all the other conditions.

High contribution Gene/RE sets were determined for each NMF program by taking a cumulative contribution score threshold of 0.9 (the top contributor genes/REs making up 90% of the total contribution score for the program). The functional enrichment of the NMF program marker gene sets was assessed by Enrichr^151^ through GSEApy v1.1.8^152^.

#### Differential gene expression test

To obtain differentially expressed genes (DEGs) in mutants, we used the cell state-by-embryo replicate pseudobulk expression data with a minimum number of cells per pseudobulk of 10. For each cell state, gene expression comparison was performed using the Seurat’s FindMarkers function with the DESeq2^153^ wrapper implementation if the number of embryos in KO and WT conditions were both at least four. Adjusted *P*-value threshold of 0.01 was applied to the result, and DEGs with Log2FC >= 1 were used to construct the regulator network presented in **Extended Data Fig. 11b, c.**

#### Bisulfite sequencing analysis

Raw reads from our Tasor and Nsd1 mutants were subjected to adapter and quality trimming using cutadapt (version 4.4; parameters: --quality-cutoff 20 --overlap 5 --minimum-length 25; Illumina TruSeq adapter ‘AGATCGGAAGAGC’ clipped from both reads), followed by trimming of 10 and 5 nt from the 5’ and 3’ end of read one and 15 and 5 nt from the 5’ and 3’ end of read two. Trimmed reads were aligned to the mouse genome (mm10) using BSMAP^148^ (version 2.90; parameters: -v 0.1 -s 16 -q 20 -w 100 -S 1 -u -R). Duplicates were removed using the ‘MarkDuplicates’ command from GATK (version 4.6.2.0; -- VALIDATION_STRINGENCY=LENIENT --REMOVE_DUPLICATES=true). Methylation rates were called using mcall from the MOABS^154^ package (version 1.3.9.6; default parameters). Only CpGs covered by at least 10 and at most 150 reads were considered for downstream analyses.

We assembled the data with the published data of epiblast tissue (E6.5 for all other mutants and WT^33,98^). We defined sequences up/downstream of 2 Kbp from the transcriptional start sites as promoters, and we collected promoter sequences of the GEP8/29/37 gene set. CpG methylation profiles of the promoter regions were analyzed for each mutant.

#### RE expression quantification

Given the multiple configurations of REs in the host genome (**Extended Data Fig. 13a**), we prepared two versions of RE annotation GTF files (mm10). The first reference file records REs if they are intergenic or inserted into host genes in the discordant direction according to the DFAM v3.8 database^155^. The second records the whole set of REs regardless of the inserted loci, and unique suffixes were added to all the RE copy names to enable independent quantification of each individual RE copy. In both files, we set the same start and end coordinates for gene, transcript, and exon entries for an RE copy.

For the comparisons of RE expression counts between STARsolo and scTE v1.0^156^ or IRescue v1.2.0b1^157^, we used STAR v2.7.11b with the following options: “--outFilterMultimapNmax 100 -- winAnchorMultimapNmax 100 --outMultimapperOrder Random --runRNGseed 1 --soloStrand Unstranded --soloMultiMappers EM.” Since neither scTE nor IRescue considers the strand directions of REs, this option set allows for fair comparison among software by distributing multi-mapped reads using the EM algorithm used in the RE quantification field^158^.

For the family-based analysis used in **Fig. 4**, **5**, and **Extended Data Fig. 13**, we used the first filtered RE annotation file as the reference and used STAR with an option set “--outFilterMultimapNmax 1 -- soloStrand Forward”, so that we consider only uniquely mapped, autonomous RE transcriptions.

For single-copy RE quantification, we used the second locus-aware RE reference file with the same STAR option above for the sense-count quantification, and for antisense-counting, we used “--soloStrand Forward” instead. To reduce the size of the unique RE copy expression matrix, we pre-filtered RE copies with fewer than 50 UMI counts for each condition before merging. For the dimensional reduction of the single-copy expression data, we used the same Seurat pipeline described above, and we used the top 14 PCs for UMAP embedding and unsupervised clustering.

For inspecting the read mapping onto the reference genome, we first converted BAM files created by CellRanger into BigWig files by bamCoverage in deepTools v3.5.5^159^ with an option set “--binSize 25 -- normalizeUsing CPM.” Then, pyGenomeTracks v3.9^160^ was used for visualization.

#### Co-expression analysis of REs and proximal genes

In the sense-counting single RE copy expression data, we extracted differentially expressed RE copies (DEREs) for the unsupervised clusters by Seurat’s FindAllMarkers function with the options “only.pos = T, logfc.threshold = 1, return.thresh = 1e-2”, and further filtered out DEREs with adjusted *P*-value > 1e-10. For each DERE, upstream and downstream 100kbp regions were extracted, and genes were assigned to 10kbp bins if their transcription start sites fell in this range. Then, gene expression Z-scores of the neighboring genes were assessed across bins and sample conditions.

#### Construction of cell state lineage tree

To summarise the MC-level network flow graph as a cell state transition graph, we first aggregated the MC nodes into cell state nodes by summing up transition weights, followed by weight normalization with the number of MCs per source cell state. This cell state transition graph was converted into a unidirectional graph by subtracting the normalized weights of bidirectional edges, retaining only positively weighted edges.

To further summarise the cell state transition graph as a cell lineage trajectory tree, we excluded extraembryonic, blood/vascular, germ, and axial mesoderm cell states due to the graph disconnectivity and/or undersampling of cells within the reference dataset. We then selected cell states with at least 100 cells at WT E8.5 as terminal nodes of the lineage tree, and constructed the tree by taking the shortest paths from the undifferentiated epiblast state to those states. We manually inspected the trajectories and removed paths towards the neural crest and lateral plate mesoderm limb states due to inconsistency with the literature.

#### Identification of dynamically regulated gene sets in a developmental path

For each trajectory path of the lineage tree, we extracted state-by-replicate pseudobulk cells included in the path from the E6.5-8.5 reference data. Highly variable genes along the path were selected by a Z-score standard deviation threshold of 1. We then performed K-means clustering (k=4) for the gene expression Z-score matrix to cluster the expression pattern along with the path. To identify clusters regulated coordinately, Pearson’s correlation of gene expression profiles was computed among genes within a cluster, and we selected clusters with a minimum correlation coefficient of 0.3. As a substantial amount of gene overlap is expected between clusters, we computed Jaccard indices of cluster pairs and took the unions within hierarchical clusters, defining the union sets as dynamically regulated gene sets.

#### RNA velocity

We used velocyto v0.17.17^121^ to quantify exon and intron UMI count separately, and the spliced and unspliced UMI count matrices were processed by scanpy v1.10.2^161^ and scVelo v0.3.2^162^. The WT and Tasor time course embeddings were both obtained by the abovementioned Seurat pipeline, in which we used the top 13 PCs for the Tasor embedding. In the dynamic modelling of the RNA velocity, we used the top 15 PCs for computing nearest neighbors. The other steps were performed with the default parameters.

### Statistical analysis

Box plots represent the median, 25th and 75th percentiles, and 1.5x interquartile ranges (whiskers). Outliers were excluded for the visualization purpose, but we included all the data points for statistical tests. We selected the two-sided (or one-sided if there is a clear expectation in direction) Wilcoxon rank-sum test since we do not assume a normality in data distribution. FDR adjustment was applied when multiple tests were conducted.

## Code availability

All the codes we used for analyses will be available on GitHub and Zenodo upon publication.

## Data availability

We reanalyzed our published mutant scRNA-seq datasets for Kdm2b, G9a, and Trim28 (GEO: GSE137337 and GSE159468) and developmental scRNA-seq reads from E1.5 to E7.5 (GSA: CRA003171; Cao et al., 2026^115^). The processed scRNA-seq data for the testis development^163^ was downloaded from ArrayExpress (Accession ID: E-MTAB-6946). All the other single-cell raw sequencing data and processed data will be available upon publication.

---

**Supplemental Table 1. Historical descriptions of regulator mutants in mice.**

Collection of previously reported mouse phenotypes for selected genes, with an emphasis on functionally validated null alleles rather than clinical or dominant negative mutants (which are particularly enriched for regulators such as Lmna). Reported lethalities reflect the limited ranges offered by traditional approaches. Alongside these historical reports, we include the number of embryos sampled in this study, their mean inferred transcriptional rank, and mean inferred knockout efficiency as estimated from read-level analysis of demultiplexed replicates (see **Extended Data Fig. 2** and **Methods**).

**Supplementary Table 2. sgRNA sequences for high efficiency zygotic gene disruption**

Protospacer sequences and genomic coordinates for the 2-5 sgRNA’s selected to disrupt each target gene. In most cases, sgRNA design was tailored to minimize off target effects and be maximally distributed across the coding sequence.

**Supplementary Table 3. Differentially expressed genes in mutants versus WT**

Lists of differentially expressed genes (DEGs) in each mutant versus the same cell type in WT. Statistical tests were performed using DESeq2 for replicate-resolved per-cell type pseudocount data.

**Supplementary Table 4. Gene Ontology (GO) term enrichment in GEP contributor genes**

Results of GO Biological Process term enrichment analysis for each NMF program using Enrichr^168^.

**Extended Data Figure 1.**
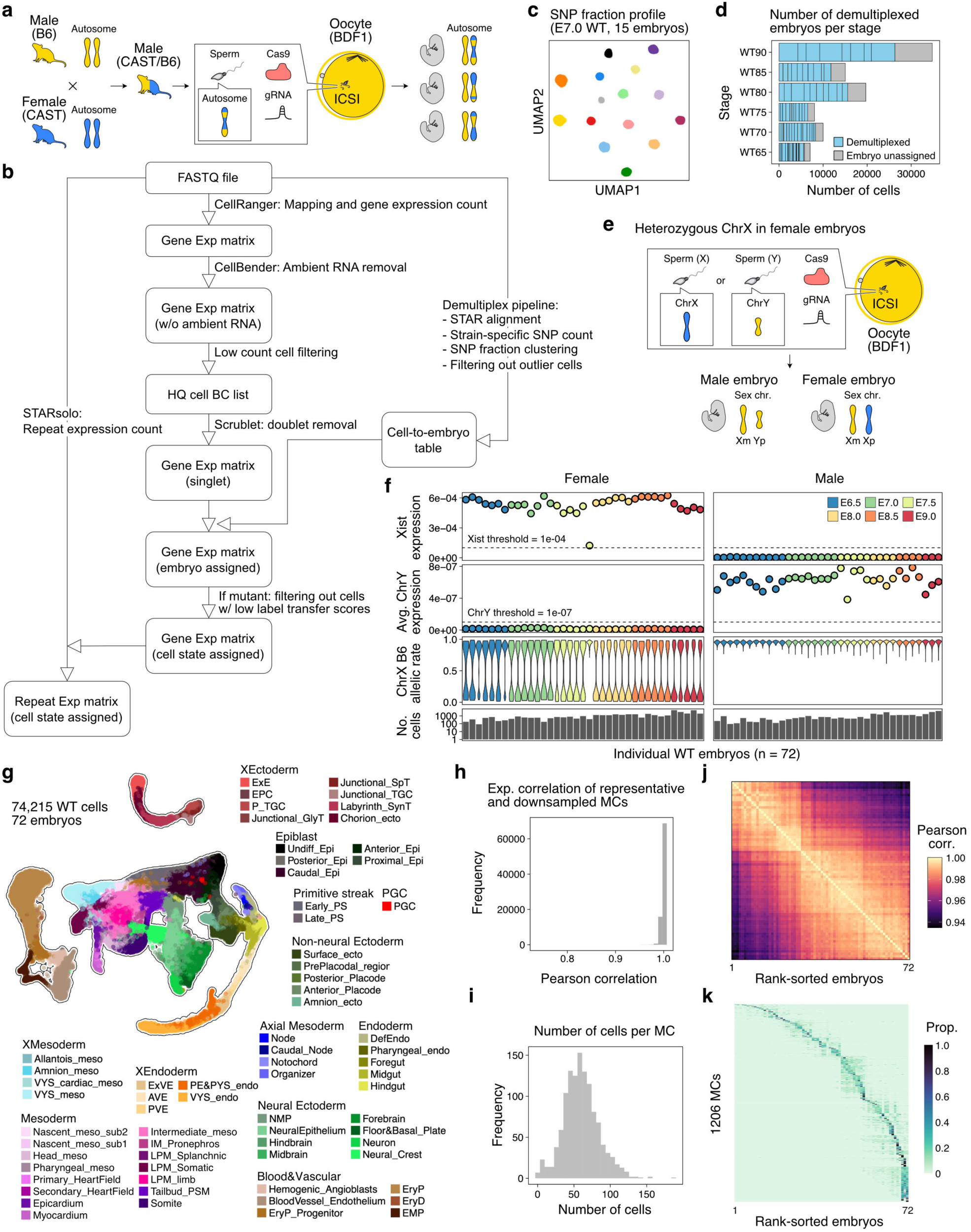
Deconvolution and cell state assignment strategy for assessing mutant embryo cohorts. **a.** Schematic of embryo generation strategy using CAST/B6 F1 sperm to generate SNP-typable embryo cohorts. Sperm donors are derived from a CAST/EiJ female crossed to a C57BL6/J male, such that each replicate embryo receives a random CAST haplotype across its 19 autosomes. Zygotes are generated from a limited number of donor males using the same cross using Intracytoplasmic sperm injection (ICSI), followed by delivery of Cas9 and 2-5 single guide RNAs (sgRNAs) distributed throughout the coding sequence of the target locus or loci. Surviving blastocysts are transferred into pseudopregnant females and recovered at the time point of interest (generally E8.5), after which cohorts of ∼10 embryos are profiled as a clutch via single-cell RNA-sequencing (scRNA-seq) using the 10x Chromium Platform. **b.** Overview of the computational pipeline used in this study, including key steps in embryo demultiplexing by genotype (CAST SNP profile), correction for ambient RNA, and the assembly of cell state resolved gene- and repeat-level transcriptomes. **c.** Representative UMAP of SNP-typable data, taken from our control E7.0 clutch of 15 pooled embryos. Randomly-transmitted CAST/EiJ SNP profiles provide a unique embryo barcode that reflects the transmission of grandparental information from the male donor. **d.** Stacked barplot highlighting the number of cells recovered and confidently assigned to each embryo of our reference manifold, as well as the number of cells that are confidently assigned to a cell state but do not pass our criteria for replicate-level assignment. **e.** Strategy for resolving allelic expression of ChrX in female embryos. Briefly, our crossing method is designed such that all female embryos receive their paternal grandmother’s CAST/EiJ ChrX, allowing high-resolution determination of ChrX silencing behavior and transcriptional dynamics. **f.** Sex typing information for deconvolved female and male replicate embryos, including for Xist expression (top), Average expression of ChrY transcripts (middle), and per cell ChrX allelic ratios (bottom). Out of 36 recovered “female embryos,” all but one show high Xist expression and bimodal ChrX expression. **g.** UMAP embedding for our control reference time series, including assignments into one of 68 transcriptional states. **h.** Distribution of gene expression correlations between representative metacells and their proximal bootstrap metacell samples, as previously described^27^. Bootstrap samples were prepared by the leave-one-out strategy among the 72 embryos, and the sharp peak around the correlation value = 1 indicates that the metacell definition does not depend on specific individual embryos. **i.** Distribution of the number of cells in each metacell for our network flow model. **j.** Bulk-level transcriptome correlation among temporally sorted WT embryos, showcasing the gradual transition from the early through the late transcriptional ranks. **k.** Proportion of metacell occurrence across embryos, which indicates the temporal transitions of cell states throughout development.

**Extended Data Figure 2.**
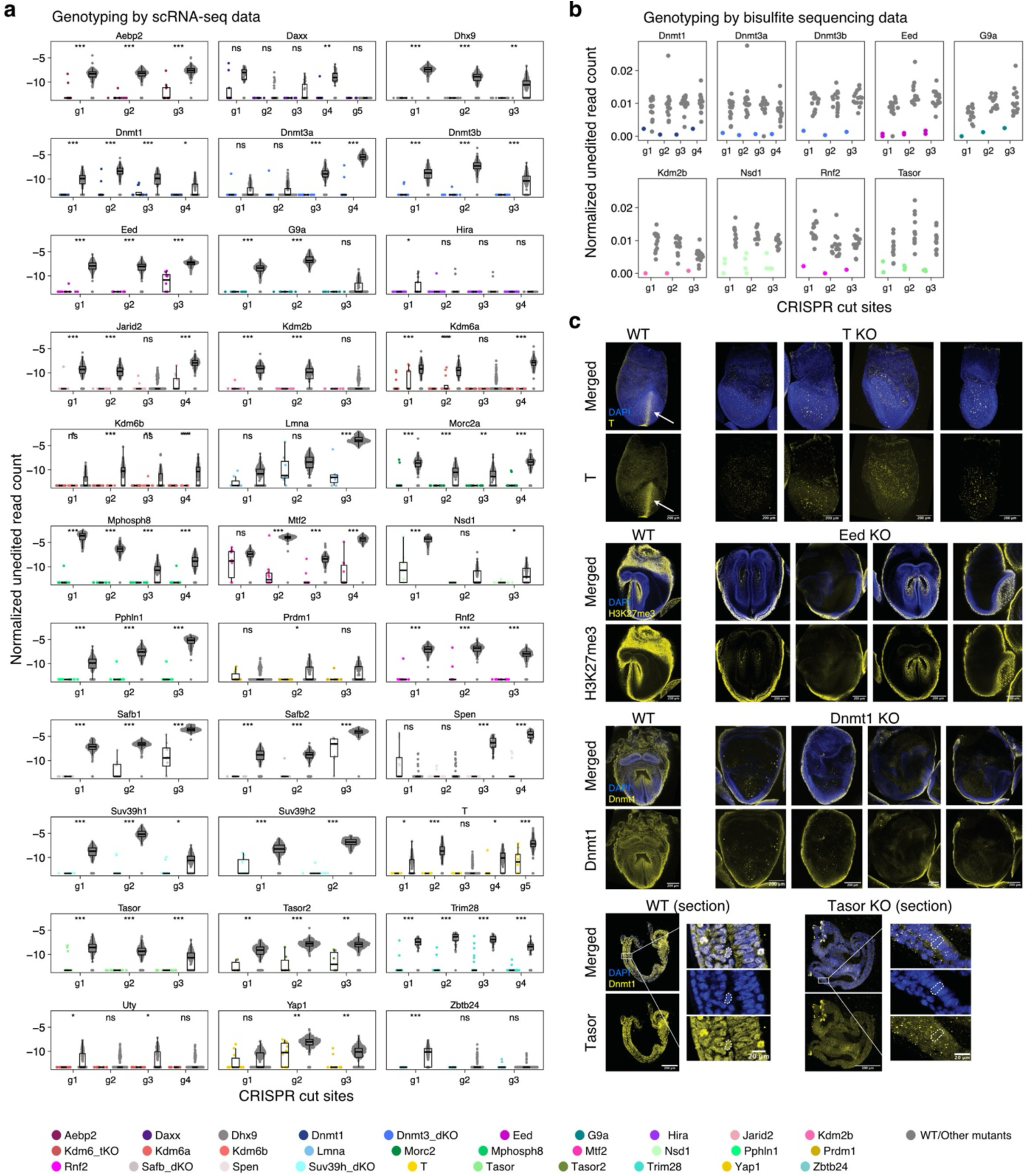
Validation of Crispant-based zygotic disruption of nuclear regulators. **a.** Relative recovery of unedited reads from each demultiplexed embryo across each co-injected sgRNA. Briefly, our Crispant method acts by simultaneously injecting 2-5 sgRNAs targeted to the locus of interest and designed to cover as much of the coding sequence as possible. This leads to multiple distinct editing outcomes, including indels and large truncations, which often trigger non-sense mediated decay. To summarize the cumulative effect of these outcomes, we calculated the normalized expression of each sgRNA’s predicted cut-site, which represents the remainder of unedited transcripts within each embryo. Recovery data from the on-target sample (colored dots in each panel) are compared to those from off-target samples (other mutant embryos and the WT time series data; grey dots) collected at the matching time point. Overall, very few embryos showed any evidence of residual unedited transcripts, with mean inferred edited rates of 99.4% (SD = 7.2%). Statistical tests were performed by one-sided Wilcoxon’s rank-sum test followed by FDR adjustment. ns, not significant; \**P* <= 0.05; \*\**P* <= 0.001; \*\*\**P* <= 0.0001. **b.** Same as in a but for Whole Genome Bisulfite Sequencing (WGBS) data presented in Figure 3. Here, percent editing rates can be slightly inflated due to the imperfect nature of alignment to bisulfite converted reads. Nevertheless, the inferred unedited rates remain very low, with mean inferred edit rates of 99.4% (SD = 2.1%). Statistical tests were performed by one-sided Wilcoxon’s rank-sum test followed by FDR adjustment. ns, not significant; \**P* <= 0.05; \*\**P* <= 0.001; \*\*\**P* <= 0.0001. **c.** Functional validation of functional knockout by Immunofluorescence, shown for four high interest candidates (Brachyury T, Eed, Dnmt1, and Tasor). For Brachyury T, embryos were collected at E7.5 to examine the loss of localized signal within the primitive streak (white arrow in WT). For Eed, we examined the presence of the target modification H3K27me3. Whole mount IF shows little to no evidence of functional mosaicism, with signal in the outer yolk sac and inner definitive endoderm likely reflecting antibody cross-reactivity from whole mount staining. Finally, for Tasor, E7.5 embryos were sectioned for higher resolution analysis, again confirming embryo-wide lack of nuclear signal in mutant cohorts. For each mutant, at least five mutant embryos were selected and showed identical results.

**Extended Data Figure 3.**
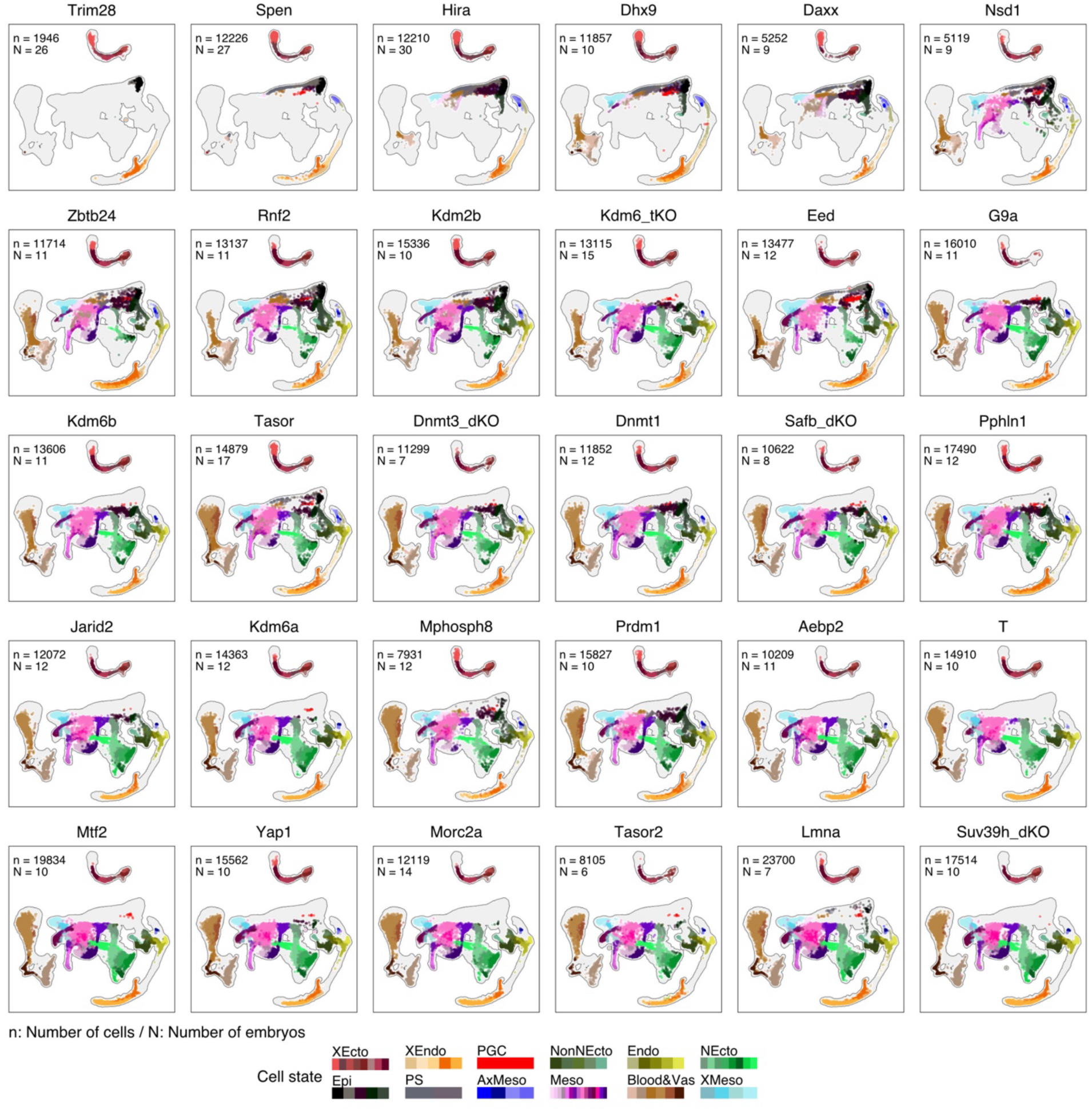
A multi-regulator dependency map of mouse gastrulation and organogenesis. Projection of mutant cells onto the reference UMAP, with the number of recovered cells (n) and embryo replicates (N) highlighted per target regulator. Samples are listed from left to right in the order of the median transcriptional rank for embryonic cells (excluding the placenta and yolk sac). Cohorts such as Trim28, Spen, Hira, Daxx, or Dhx9 show very minimal occupancy of the reference time course, and arrest within the pluripotent epiblast or early primitive streak (cell states within the top right of the UMAP). Alternatively, regulator mutants such as Lmna, Suv39h1/2, or Morc2a occupy more advanced states for each germ layer present within the bottom left.

**Extended Data Figure 4.**
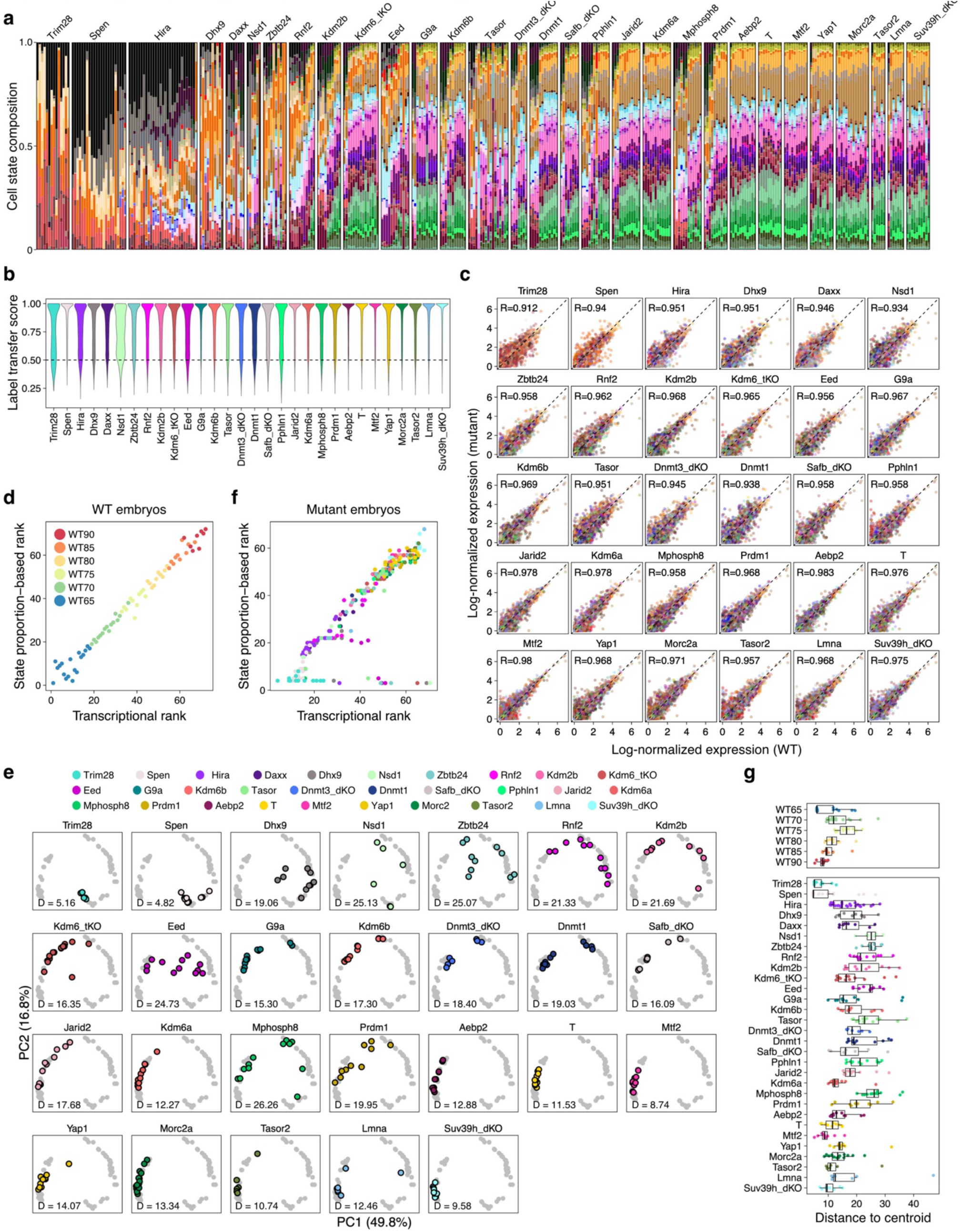
Nuclear regulator mutants are associated with increased phenotypic variation. **a.** Stacked bar plots for each mutant regulator cohort, ranked from left to right according to increasing transcriptional rank (median, calculated for the embryonic lineages). The most severe mutant phenotypes show minimal generation of formal germ layers and tend to arrest within the pluripotent epiblast or early primitive streak. Purples and greens highlight the induction of embryonic mesoderm and neural ectoderm, respectively, and are only apparent in mutants that progress through stages of early body axis patterning. **b.** Per cell label transfer prediction scores, as inferred by Seurat, for each of our regulator mutant data sets according to transcriptional states defined by the control reference. In general, cells from nuclear regulator mutants are confidently assigned, indicating that transcriptional dysregulation is rarely great enough to override a cell’s genetically encoded regulatory network. The dashed line represents a transfer threshold of 0.5, and all cells with transfer scores below this value were excluded from all analyses. **c.** State-level normalized gene expression scatterplots for the top 2,000 highly variable genes used to inform the assembly of our reference data set. Each dot represents a single gene expression value for a cell state. Pearson’s correlation coefficient is also included in the inset. Overall, marker gene level correlations are high across nuclear regulator mutants, even in scenarios where morphology is broadly disrupted. **d.** Comparison of two different estimators for developmental rank, including based on the relative transcriptional embedding of each cell (“transcriptional rank”, based on the median inferred rank for each cell based on its 20 nearest neighbors in the KNN graph, see **Methods**) and on the relative composition of individual cell states (“Cell state proportion rank”, generated on the PCA embeddings of state proportion matrix, see **Methods**). Briefly, our control embryos were ordered from 1-72 based upon these two criteria and show high correlation between one another. **e.** Projection of mutant cohorts onto the WT state proportion PCA plot often show consistent grouping of the majority of replicates within the same general time window, with areas of higher variation frequently seen for regulators with severe effects. D refers to the mean distance to centroid for the cohort. **f.** Comparison of transcriptional rank and state proportion-based rank show very high concordance. Deviations from the diagonal are observed for Trim28 (light blue), which is compositionally heterogeneous but composed of only early cell states (Epiblast, Extraembryonic Ectoderm, and early VE states), as well Eed (purple), which shows a notable posteriorization phenotype. **g.** Per embryo replicate distance to centroid measurements for our complete cohort, including WT controls highlight the higher variation in Crispant mutants, particularly those with obvious developmental defects. Note that many with normal or expected transcriptional ranks also tend to have lower overall variation.

**Extended Data Figure 5.**
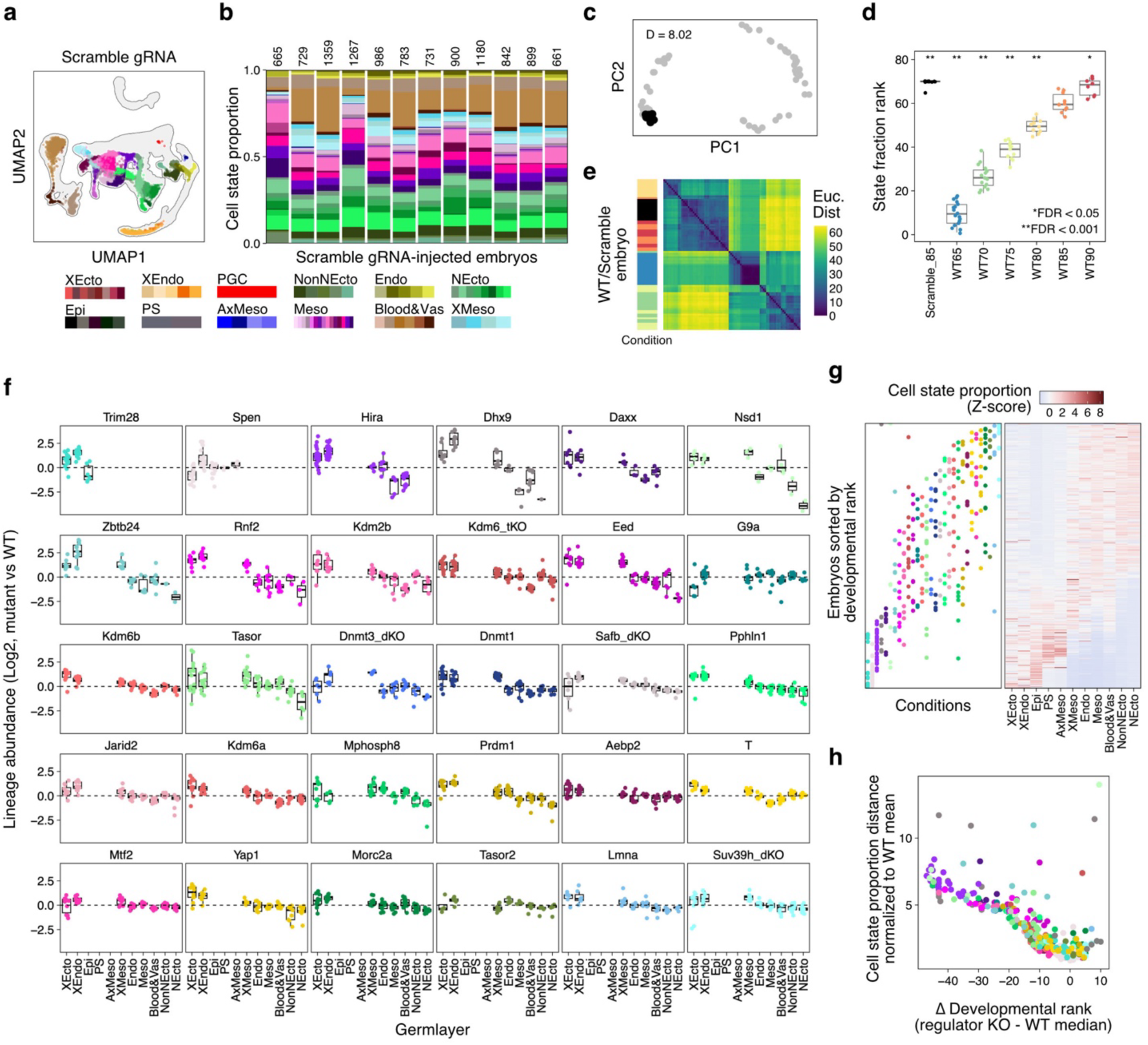
Epigenetic regulator mutants are generally depleted of neural ectodermal cell states. **a.** UMAP projection of a scrambled guide control cohort consisting of 12 E8.5 embryos onto the WT reference shows expected occupancy of late gastrulation and early organogenesis states. Note that for this control experiment, extraembryonic tissues were removed to increase the precision of embryonic progression, leaving only some residual yolk sac cells (orange). **b.** Per replicate compositional bar plot of individual scrambled guide control samples, with number of cells sampled per replicate highlighted. **c.** Projection of scrambled guide embryos onto the control PCA to infer variation by cell state composition, with the distance to centroid indicated as for the mutant embryos in **Extended Data Fig. 4e**. **d.** Mean cell state proportion rank estimate for the scrambled guide control cohort, statistically tested against each WT time point. The control cohort deviates significantly from each timepoint, but is situated between E8.5 and E9.0, validating the effects of select mutations on developmental progression. Two-sided Wilcoxon’s rank-sum test followed by FDR adjustment: ns, not significant; \**P* <= 0.05; \*\**P* <= 0.001. **e.** Clustering of embryo replicates by Euclidean distance by cell state composition. Scrambled sgRNA controls (black) cluster with E8.5 and E9.0 samples, confirming that the injection approach itself has minimal effect on embryo progression. **f.** Per replicate compositional differences compared to time-matched reference data. For each regulator mutant, the Log2-fold change is calculated for each lineage in comparison to the time-matched control samples. Samples are ranked from most to least severe according to transcriptional rank scores as in Fig. 1e. Mutant embryos often show an overall increase in the proportions of XEcto, XEndo, and XMeso cell states and a decrease in the proportions of NEcto cell states, although this is particularly pronounced for the core Polycomb subunits Rnf2 and Eed, as previously reported by Grosswendt et al^33^. Overall, cell state proportions also tend to be more variable between individual replicate embryos, particularly for those with severe phenotypes. **g.** Left: Dot plot highlighting the aggregate transcriptional rank for each embryo cohort, sorted by the severity of the mean phenotype (with Trim28 on the left and Suv39h1/2 on the right). Color scale is as in **Extended Data** Fig. 5f. Right: Z-score-transformed lineage abundance for each mutant embryo. Individual nuclear regulator mutants show a broad dynamic range in terms of their embryonic progression, and tend to reside during the window immediately following the induction of early primitive streak products (PS, Axial Mesendoderm and Extraembryonic Mesoderm, respectively) and prior to the induction of Neural Ectoderm. **h.** Per-replicate scatterplot showing the relationship between the difference in transcriptional rank (x axis) and composition-based Euclidean distance (y axis) for each regulator mutant replicate highlights the degree to which phenotypic severity (at the level of developmental progression) corresponds with higher degrees of phenotypic variation (at the level of replicate-level cell state composition).

**Extended Data Figure 6.**
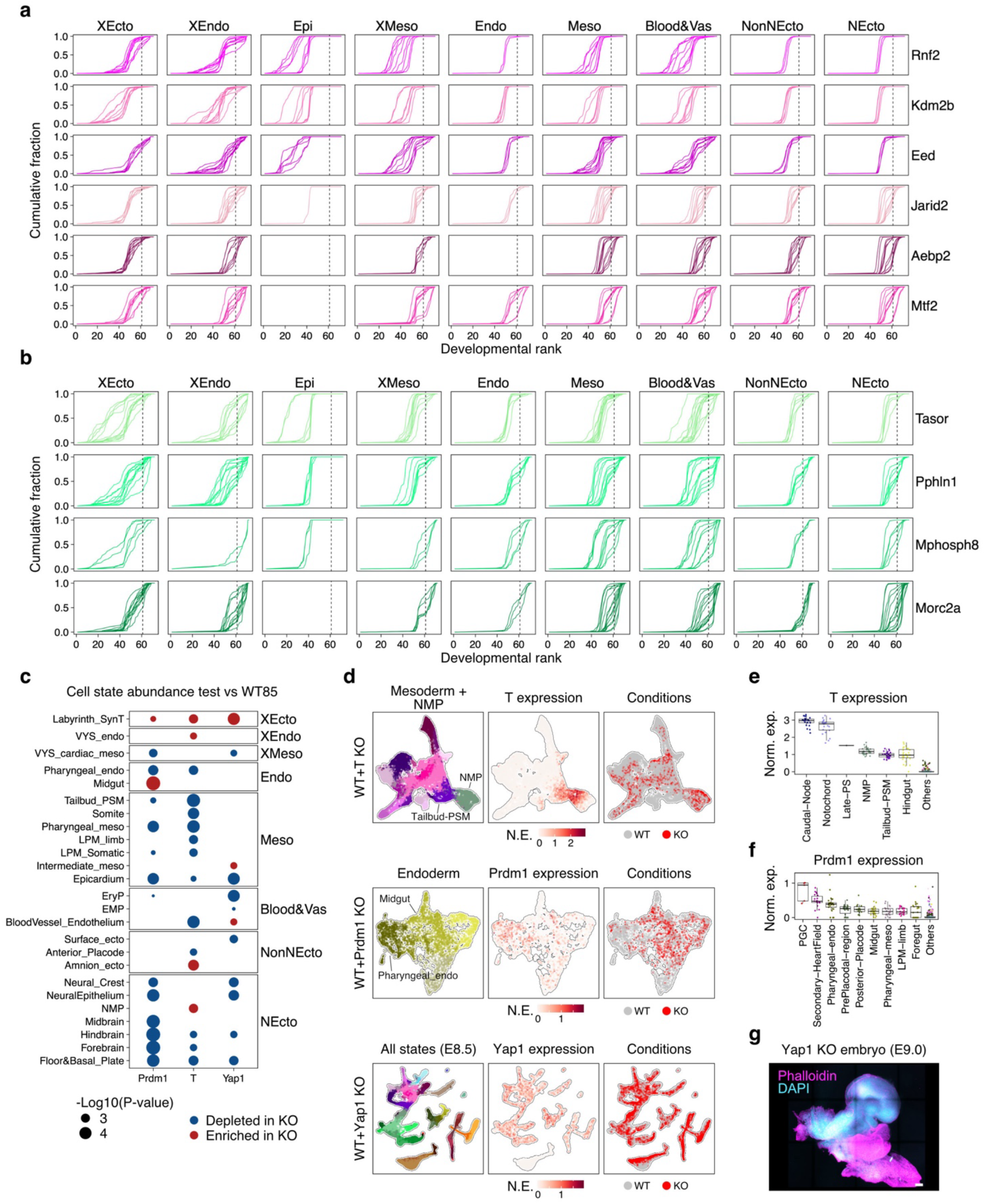
Mutant phenotypes stratify regulators into primary and secondary roles. **a.** Per replicate lineage level cumulative distribution functions (CDFs) of transcriptional rank for Polycomb associated factors, including Rnf2, Kdm2b, Eed, Jarid2, Aebp2, and Mtf2. Regulator mutants are ranked top to bottom from most to least delayed. The dashed vertical line highlights the mean transcriptional rank of E8.5 reference embryos. Rnf2, Kdm2b, and Eed retain pluripotent epiblast states and exhibit greater delays in embryonic progression compared to Jarid2, Aebp2, and Mtf2. **b.** CDFs of transcriptional rank for HUSH-associated factors, including Tasor, Pphln1, Mphosph8 and Morc2a. The dashed vertical line highlights the mean transcriptional rank of E8.5 reference embryos. Tasor is the most delayed, followed by Pphln1 and Mphosph8, while Morc2a replicates tend to be distributed closer to the reference. **c.** Transcription factor (TF) mutations cause specific developmental phenotypes. To test the relative enrichment or depletion of individual cell states, we compared the per-replicate cell state abundance in the three TF conditions to control reference data. *P*-values were calculated by the two-sided Wilcoxon rank sum test, followed by FDR adjustment. Prdm1 mutant regulators exhibit expected depletion of pharyngeal cell types, including the Pharyngeal mesoderm and endoderm, as well as broader depletion of neural cell types that could reflect developmental delay^80^. In contrast, T mutant embryos are depleted of Tailbud Pre-Somitic Mesoderm, Somites, Lateral Plate (LPM) and other mesodermal products. PGCs are not observed in Prdm1 mutants but were not tested due to their overall poor sampling. Yap1 mutants show weaker overall depletion of specific targets, with significant depletion of Epicardium, Neural Epithelium, and Erythroid Progenitors (EryP), consistent with prior reports of vascular defects^81^. **d.** (Top) Co-embedded reference UMAP of WT mesodermal cell types for Control and E8.5 Brachyury T mutant data. Leftmost panel shows the cell state composition with NMP and Tailbud-PSM highlighted, middle panel shows the expression of *T* within this subset, and conditions highlights the relative position of mutant cells, which are depleted within the tailbud mesoderm as previously reported for T-/- mESCs^82^. (Middle) Co-embedded reference UMAP of WT endodermal cell types for Control and E8.5 Prdm1 mutant data. Leftmost panel shows the cell state composition with Pharyngeal endoderm and Midgut highlighted, middle panel shows the expression of *Prdm1* within this subset, and conditions highlights the relative position of mutant cells, which are depleted within the Pharyngeal endoderm as previously reported^80^. (Bottom) Co-embedded reference UMAP of all WT cell types for E8.5 Control and Yap1 mutant data, including state annotations (leftmost panel) and expression (middle panel). Unlike *T* or *Prdm1*, *Yap1* is broadly expressed across these cell states, and joint projection shows that Yap1 mutant cells tend to occupy cell states from all three germ layers. **e.** Per replicate-state level normalized expression for *T* for relevant cell types. **f.** Per replicate-state level normalized expression for *Prdm1* for positive cell types. **g.** Fluorescence-based validation of the Yap1 mutant morphology as previously reported in Ref^81^ using phalloidin as a stain for general morphology. In general, Yap1 mutants are smaller than controls and exhibit diminished axial elongation, but all major morphological structures otherwise appear to be present. Scale bars = 100 µm.

**Extended Data Figure 7.**
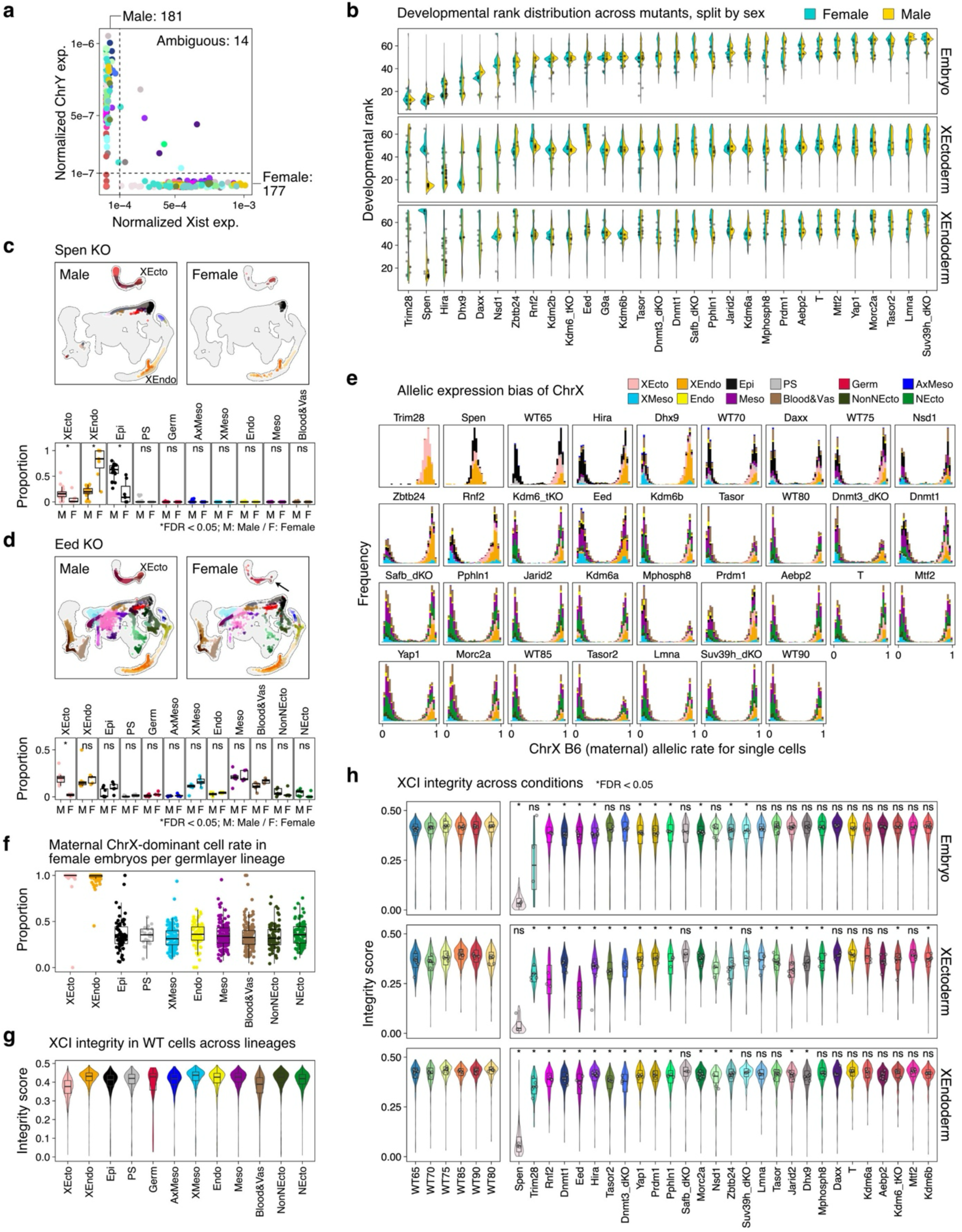
Distinct mechanisms for constitutive versus extraembryonic X inactivation. **a.** Scatterplot scoring of per replicate sex type using Xist versus ChrY expression for our full mutant cohort. Replicates are colored by regulator as in Fig. 2b. Note that Spen mutant females (light grey dots) show overall diminished Xist expression, consistent with a general failure to successfully consolidate ChrX inactivation^20,165–167^. Overall, we could successfully sex-type 339/352 (96%) of our mutant cohort in this manner. **b.** Transcriptional rank scores for our full mutant cohort, stratified by sex and embryonic versus extraembryonic status. Disrupted X regulation is observationally associated with accelerated differentiation within the extraembryonic tissues, particularly for Spen mutants, which primarily generate parietal endoderm (see below), but statistical significance was not observed at multiple-hypothesis-corrected *P*-value level (two-sided Wilcoxon’s rank-sum test). **c. Top:** UMAPs of sex-stratified Spen KO embryos show minimal differentiation of the embryonic (black) lineage, with substantial forward differentiation towards mature trophoblast cell types (dark red) within the placental lineage and of parietal endoderm (dark orange) in the yolk sac lineage. **Bottom:** Replicate level cell state proportions highlight the overabundance of XEndo cell states and depletion of embryonic or placental (XEcto cell states). Two-sided Wilcoxon’s rank-sum test followed by FDR adjustment: ns, not significant; \**P* <= 0.05. **d. Top:** UMAPs of male and female Eed KO embryos show that both sexes initiate gastrulation, but that female embryos are depleted of extraembryonic cell types beyond differentiated trophoblast cells. **Bottom:** Replicate level cell state proportions highlight the general underproduction of placental tissues within female embryos. Two-sided Wilcoxon’s rank-sum test followed by FDR adjustment: ns, not significant; \**P* <= 0.05. **e.** Per cell histograms of ChrX expression ratio between CAST/EiJ (Paternal) and B6 (Maternal), with cells separated and colored according to their broad developmental lineage. Spen mutant females show equal allelic ratios across embryonic and extraembryonic lineages, while the rest of our regulator cohort generally maintains stably bimodal transcriptional signatures consistent with ChrX inactivation. Extraembryonic ectoderm and endoderm exhibit imprinted X inactivation, such that ChrX is exclusively maternal (highlighted by the heavily right shifted, unimodal distribution of orange and pink cell states). **f.** Per replicate assignment of ChrX allelic ratios, stratified by major lineage. Each dot represents female replicates in the entire mutant and reference cohort (except Spen mutants), with the ratio estimated as the fraction of cells expressing the maternal allele divided by the total. As expected, extraembryonic ectoderm and extraembryonic endoderm states are imprinted, such that only the maternal chromosome is expressed^89^. ChrX selection is bimodal in embryonic states and favors expression of the CAST/EiJ (paternal) chromosome, hence the mean X maternal vs X paternal ratio of ∼30%^98^. **g.** Per cell XCI integrity scores for the control reference, calculated as the normalized fraction of transcripts recovered from the inactivated ChrX within single cells (see **Methods**). Cells are grouped by broad lineage. Within embryonic cells and the yolk sac, very few transcripts are recovered from the inactivated ChrX, and XCI integrity is generally high. However, in the placenta (XEcto), XCI integrity appears to erode, with a detectable proportion of inactivated ChrX transcripts found within the majority of cells. **h.** XCI integrity scores for our full mutant cohort, stratified by embryonic, placental (XEcto) and yolk sac (XEndo). Spen mutants show equivalent transcription from both chromosomes, while the other regulators show weak or no transcription from the inactivated X within embryonic or yolk sac lineages. In contrast, the placental lineage exhibits leaky XCI, with higher levels of inactive Chr X transcription observed for Eed, Rnf2, Trim28, Hira, Dnmt1, Dnmt3a/b, and other mutants. As such, the placenta appears to exhibit a greater amount of permissive transcription, even within heavily heterochromatic regimes. Each dot represents an embryo replicate. Wilcoxon’s two-sided rank-sum test with FDR adjustment: ns, not significant; \**P* <= 0.05.

**Extended Data Figure 8.**
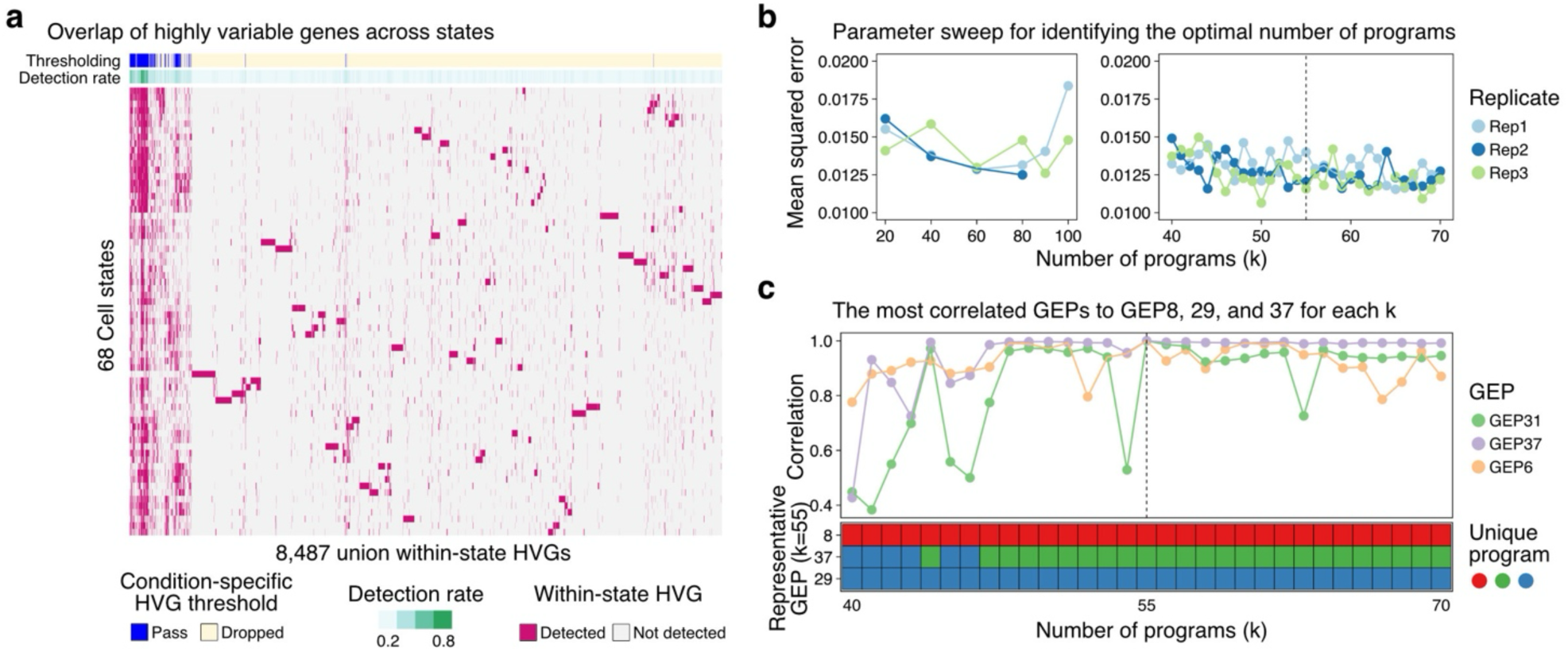
Identifying major axes of gene expression control by NMF. **a.** Selection criteria for including genes into our NMF-based description of Gene Expression Programs (GEPs). Briefly, the heatmap shows the collective union of 68 transcriptional state level comparisons between the control reference and our mutant cohorts, from which the top 500 most highly variable genes (HVGs) from each comparison were selected. Of this initial set of 8,487 regulator-specific HVGs, 308 were selected based upon a detection rate >= 0.2 (meaning, called as highly variable in at least 20% of state-level comparisons). To balance the selection of regulator mutant-associated expression patterns with those associated with normal embryonic development, we combined this gene set with the 2,000 most HVGs regardless of the sample conditions. In total, this led to a set of 2,261 genes for NMF-based analysis. **b.** To determine the optimal number of GEPs, we performed a parameter sweep for set factors (k) from 20-100 in intervals of 20 and from 40 to 70 in intervals of 1. “Mean squared error” on three independent parameter sweeps of the data suggested that the error was stable over this interval, suggesting good decomposition of the data in program usage and structure matrices (see Fig. 2). **c.** Our decision to decompose our data into 55 factors was determined empirically by investigating the relative stability of PRC (GEP8), HUSH (GEP29), and DNA/Chromatin (GEP37) programs as a function of k. Specifically, we plotted the maximal correlation of each GEP selected at k=55 to any program detected for each interval of our 40-70 parameter sweep. GEP8 was stable across this k range, with a resulting factor (program) showing high correlation to GEP8 at any seed number. In contrast, GEP29 and GEP37 tended to merge at lower k in a manner consistent with a higher degree of transcriptional interaction between the DNA/Chromatin regulators and the HUSH subunits. k=55 represented an early window where these two programs were stably separated and maintained their independent enrichment for DNA/Chromatin methyltransferases and the HUSH complex members, respectively.

**Extended Data Figure 9.**
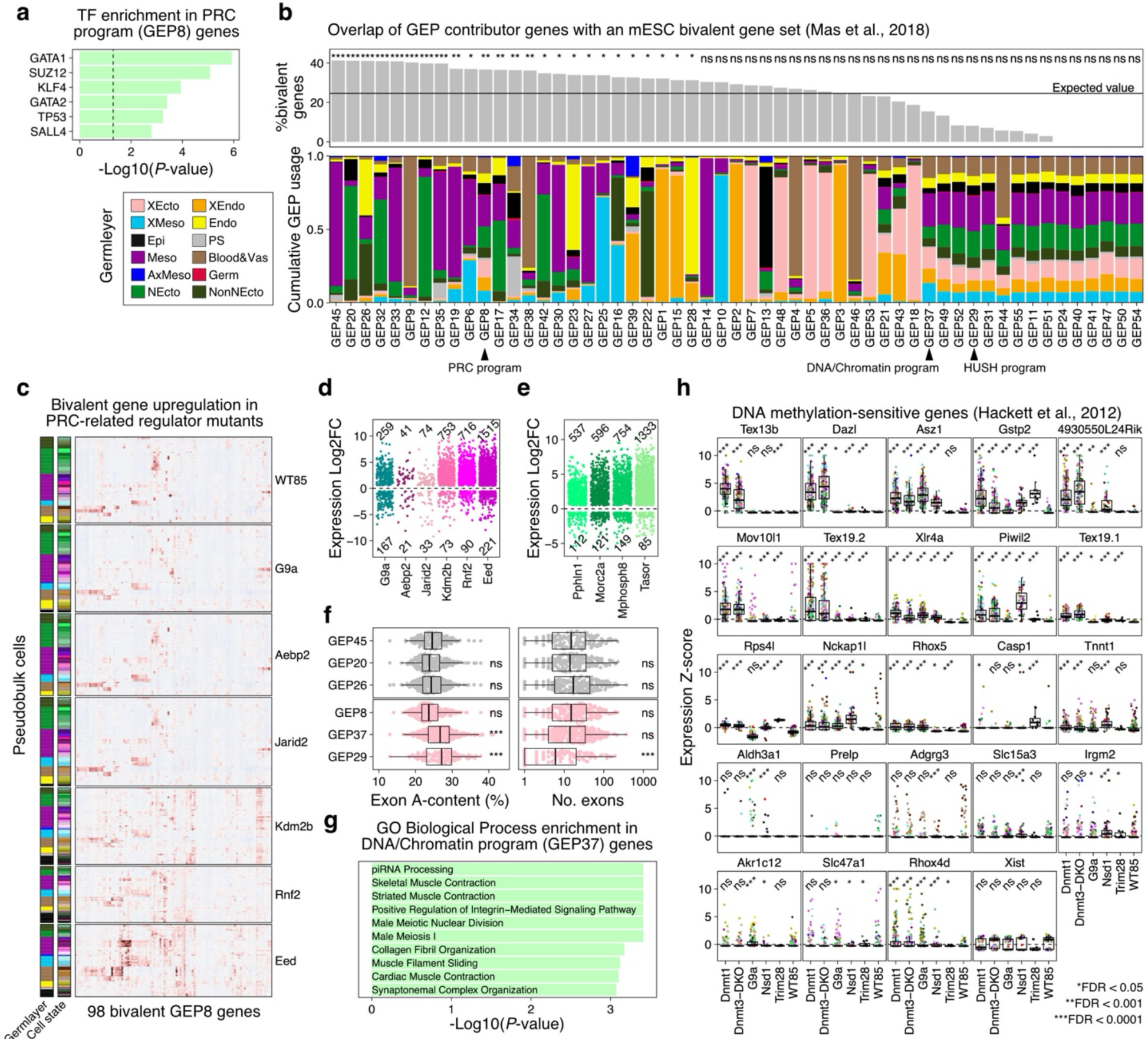
Genomic features of key gene expression programs. **a.** TF enrichment for PRC program (GEP8) genes, tested on ENCODE and ChEA consensus TF set using Enrichr^168^. In addition to enrichment for GATA and KLF factor motifs, GEP8 shows a strong enrichment for the PRC2 component SUZ12. **b. Top:** Percent of positively contributing genes for each GEP that are characterized as bivalent genes in mESCs (Data taken from Ref^169^), with PRC program (GEP8), DNA/Chromatin program (GEP37), and HUSH program (GEP29) highlighted. Significance is tested using the two-sided Fisher’s exact test followed by FDR adjustment. ns, not significant; \**P* <= 0.05; \*\**P* <= 0.001, \*\*\**P* <= 0.0001. **Bottom:** Cumulate GEP usage for each individual GEP, colored by germ layer. The PRC program is distinguished from the DNA/Chromatin and HUSH programs by a significant enrichment for bivalent genes. **c.** Normalized expression heatmap of 98 bivalent genes that intersect with GEP8. Rows correspond to individual replicate-states and are grouped by mutant cohort. In WT, these genes show more lineage specific expression patterns, characterized by stripes of positive expression within select lineages. In core PRC regulator mutants (Eed and Rnf2), expression is expanded to more cell types, a trend that is more muted for auxiliary cofactors such as Kdm2b or Jarid2. **d.** Expression Log2-fold change for individual PRC program (GEP8), where each dot represents a given gene in a given replicate state. Only genes that are called as significant by DESeq2-analysis are shown (**Methods**). Overall, Rnf2, Eed, and Kdm2b show the highest fraction of upregulated GEP8 genes across different cell states, followed by G9a. Transcriptional effects in Jarid2 and Aebp2 are more muted. **e.** Expression Log2-fold change as in **d** for individual HUSH program (GEP29) genes, where each dot represents a given gene in a given replicate-state. Overall, Taser shows the largest fraction of upregulated genes across the broadest number of states, followed by Mphosph8. **f.** Exon-A content (left) and number of exons (right) for three selected lineage specific GEPs (GEPs 45, 20 and 26) as they compare to the PRC program (GEP8), DNA/Chromatin program (GEP37) and HUSH program (GEP29). Both the DNA/Chromatin and HUSH programs are significantly A-rich, while only the HUSH program shows the biochemically expected association with smaller or single exon genes. Wilcoxon’s two-sided rank-sum test with FDR adjustment: ns, not significant; \*\*\*\**P* <= 0.0001. **g.** GO Biological Process enrichment for DNA/Chromatin program (GEP37) genes show significant enrichment for functions related to meiosis and genome defense, including piRNA processing and Male Meiosis. **h.** Normalized replicate state level expression for 24 genes previously defined as DNA methylation sensitive and associated with germline genome defense^94^. All but Xist show significant expression in at least one GEP37-positive mutant cohort, with expression dynamics reflecting differential repressive roles for select cofactors. Wilcoxon’s two-sided rank-sum test with FDR adjustment: ns, not significant; \**P* <= 0.05; \*\**P* <= 0.001; \*\*\**P* <= 0.0001.

**Extended Data Figure 10.**
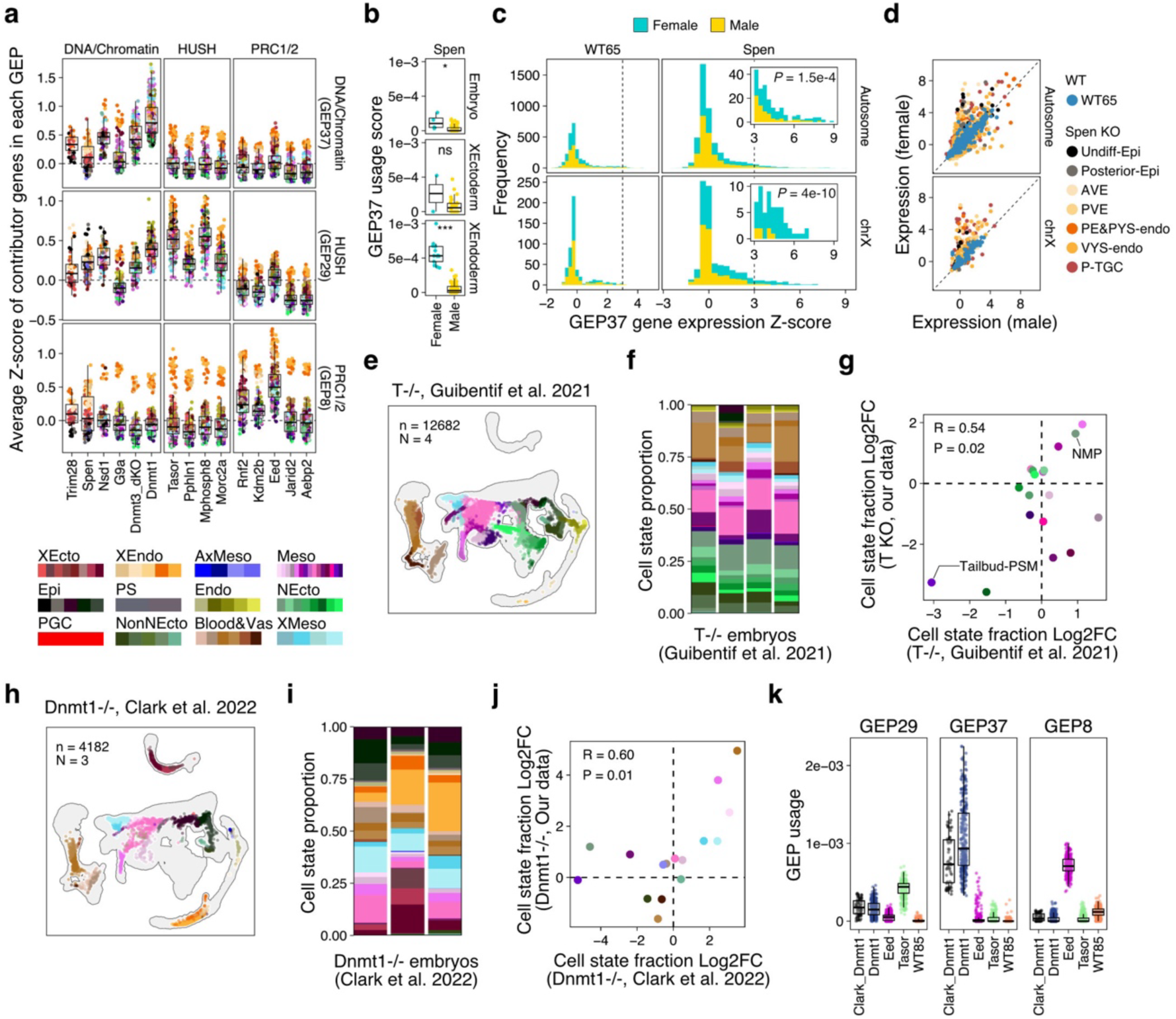
Stratification of GEP activity across replicates and cell states. **a.** Average Z-score expression for GEP-contributing genes across replicates and knockouts, shown for GEP-positive conditions across the DNA/Chromatin (GEP37), HUSH (GEP29), and PRC (GEP8) programs. Each dot corresponds to an individual replicate state. Overall, we observe consistent expression differences between regulators of the same class, with DNMT1 showing dominant upregulation within the DNA/Chromatin cohort and many of these exhibiting interactions with the HUSH cohort. Within the PRC cohort, Eed mutants show the highest level of de-repression, followed by Rnf2 and Kdm2b. These expression differences are consistent for the majority of states. Note also that GEP8 appears to show heightened average expression within extraembryonic endodermal cell states (oranges), which could indicate additional interactions with genes related to this specific lineage. **b.** DNA/Chromatin program (GEP37) usage score for our Spen mutant cohort, stratified by sex. Spen mutants exhibit female-specific program usage, which is significant in both embryonic and extraembryonic endodermal cell states. ns, not significant; \**P* <= 0.05; \*\*\**P* <= 0.0001. **c.** Expression Z-score histogram for DNA/Chromatin (GEP37) genes in E6.5 WT and Spen mutant embryos, separated by sex (color) and by autosomal (top) and ChrX (bottom). Spen mutant females show specific enrichment for upregulated GEP37 genes, including a larger fraction on the X chromosome (inset highlights genes with an expression Z-score >3 in Spen mutants. Plot represents the computed change in regulation at the level of individual cell states. Statistical tests were performed for the sex ratio in the Z-score > 3 fraction using Fisher’s exact test. **d.** Scatterplot of DNA/Chromatin (GEP37) program gene expression in male versus female embryos, colored by state for Spen mutants and overlaid by WT E6.5 cells. Overall, many X-linked GEP37 genes show female specific deregulation in Spen mutants, indicating an interaction between the Dnmts and Spen in ChrX silencing. **e.** Projection of T mutant cells generated by embryo injection and collected at E8.5 by Guibentif et al., projected onto our WT reference for comparative analysis. As these are mESC-based, there is no contribution to extraembryonic cell types (extraembryonic ectoderm or endoderm, respectively). **f.** Relative cell state composition of individual replicate embryos generated by Guibentif et al. according to our reference labels. **g.** Scatterplot comparing the relative cell state abundance (Log2-FC) between Guibentif et al. experimental and control samples compared to our WT reference and T mutant cohort. Overall, both datasets confirm a general depletion of Tailbud PSM and enrichment for NMPs, with off-diagonal measurements likely reflecting subtle deviation in the relative timing of embryo isolation. Overall correlation (R) and *P*-value are also highlighted. **h.** Projection of Dnmt1 mutant embryos generated by standard breeding of transgenes by Clark et al.^32^, projected onto our WT reference for comparative analysis. **i.** Relative cell state composition of individual replicate embryos generated by Clark et al. according to our reference labels. Overall, Dnmt1 embryos from Clark et al. appear to be developmentally delayed, as apparent by an enrichment for epiblast cells at E8.5, though it is not clear if this reflects difficulties in selecting embryos via breeding or differences in embryonic sampling. **j.** Scatterplot comparing the relative cell state abundance (Log2-FC) between Clark et al. experimental and control samples compared to our WT reference and Dnmt1 mutant cohort. Despite apparent differences in sampling, overall enrichment and depletion of specific lineages are generally correlated. Overall correlation (R) and P-value are also highlighted. **k.** GEP usage scores for individual replicate-states for our three major epigenetic regulator programs, including HUSH (GEP29), DNA/Chromatin (GEP37), and Polycomb (GEP8), shown for three select mutants from our cohort as they compare to the Dnmt1 mutant embryos sampled by Clark et al. Overall, GEP37 shows very high specificity, with shared misexpression for breeding-based mutant embryos compared to our Crispant approach.

**Extended Data Figure 11.**
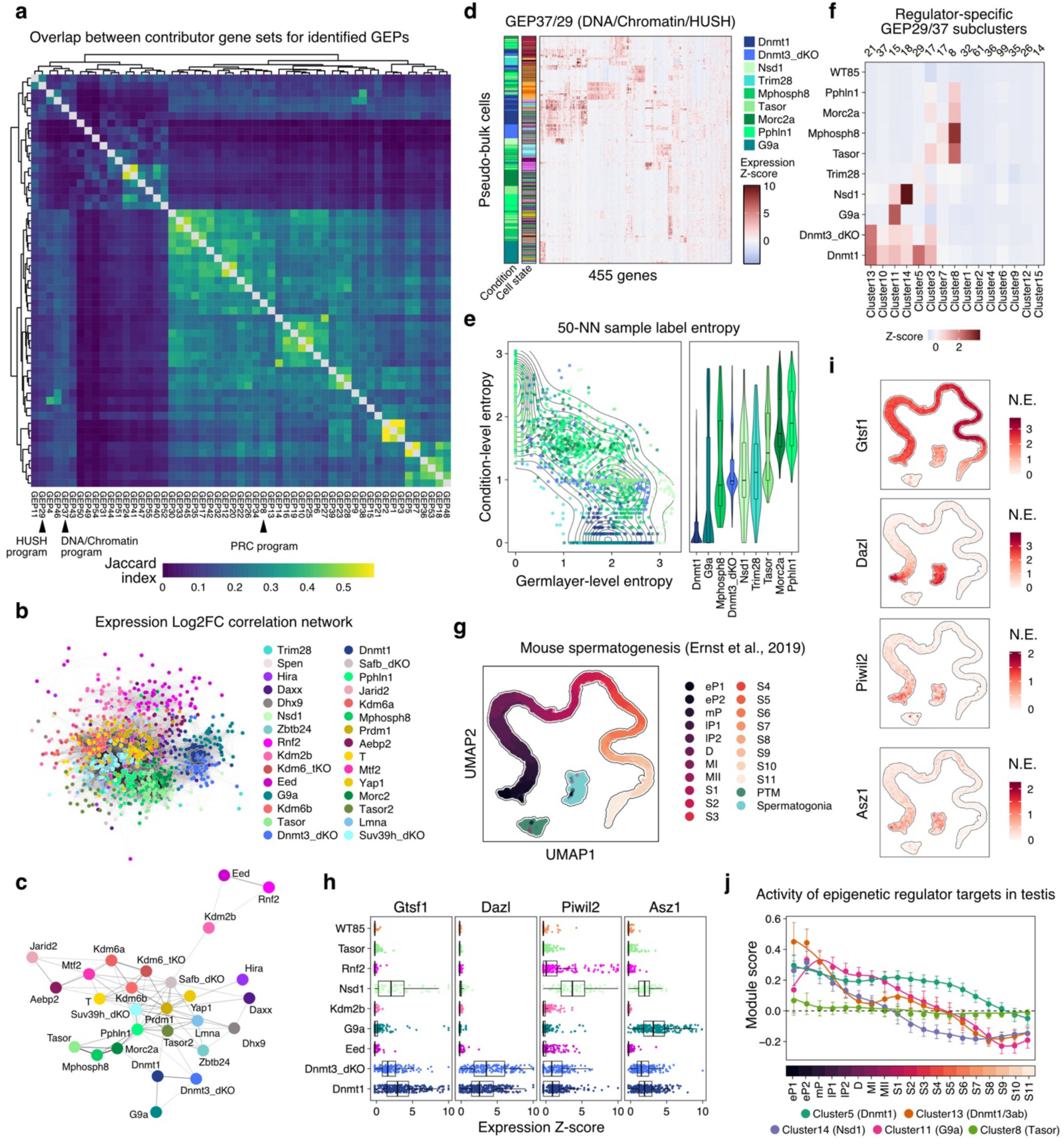
Epigenetic networks show a diversity of heterochromatin-based gene repression. **a.** Heatmap of the Jaccard index showing overlap between individual GEPs. Arrows point to the PRC program (GEP8), HUSH program (GEP29), and DNA/Chromatin program (GEP37), respectively. The PRC program is positioned within a large network of endogenous GEPs with developmental functions. In contrast, GEPs 37 and 29 are disconnected from this dense hub and share comparatively minimal overlap with each other. **b.** Reconstruction of network interactions based on differential gene expression, rather than GEP contribution scores (as shown in Figure 3a). Briefly, the differentially expressed genes and their Log2-fold changes were calculated for each cell state between a given mutant cohort and wild type reference by DESeq2, and overexpressed genes with *P*-value < 0.01 and Log2-fold change > 1 were used to build a correlation network between individual states. Graph edges with Pearson’s correlation < 0.2 were removed. As with GEP usage, states from the same mutant cohort tend to cluster together, and show general interactions with members of the same pathway (greens, HUSH mutant regulators; Blues, DNA/Chromatin; Purples/magentas, PRC). **c.** Same as in **b** but collapsed to the level of individual regulators, rather than regulator cell states. The network broadly maintains strong interactions between core regulators from either the PRC, DNA/Chromatin, and HUSH programs. However, more subtle interactions, such as those between PRC auxiliary subunits such as Jarid2 and Aebp2, are lost at this level of resolution. **d.** Hierarchical clustering of cell state level expression values for HUSH (GEP29) and DNA Chromatin (GEP37) contributing genes. Cell state and sample identity are indicated on the left. GEP29 and 37 cluster more frequently by the individual regulator mutant than by state, suggesting that each regulator oversees a distinct subset of target genes. **e.** Summary analysis of the heatmap **d**, focusing on the condition versus germ layer transcriptional response for GEPs 29 and 37 positive regulators. Specifically, for each regulator-state pseudobulk cell, the 50 nearest neighbors were selected, and their label profile (either condition or germlayer state) was converted into an entropy score. Most regulator mutants show substantially lower condition-level entropy, suggesting that these regulators have discrete target preferences within the larger umbrella of regulation by DNA methylation. **f.** Summary expression levels for GEP29/37 subclusters highlight the degree to which these programs represent an interconnected hub. Many clusters reflect co-regulation between one or several specific co-factors (such as the Dnmt3s, Nsd1, G9a, Trim28, or Tasor/Mphosph8) and Dnmt1. HUSH mutants also have a Dnmt1-independent subcluster (Cluster 8). The number of genes in each cluster is annotated above the heatmap. **g.** UMAP representation of mouse male germ cell differentiation taken from Ernst et al.^163^ that samples from pre-meiotic spermatogonia through meiosis and sperm development. **h.** Expression dynamics for representative genes with known functions in germline gene development, including those that show more specific regulation by Dnmt3 (Dazl), Nsd1 (Gtsf1, Piwil2) or G9a mutation (Asz1). **i.** Corresponding expression dynamics for Gtsf1, Ddx4, Piwil2 and Asz1 over male germ cell differentiation highlight the relative enrichment for many of these genes towards the early stages of meiosis and prior to the period of spermatid differentiation (s1-s11). **j.** Seurat-calculated single-cell module score of selected GEP29/37 subclusters over spermatogenesis. In general, Dnmt1-exclusive targets (Cluster 5) are distributed more broadly, while Nsd1-, G9a-, or Dnmt3-shared genes (Clusters 13, 14 and 11, respectively) are more enriched for early meiosis stages (pachytene and diplotene). Number of cells for each cell state-by-gene subcluster combination = 375-1513 (mean = 821.9; s.d. = 328.0). Error bar indicates standard deviation.

**Extended Data Figure 12.**
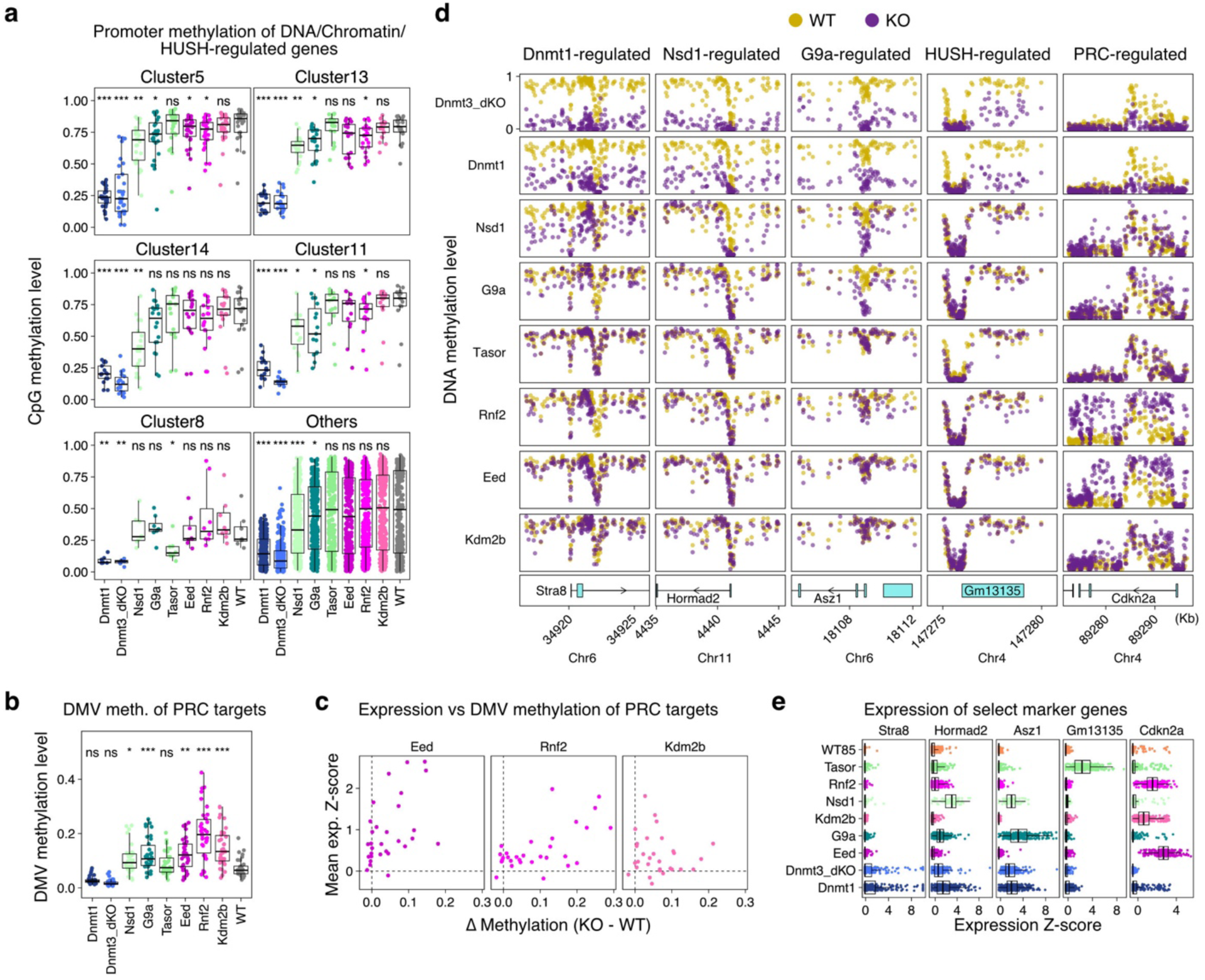
Epigenetic changes precede gastrulation phenotypes. **a.** Promoter methylation levels corresponding to the difference in methylation plots shown in Fig. 3e, including statistical testing in comparison to WT. Note that, of these clusters, the Tasor-specific subgroup (cluster 6) generally reaches lower promoter methylation values in WT Epilast, reflecting the distinct nature of these targets. Wilcoxon’s two-sided rank-sum test with FDR adjustment: ns, not significant; \**P* <= 0.05; \*\**P* <= 0.001; \*\*\**P* <= 0.0001. **b.** Promoter methylation levels of bivalent DMVs corresponding to the difference in methylation plots shown in Fig. 3f. Wilcoxon’s two-sided rank-sum test with FDR adjustment: ns, not significant; \**P* <= 0.05; \*\**P* <= 0.001; \*\*\**P* <= 0.0001. **c.** Scatterplot comparing the average gene expression Z-score for individual PRC program genes (GEP8) in PRC mutants as they correspond to changes in DMV methylation. Eed and Rnf2 mutants show the strongest relationship, including substantial deregulation of many of GEP8 genes, while the effects of Kdm2b mutation are more muted and variable. **d.** Genome browser tracks of select genes that show distinct expression and DNA methylation patterns, including those associated with individual factors of the DNA/Chromatin/HUSH programs (GEPs 29 and 37). The PRC-program associated gene Cdkn2a is shown for comparison, which shows broad increase in methylation in surrounding DMVs. **e.** Expression of example genes shown in **d**, highlighting the relative dominance of individual DNA/Chromatin program-associated factors over distinct subprograms.

**Extended Data Figure 13.**
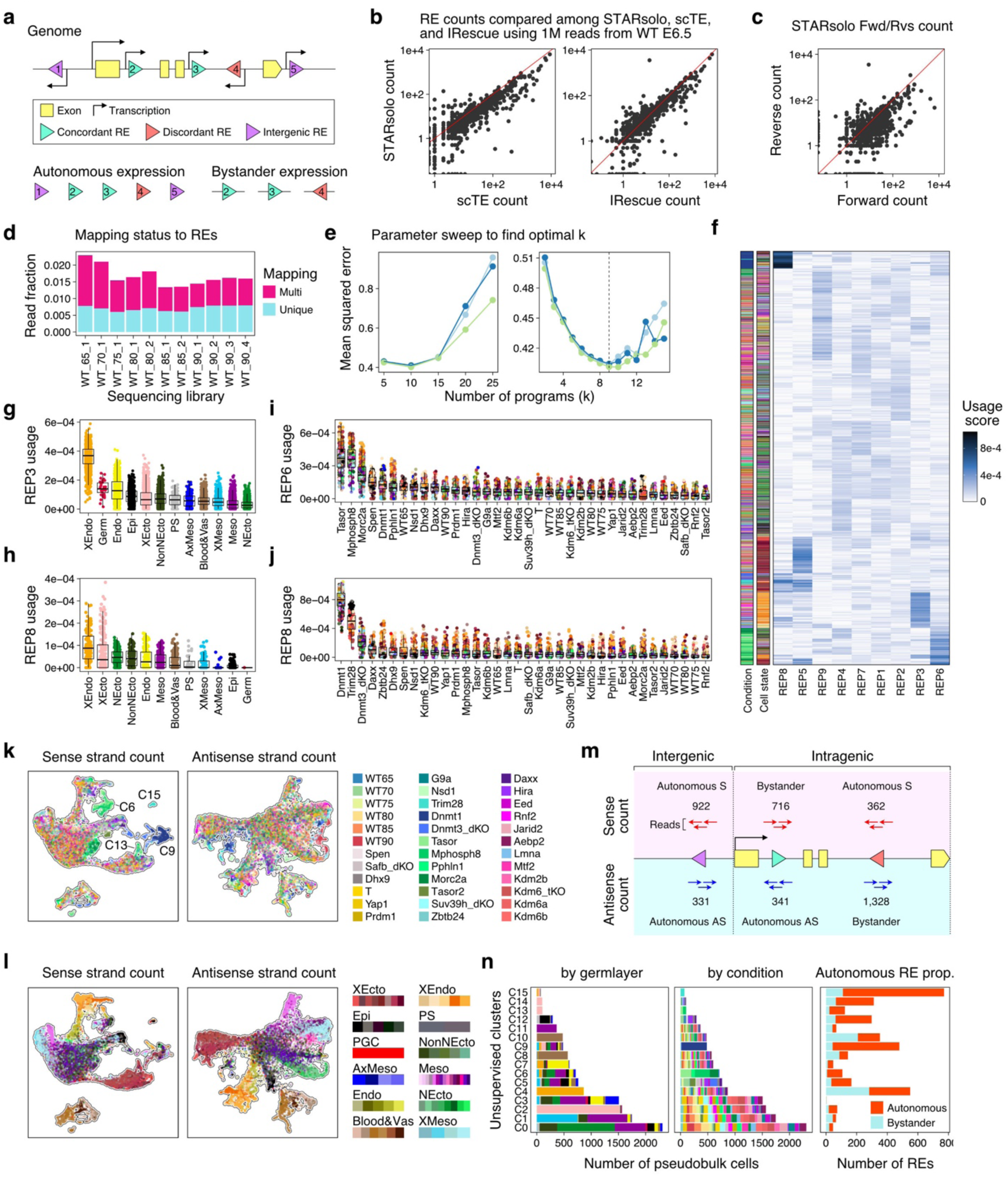
Discriminating autonomous versus bystander repeat expression. **a.** Schematic depicting various repetitive element (RE) contexts that could impact the interpretation of single-cell transcriptional profiles. Briefly, autonomous expression refers to transcripts that are driven by promoter elements within the RE element itself. In the majority of contexts, this will be for reads that align to the sense strand. Reads that fall within gene bodies (intragenic, intronic) could also reflect “bystander” transcription, regardless of the direction: “concordant” RE insertions are on the same strand for both the host gene and the embedded RE, while “discordant” RE insertions are opposite to the host gene direction. To interrogate the degree of autonomous RE expression in Fig. 4 and 5, we primarily focused on sense-strand RE alignments that are either intergenic or intragenic and discordant. Leveraging the modularity of the STARsolo pipeline, we examined the ratio of sense to antisense, intergenic versus intragenic, and concordant versus discordant insertions in order to examine how autonomous versus bystander RE transcripts determine the identity of cell clusters (see below, **Extended Data Fig. 13k-n**). **b.** Systematic comparison of RE family counts by our STARsolo pipeline against previously published single-cell repetitive element aligners, including scTE^156^ and IRescue^157^, using 1M reads from our reference E6.5 library. Overall, STARsolo returned RE expression counts consistent with the previously built pipelines. **c.** Comparison of RE family counts for STARsolo after splitting reads by their sense (forward) and antisense (reverse) alignments. Overall, more counts are recovered from the forward count, in keeping with the recovery of at least some autonomous RE transcription. **d.** Mapping status of REs after alignment to individual elements using STARsolo for the control reference data set. In general, approximately one-third of repetitive element alignments can be uniquely assigned within the control reference. **e.** Parameter sweep for the optimal k for NMF-based detection of Repetitive Element Expression Programs (REPs) across three independent trials (colored by blue, light blue and light green), including coarse (5,10,20, 30) and fine (1-15, interval 1) scales. Ultimately, mean squared error reaches a minimum at k=9 (our selected parameter) and rises shortly after k=10. **f.** Hierarchical clustering of program usage scores (loadings) for each of 9 Repetitive Expression Programs (REPs) identified using our regulator-state by RE data frame, with regulator and transcriptional state indicated on the left. In general, REP3 and REP5 cluster by cell type, with strong usage scores for the XEndo and XEcto, respectively. REP6 and REP8 cluster by sample, namely HUSH complex mutants and Dnmt1/Trim28 mutants. Additional REPs tend to show weaker usage scores that are distributed across a broader number of element families, and to generally separate different embryonic germ layers. Further investigation below suggests that these tend to reflect passive capture of older elements within the introns of lineage-specific genes. **g.** REP3 (XEndo) usage scores of individual replicate states across our control reference, stratified by major lineage. REP3 is highest for the XEndo lineage, but also shows some residual activity in the germ cell, endoderm, epiblast, and XEcto. **h.** Per replicate state level usage scores for REP8 (Dnmt1/Trim28) across our control reference, which shows higher endogenous activity in XEndo (Yolk Sac) and XEcto (Placental) states. **i.** Per-replicate cell state usage scores for REP6 (HUSH) across our full mutant cohort show the highest levels of activity for Tasor and Mphosph8, followed by Morc2a, Spen, and Pphln1. Notably, these signals are always strongest for extraembryonic states (orange and red hues for the XEndo/Yolk Sac and XEcto/Placenta, respectively) within the HUSH complex but show enriched activity within Dnmt1 mutant embryonic states (blue and green hues, respectively). See Fig. 4b for a summary description of these interactions. **j.** Per replicate cell state usage scores for REP8 (Dnmt1/Trim28) across our full mutant cohort show the highest levels of activity for Dnmt1 and Trim28, followed weakly by Dnmt3a/b, Daxx, and Zbtb24. In general, control states show higher basal activity within the extraembryonic lineages (orange and red hues, respectively). See Fig. 4b for a summary description of these interactions. **k.** Comparative UMAP embeddings of the top 2,000 highly variable, uniquely aligned RE copies (HVREs) that align to the sense (left) or antisense (right) strand of the annotated RE. Each dot represents an individual pseudobulk cell state colored by sample identity. Clear clusters can be observed for Dnmt1, Trim28, and HUSH. Note that, in the antisense alignment, Dnmt1/Trim28 clusters remain separated, but HUSH samples merge with the background. **l.** Same UMAPs as shown in **k** but colored by cell state, rather than sample. Dimensionality reduction shows more complexity when examining the sense strand data versus antisense, particularly for the embryo proper. **m.** Distribution of the 2000 most HVREs based on sense and antisense alignments. Alignments are distributed across three possible contexts: intergenic (scored as autonomous in either the sense or antisense direction), intragenic autonomous (if aligned in the opposite direction of the surrounding host gene), and intragenic bystander (if aligned in the same direction as the host gene). We found that, in the antisense count pipeline, a substantial amount of HVREs were derived from intragenic bystander fragments. Note that these scores represent an inference of the most conservative explanation for each transcript and are intended to evaluate the degree to which state-specific RE expression reflects autonomous RE activity. **n.** Clustering of pseudobulk cell states according to the sense alignment pipeline by Seurat’s unsupervised clustering, including the relative composition of cell states (by germ layer, left) and condition (right). Clusters 6, 9, 13 and 15 represent condition-specific programs related to Dnmt, Trim28 or HUSH-mediated repression and are heavily enriched for autonomous expression. Alternatively, Clusters 2, 4, 12 and 14 represent endogenous “autonomous RE enriched” programs and map to XEcto, early embryo (epiblast and early primitive streak), or XEndo cell types.

**Extended Data Figure 14.**
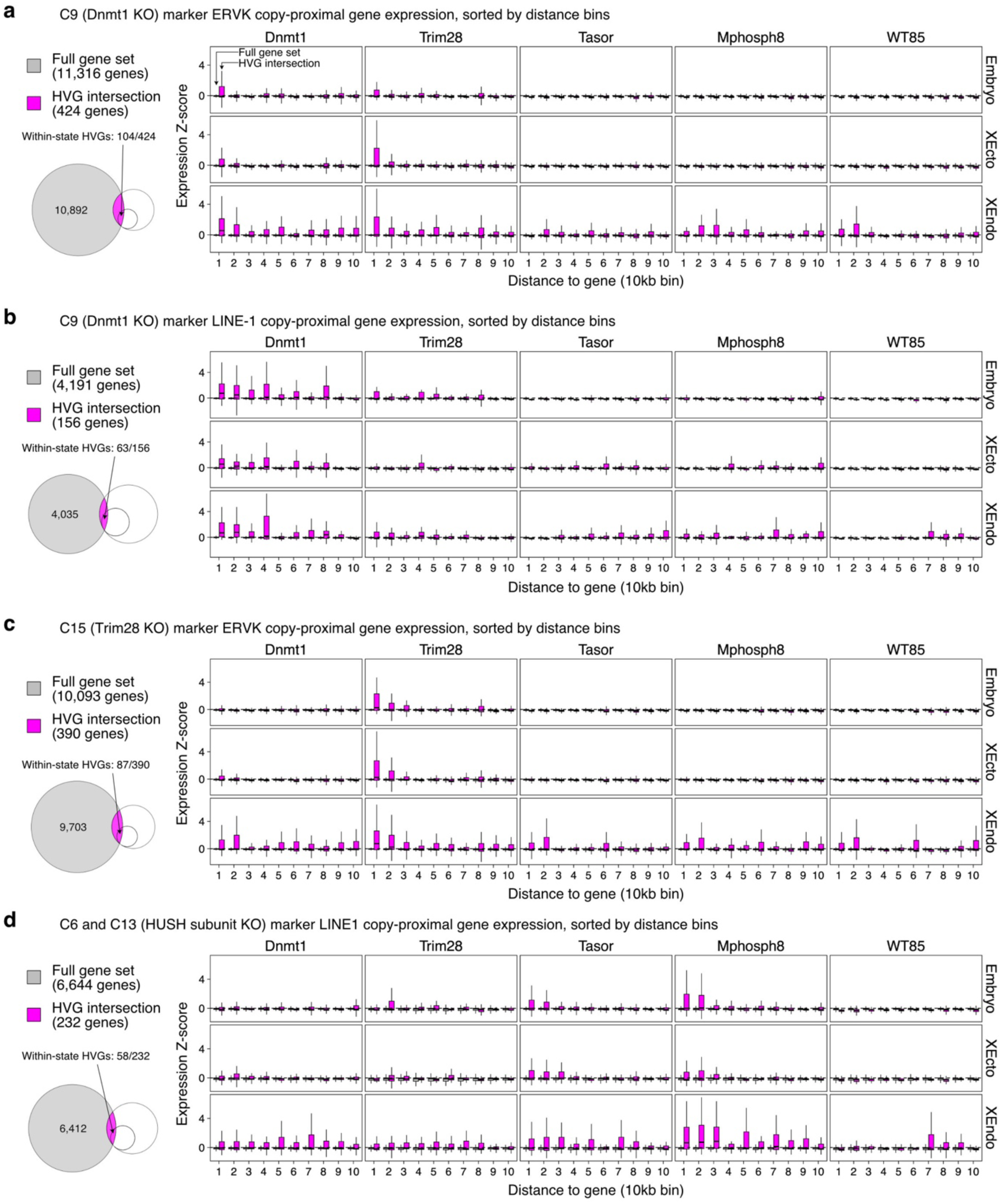
Examining cis-based effects of retrotransposon activation. **a.** Examination of proximity-based transcriptional associations between REs and genes for Dnmt1/Trim28-targeted ERVK elements (Cluster 9 in **Extended Data** Fig. 13n). Left: Euler plot highlights the number of genes whose TSS falls within 100 kb of a Cluster 10-specific ERVK, binned into 10 kb intervals (called from the sense-strand RE alignment in **Extended Data Fig. 13**). In general, we see no proximity based relationship for this larger gene set (grey), suggesting that the transcriptional status of most proximal genes are well insulated from the activity of nearby repetitive elements. However, when we examine the set of genes that overlap with highly variable genes (pink), we see some weak proximity based relationship within ∼10 kb of the gene TSS. This relationship is apparent for Dnmt1 and Trim28 mutants but not observed for other RE-associated regulator mutants. **b.** Same as in **a** but for Cluster 9 (Dnmt1/Trim28) associated LINE-1 elements. While the global stability of proximal gene expression holds, de-regulated LINE-1s show an extended proximity-based relationship to highly variable genes over a greater interval. **c.** Same as in **a** but for Cluster 15 (Trim28) associated ERVK elements, which show a heightened proximity based correlation between ERVKs and highly variable genes. This trend holds despite these responses being directed towards a different set of genes as observed for Dnmt1/Trim28 elements in **a**. RE-Gene transcriptional relationships appear to be distinct according to their specific regulatory targets. **d.** Same as in **a** but for Clusters 6 and 13 (HUSH) associated LINE-1 elements, which show the same general relationships, including a heightened proximity based correlation between LINE-1s and highly variable genes. This trend holds despite these responses being directed towards a different set of genes as observed for Dnmt1/Trim28 elements. RE-Gene transcriptional relationships appear to be distinct between HUSH and Dnmt1/Trim28 targets.

**Extended Data Figure 15.**
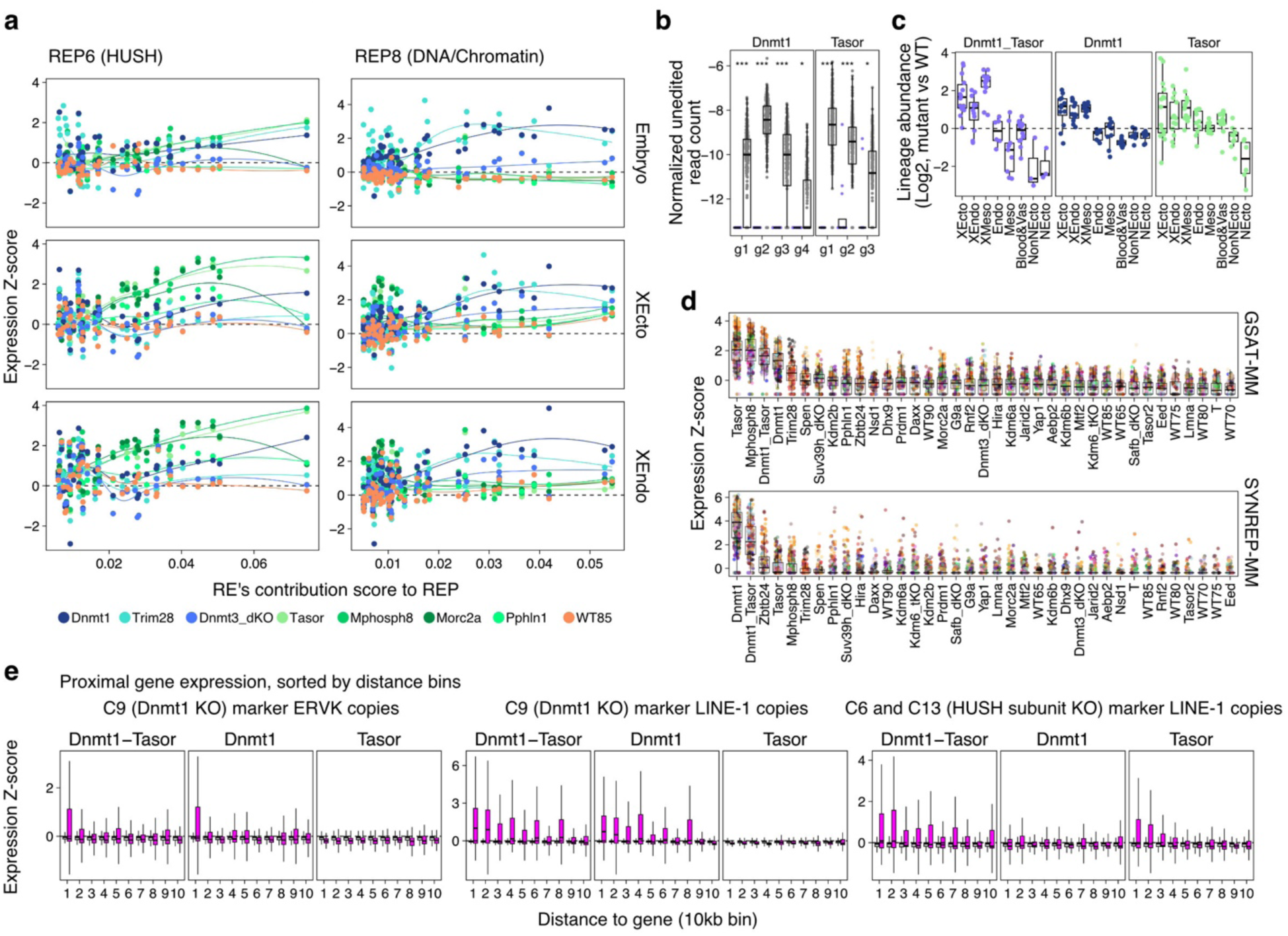
Dnmt1 and HUSH cooperate more extensively in embryonic tissues. **a.** Scatterplots showing the relative contribution scores of individual RE families to the HUSH and Dnmt1/Trim28 programs (REP6 and REP8) respectively, alongside their family-level expression. Mean expression Z-score is calculated for the full data set and shown individually for the Embryo, Placental (XEcto) and Yolk Sac (XEndo) lineages, respectively. Within the HUSH program, Dnmt1, Trim28, Tasor, and Mphosph8 mutants show generally consistent expression changes for the most highly contributing REs in embryonic cells, while Tasor and Mphosph8 show enhanced levels of derepression within the extraembryonic lineages. The Dnmt1/Trim28 program shows very little interactions with the HUSH subunits. However, these targets exhibit higher latent activity within extraembryonic tissues, which can be seen in both the HUSH subunit mutants and E8.5 WT embryos (orange). **b.** Validation of simultaneous Tasor/Dnmt1 gene disruption by measuring recovered transcriptions at sgRNA cut sites. Data is presented as in **Extended Data Fig. 2,** with a mean inferred edited efficiency of 100% for both Tasor and Dnmt1. The number of replicates = 12 for dKO and 72 for WT. Statistical tests were performed by one-sided Wilcoxon’s rank-sum test followed by FDR adjustment. \**P* <= 0.05; \*\**P* <= 0.001; \*\*\**P* <= 0.0001 **c.** Per lineage fold change of cell abundance between time matched mutant and WT embryos. Subtle distortions in embryo patterning within Tasor and Dnmt1 single mutants, including over-representation of extraembryonic states and under-representation of Neural Ectoderm (NEcto) states, are compounded in double mutant embryos, including severe over-representation of early primitive streak products and minimal germ layer induction. Single mutant data are also shown in **Extended Data Fig. 5**. **d.** Gene expression Z-scores for major/pericentromeric (GSAT-MM) and minor/centromeric (SYNREP-MM) expression across single regulator mutants, ranked from highest median value to lowest. GSAT derepression is highest for Tasor and Mphosph8, followed by Dnmt1, Trim28, Spen, and Suv39h1/2. In contrast, SYNREP derepression is greatest for Dnmt1 and Zbtb24, followed by Tasor and Mphosph8. **e.** Gene by element proximity analysis as in **Extended Data** Fig. 14, including for Dnmt1-specific ERVKs and LINE-1s as well as HUSH-specific LINE-1s. This comparison highlights the specificity of these interactions between individual regulators, as the double mutant embryo recapitulates both transcriptional responses simultaneously.

**Extended Data Figure 16.**
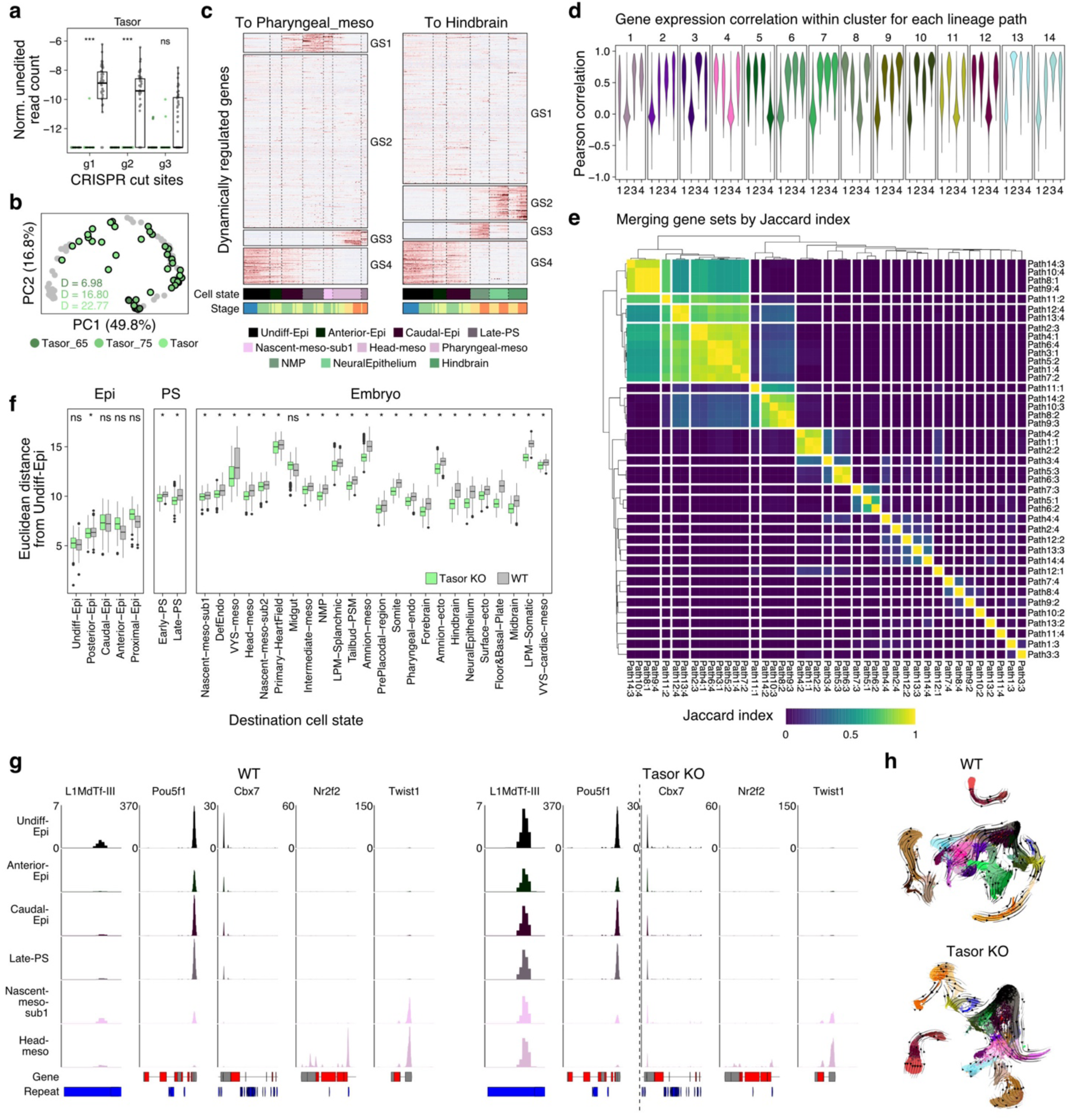
Tasor mutant embryos display compromised transcriptional robustness. **a.** Validation of Tasor disruption in the time course data by measuring recovered unedited transcripts at the editing sites of injected sgRNAs. Data is presented as in **Extended Data Fig. 2**, with inferred editing efficiencies of 100% for both E6.5 and E7.5, respectively. Statistical tests were performed by one-sided Wilcoxon’s rank-sum test followed by FDR adjustment. \*\**P* <= 0.001; \*\*\**P* <= 0.0001. **b.** Projection of individual Tasor mutants onto the cell state composition PCA as shown in Fig. 1f, with distance to centroid (D) highlighted. In general, arrest points in gastrulation tend to emerge over the gap between E7.5 and E8.0 during a major jump in embryonic complexity. Furthermore, embryonic variation increases as a function of developmental time, suggesting that it reflects a bona-fide feature of the developmental phenotype rather than a reflection of our Crispant strategy. **c.** Gene expression K-means clustering (k=4) for two representative trajectory paths used to infer differential expression dynamics between control and mutant cells: from undifferentiated epiblast to pharyngeal mesoderm and from undifferentiated epiblast to hindbrain (Paths1 and 6). GS: Gene set. **d.** Pearson’s correlation coefficient of gene expression Z-scores within the K-means clusters. Values were plotted for each cluster across the 14 paths. **e.** Pairwise Jaccard index matrix of the K-means cluster gene sets with a minimum Pearson’s correlation coefficient of 0.3. The heatmap is split by hierarchical clustering. Genes within the same Jaccard index clusters were merged to summarize the transcriptional state of a single shared progenitor. **f.** Summary of inherited cell state similarities in Tasor KO embryos from the undifferentiated epiblast (see Fig. 6g for a single trajectory). Euclidean distances between the undifferentiated epiblast and each cell state were collected for the WT and Tasor KO data. In general, even late stage gastrulation products (such as the Neural Epithelium) maintain a relatively closer transcriptional similarity to undifferentiated epiblast cells in Tasor mutants. Significance of the similarity to the undifferentiated epiblast was assessed by the one-sided Wilcoxon rank sum test, followed by FDR adjustment. ns, not significant; \**P* ≤ 0.01. **g.** Representative genome-browser tracks for loci that are emblematic of the breakdown in cell state transitions in Tasor embryos, including failure to deactivate aspects of the pluripotency program and reactivate differentiated lineage-specific genes. Tracks are colored by cell state and highlight the imputed progression from Undifferentiated Epiblast (black) to Head Mesoderm (pink). Loci include: an L1MdT element that is associated with preimplantation development and remains constitutively deactivated in Tasor mutants; Cbx7, which is associated with the pluripotent state and remains active in later states within Tasor mutants; Nr2f2, which is induced in WT Head Mesoderm but not in Tasor mutants; and Twist1, which is induced in both WT and mutant cells with similar dynamics. **h.** Independently generated WT and Tasor KO UMAP embeddings, overlaid with RNA velocity flows. In general, the UMAP embedding of Tasor KO differentiation shows lower transcriptional diversity, with a higher density of cells within early gastrulation states and heavy arrest upon entry into the Neural Ectoderm (green states). Although qualitative, these observations are broadly consistent with diminished transcriptional boundaries and a failure to robustly induce cell states associated with the phylotypic period.

**Extended Data Figure 17.**
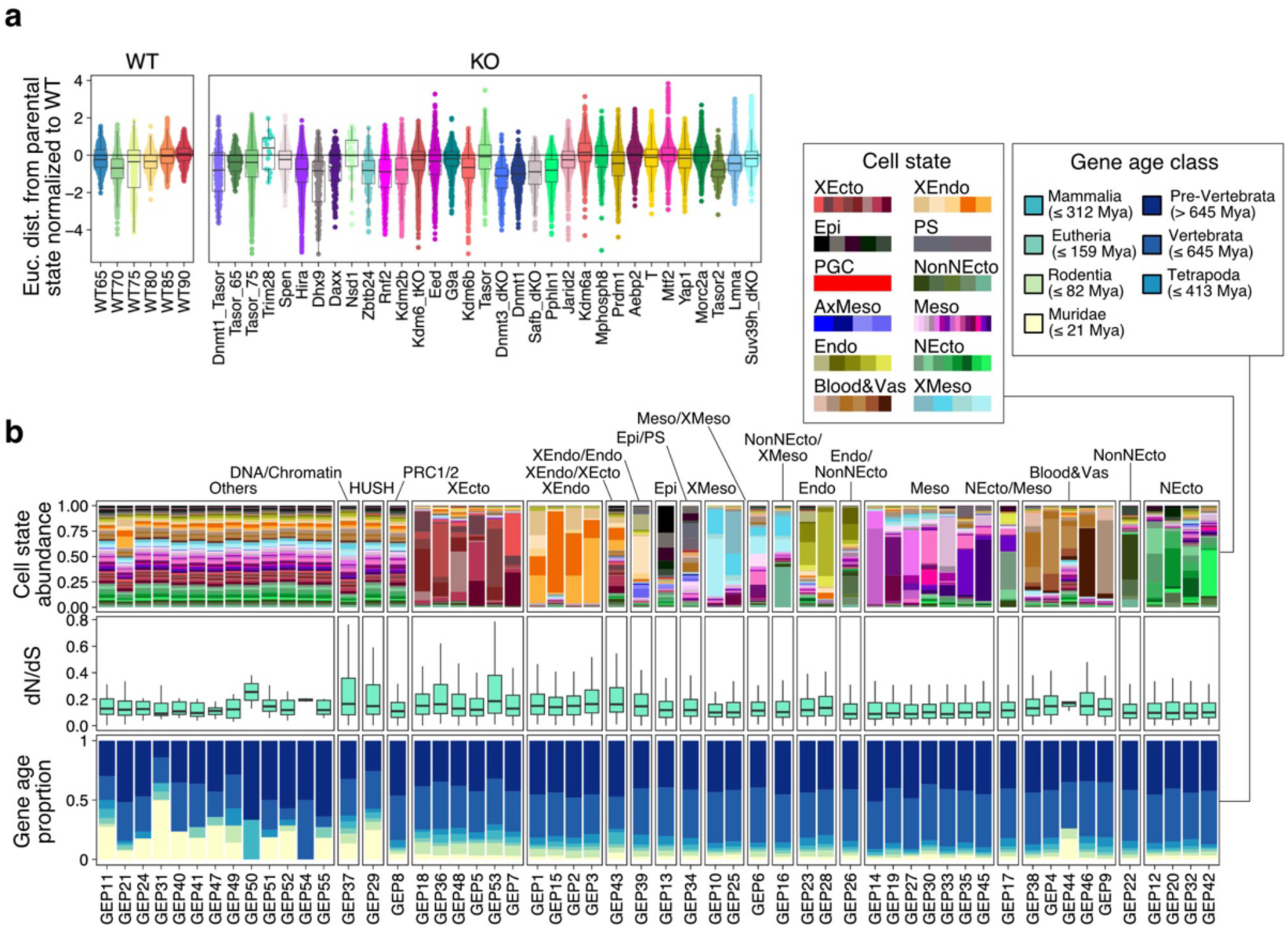
Epigenetic regulators stabilize cell state transitions according to evolutionary constraint. **a.** Euclidean distances from parental to child cell state pairs, according to the cell lineage tree in Fig. 6f, for all the conditions we used for this study. The x-axis is sorted by additional datasets, namely Dnmt1/Tasor double mutant and Tasor time course mutants, followed by the severity of developmental arrest shown in Fig. 1e. In general, regulators with more severe phenotypes show lower pairwise transcriptional relationships between developmentally connected cell states, consistent with a model where epigenetic regulators support differentiation by consolidating distinct transcriptional states. **b.** Evolutionary traits, i.e., dN/dS scores and gene age distribution, of the marker genes of each GEP.

